# A novel genetic fluorescent reporter to visualize mitochondrial nucleoids

**DOI:** 10.1101/2023.10.23.563667

**Authors:** Joshua O. David, Jingti Deng, Mashiat Zaman, Lucy Swift, Fatemeh Shahhosseini, Abhishek Sharma, Daniela Bureik, Francesco Padovani, Alissa Benedikt, Amit Jaiswal, Armaan Mohan, Craig Brideau, Savraj Grewal, Kurt M. Schmoller, Pina Colarusso, Timothy E. Shutt

## Abstract

Mitochondria contain their own genome (mtDNA), which is present in hundreds of copies per cell and organized into nucleoid structures that are distributed throughout the dynamic mitochondrial network. Beyond encoding essential protein subunits for oxidative phosphorylation, mtDNA can also serve as a signalling molecule when it is present into the cytosol. Despite the importance of this genome, there are still many unknowns with respect to its regulation. To study mtDNA dynamics in living cells, we have developed a genetic fluorescent reporter, mt-HI-NESS, which is based on the HI-NESS reporter that uses the bacterial H-NS DNA binding domain. Here, we describe how this reporter can be used to image mtDNA nucleoids for live cell imaging without affecting the replication or expression of the mtDNA. In addition to demonstrating the adaptability of the mt-HI-NESS reporter for multiple fluorescent proteins, we also emphasize important factors to consider during the optimization and application of this reporter.

## Introduction

Mitochondria are endosymbiotic organelles that harbour their own genome (mtDNA), evidence of their bacterial origins. In humans, mtDNA is ∼16.5 kbp and is located in the mitochondrial matrix; it encodes 2 ribosomal RNAs, 22 tRNAs, and 13 proteins that are essential components of oxidative phosphorylation complexes^1^. Unlike the nuclear genome, mtDNA is circular and is present in hundreds of copies per cell. Although the mtDNA copy number varies in different cell types, the mechanisms regulating this variability are not fully understood. The packaging, maintenance, and expression of mtDNA is carried out by a dedicated set of proteins that are encoded in the nucleus, translated in the cytosol, and imported into mitochondria. The mtDNA is packaged into nucleoid structures, in large part by the mtDNA binding protein TFAM^2^, which also acts as a transcription factor^3^.

In mammals, nucleoids typically contain ∼1.4 mtDNA molecules^4^ and are distributed evenly throughout the dynamic mitochondrial network. Impairments in mitochondrial fission and fusion can impact both mtDNA copy number, and the distribution of nucleoid^5–8^. Fission events are implicated in licensing mtDNA replication^9,10^, while fusion is thought to be important for an even distribution of the machinery required for mtDNA replication and maintenance^11^. Meanwhile, impairments to either fission^12–15^ or fusion^11,16,17^ can also lead to clustering of nucleoids into larger aggregates. Further, reductions in mitochondrial fusion can also lead to individual mitochondria that lack nucleoids entirely^18^. More recently, it is recognized that mtDNA can be released into the cytosol, where it can act as a signaling molecule that triggers innate immune inflammatory pathways^19^. In addition, mtDNA can also be released into extracellular vesicles^20^. However, the exact mechanism mediating cytosolic and extracellular mtDNA release are not fully understood, in part due to technical challenges to visualize mtDNA dynamics via microscopy.

The direct visualization of mtDNA distribution and dynamics in living cells is essential for understanding these processes. Several tools are available to image mtDNA nucleoids, including immunofluorescence, DNA dyes, and fluorescent protein tags^21^. However, these tools are limited in their applicability for live-cell imaging. For example, immunofluorescence, against either DNA or nucleoid proteins requires fixation, and can thus only provide a snapshot of mtDNA dynamics. Meanwhile, DNA-specific dyes, such as PicoGreen^22,23^ or SYBR Gold^24,25^ can label mtDNA for live cell imaging. However, these dyes are not specific for mtDNA as they will bind any DNA in the cell, and can sometimes bind RNA. Moreover, due to their DNA intercalating activity, these dyes can impair mtDNA replication and transcription^21^. Finally, the binding affinity of these dyes can be impacted by the degree of mtDNA packaging, which may prevent intercalation^26^. On the other hand, when considering genetic constructs to label nucleoids, selectivity is a barrier for using fluorescently tagged proteins. Although several nucleoid proteins have been identified, they are not all exclusive to nucleoids (*e*.*g*., ATAD3A)^26^ or do not label all nucleoids (*e*.*g*., TFAM^27^, TWINKLE^28^, mtSSB^28^). Moreover, overexpression of nucleoid proteins can impact mtDNA regulation (*e*.*g*., TFAM^29,30^, TWINKLE^31,32^, M19^33^). Thus, using dyes or overexpressing nucleoid proteins tagged with a fluorescent protein can have adverse effects on mtDNA that preclude observing nucleoids in their natural state.

To avoid nucleoid protein overexpression artifacts, it may be possible to introduce a fluorescent protein tag into a particular gene via CRISPR technology, enabling endogenous expression of the tagged nucleoid protein. However, this complex approach is limited in its applicability. It is also possible that adding a tag could impact the function of the tagged nucleoid protein. For example, many fluorescent proteins tend to oligomerize^34^, which can introduce artefacts^35^. Meanwhile, tagging TFAM, the most well-characterized mtDNA nucleoid protein may be problematic, as it is targeted to mitochondria via an N-terminal mitochondrial targeting sequence (MTS), as are many proteins that localize to the mitochondrial matrix. Finally, TFAM also has a critical C-terminal region required for its transcription-stimulating activity^36^. In addition, overexpressing TFAM impacts mtDNA copy number and can impair mtDNA gene expression^37^. Thus, better tools are required to study mtDNA dynamics.

With the goal of developing a non-invasive genetic fluorescent marker to visualize mtDNA nucleoids for live-cell imaging, we have adapted the recently described HI-NESS genetic reporter^38^, which fuses a fluorescent protein to the bacterial H-NS DNA binding domain, and labels nuclear DNA without any reported adverse effects^39^. Notably, H-NS-like proteins have a preference for AT-rich sequences^40,41^, which should make them ideal for the AT-rich human mtDNA. Despite the bacterial history of mitochondria, the core machinery for their transcription and replication does not resemble typical bacterial machinery, but instead is derived from T-odd phages^42^, and has acquired additional unique regulatory factors^43–45^. Therefore, a HI-NESS reporter is not expected to impact mitochondrial DNA replication or transcription by interacting with this mitochondrial machinery. Thus, as a novel approach to monitor mtDNA nucleoid dynamics, we designed and tested several modified versions of the HI-NESS reporter targeted to mitochondria.

## Results

### Construct Design - mt-Kaede-HI-NESS

As with the original HI-NESS construct^38^, we started with the DNA binding domain of the *E. coli* H-NS protein (amino acids 80-137; GenBank: EIS4692233.1), which we fused to the C-terminus of a fluorescent protein. We initially observed the best results using the photoconvertible fluorescent protein Kaede, derived from *Trachyphyllia geoffroyi*, which irreversibly switches its fluorescence emission from green to red upon activation with UV or violet light^55^. Next, to target the reporter to mitochondria, we added a double Cox8 MTS^46^ to the N-terminus of the construct, as this MTS is known to target mitochondrial import of fluorescent proteins^47,46^. The MTS is followed by an unstructured linker sequence (GDPPVAT) used previously for fluorescent proteins localized to the mitochondrial matrix^48^. Finally, to facilitate immunodetection of the reporter, we added an HA epitope tag (YPYDVPDYA) to the C-terminus of the H-NS DNA binding domain, which has previously been shown not to interfere with DNA binding of the H-NS protein^49^.

The organization of this initial open reading frame, which we refer to as mt-Kaede-HI-NESS, is shown in (Fig 1A), along with a predicted 3D model of the protein binding DNA generated using AlphaFold 2^50^ (Fig 1B). The nucleotide and protein sequences for this reporter can be found in the supplemental material (Fig S1). The full-length mt-Kaede-HI-NESS protein has a predicted size of 40.2 kDa. However, following the removal of the MTS, which is cleaved and processed by a series of mitochondrial proteases^51^, the mature form is predicted to be 34.2 kDa (TargetP 2.0)^52^ to 35.6 kDa (MitoFates)^53^.

**Figure 1.**
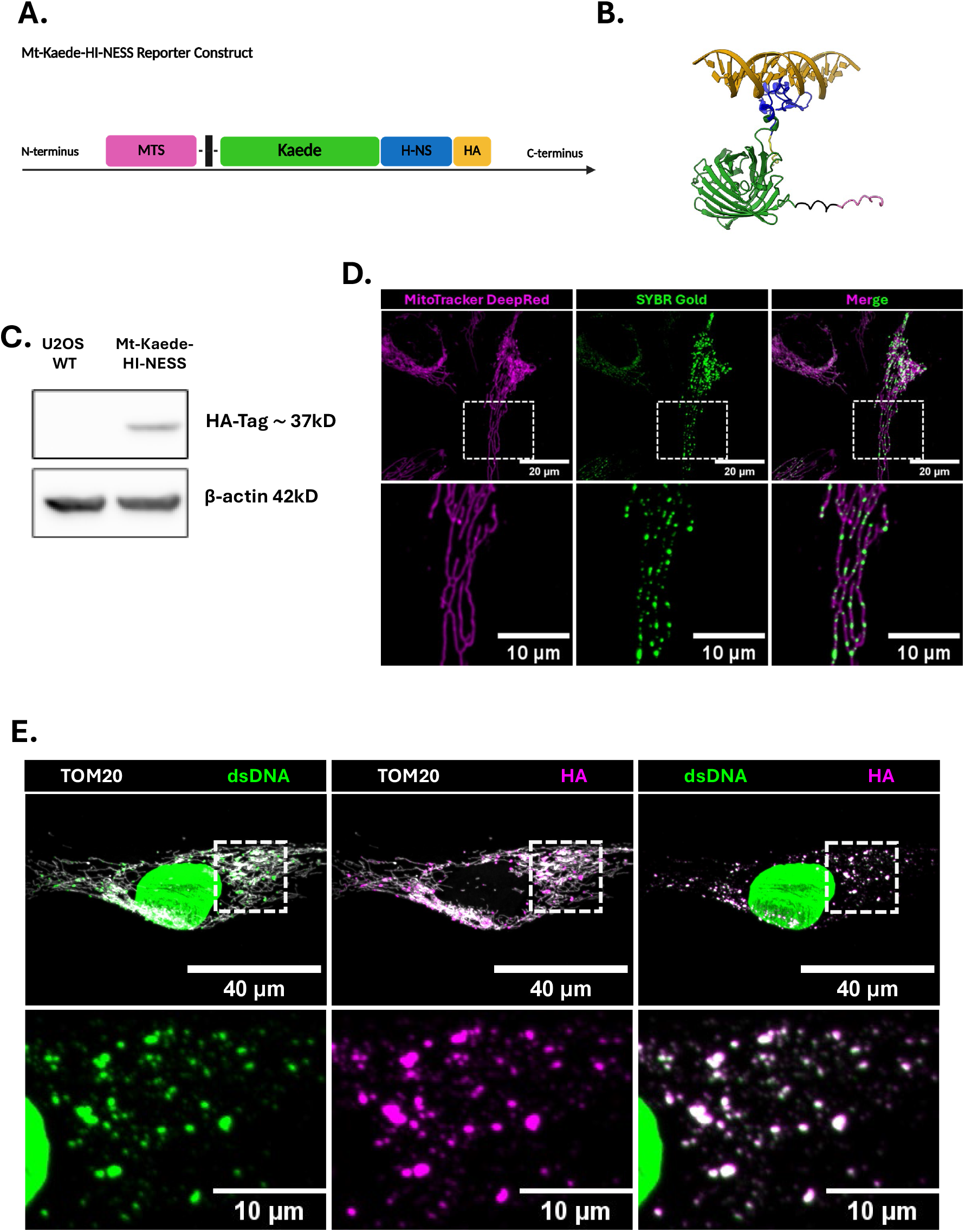
Introduction to Mt-Kaede-HI-NESS reporter. **(A)** Schematic showing the domain organization of mt-Kaede-HI-NESS protein; Mitochondrial Targeting Sequence (MTS) on the N-terminus, a linker peptide, the fluorescent protein (Kaede), a bacterial nucleoid associated protein (H-NS), and a HA-Tag on the C-terminal. **(B)** Predicted 3D structure of the mature protein of Mt-Kaede-HI-NESS generated with AlphaFold 2 bound to mtDNA as predicted via HADDOCK. (C) Colours indicate mitochondrial targeting sequence (pink), linker (black), Kaede (green), H-NS DNA binding domain (blue), HA tag (yellow) and DNA (gold). **(C)** Western blot confirming reporter expression; Whole-cell lysate from untransduced control and Mt-Kaede-HI-NESS transduced cells were probed with anti-HA, revealing a band at the expected molecular weight of the HA-tagged fusion exclusively in the transduced cell. (Vinculin shown as loading control). **(D)** Confocal live image of Mt-Kaede-HI-NESS (green) expressing cells following stable transduction. Mitochondria network labeled with MitoTracker DeepRed (Magenta). **(E)** Confocal immunofluorescence images of Mt-Kaede-HI-NESS–expressing cells showing colocalization of Mt-Kaede-HI-NESS (anti-HA, magenta) with dsDNA (green) within the mitochondrial network labeled with anti-TOM20 (gray). **(F)** Pearson’s correlation coefficient was calculated to quantify colocalization between the mt-Kaede-HI-NESS reporter HA-tag and dsDNA immunostaining in fixed cells. Analysis was performed on 25 randomly selected cells.

VectorBuilder (Chicago, IL, USA) was commissioned to synthesize the mt-Kaede-HI-NESS open reading frame and insert it into a lentiviral mammalian gene expression vector. The lentivirus vector expresses the mt-Kaede-HI-NESS open reading frame under an hPGK promoter and contains puromycin and ampicillin resistance cassettes for mammalian and bacterial selection, respectively. The vector ID for mt-Kaede-HI-NESS is VB230322-1526htn, which can be used to retrieve detailed information about the vector of vectorbuilder.com. A vector map (Fig S2) and the DNA sequence (Fig S3) are provided in the supplemental material.

### Characterization of mt-Kaede-HI-NESS

We generated lentivirus that was used to transduce U2OS cells for stable expression of mt-Tag-RFP-HI-NESS. Fluorescence-Activated Cell Sorting (FACS) was used to select cells expressing Kaede. Expression of the mt-Kaede-HI-NESS reporter was confirmed both by immunoblotting with anti-HA and live-cell confocal microscopy (Fig 1C-D). Although the expression of mt-Kaede-HI-NESS was variable among different cells, we consistently observed Kaede signal localizing to puncta within the mitochondrial network during live cell imaging (Fig 1D). To confirm that mt-Kaede-Hi-NESS specifically labels mitochondrial nucleoids, confocal imaging of fixed cells showed colocalization between the Kaede-HA signal and dsDNA within the mitochondrial network (Fig 1E). This initial result provided strong evidence that mt-Kaede-HI-NESS enables reliable visualization of mtDNA nucleoids in both live and fixed cells.

Due to the variable expression levels varied among cells in the initial transduced population, FACS was used to isolate single cells, and three clones were selected to represent low-, mid-, and high-expression as confirmed by immunoblotting analysis and fluorescence intensity measured by flow cytometry (Fig 2A-C). These three clones were used for subsequent analyses to examine whether expression levels of the mt-Kaede-HI-NESS reporter affected mtDNA maintenance or mitochondrial function.

**Figure 2.**
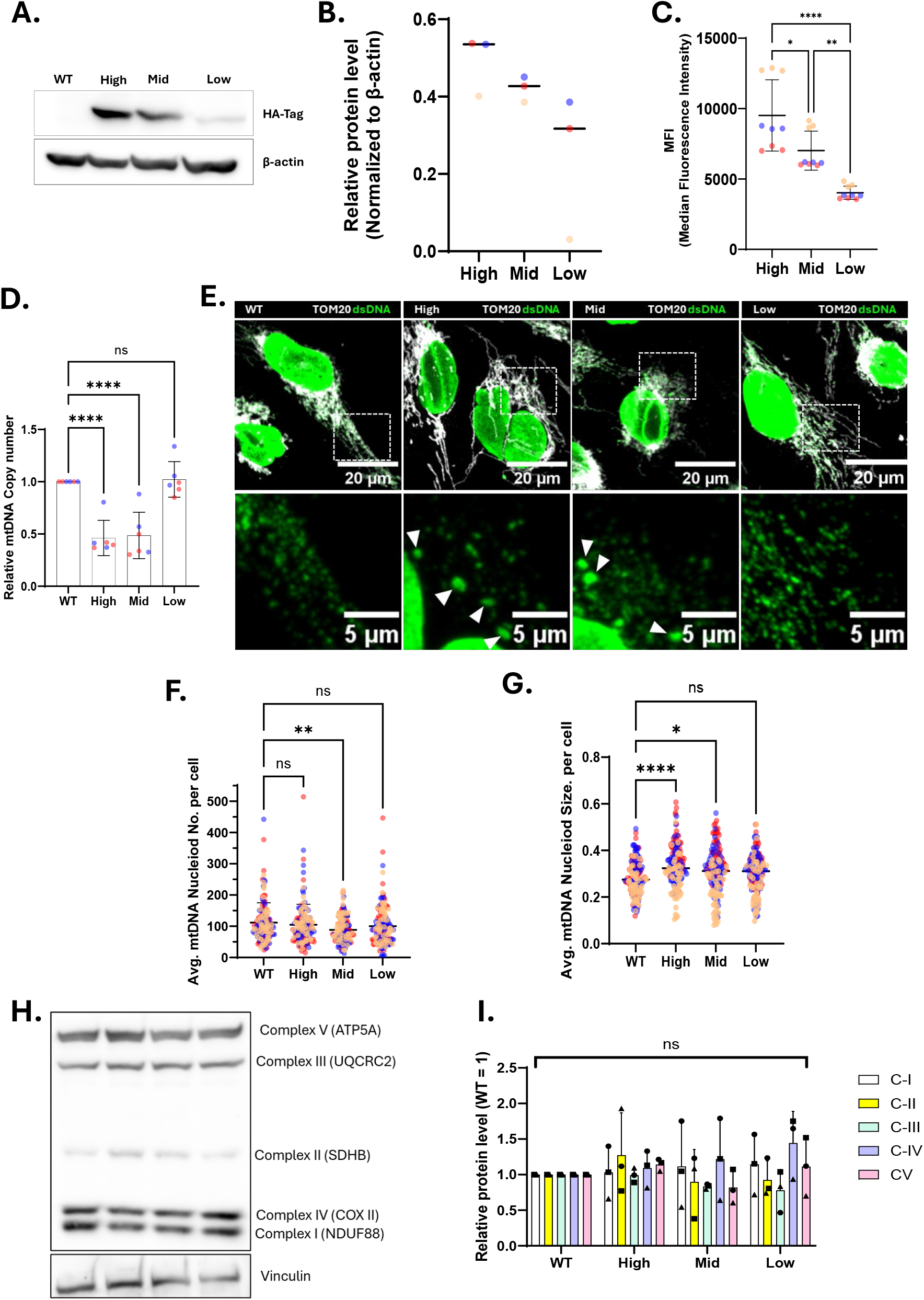
Classification of mt-Kaede-HI-NESS reporter expression and mitochondrial function: **(A)** Whole-cell lysates from high-, mid-, and low–mt-Kaede-Hi-NESS–expressing cells and untransduced controls were analyzed by Western blot using anti-HA. Equal amounts of total protein were loaded, and β-actin served as a loading control. **(B)** Densitometric quantification of HA signal immunoblotting as in panel G, normalized to β-actin. **(C)** Reporter expression levels were assessed by flow cytometry median fluorescence intensity (MFI). One-way ANOVA showed significant differences among clones. Post hoc Tukey’s multiple comparisons revealed that high-, mid-, and low-expressing cells each differed significantly from one another (High vs. Mid: p = 0.0119; High vs. Low 13: p < 0.0001; Mid vs. Low: p = 0.0026). Data represent mean ± SD. **(D)** mtDNA copy number quantified by qPCR (normalized to a nuclear reference gene; ND1 to 18S). Ordinary One-way ANOVA revealed significant differences among groups. Post Hoc Dunnett’s multiple comparisons test showed that both High- and Mid-expressing clones had significantly reduced mtDNA copy number compared with control (p < 0.0001), whereas Low expressing cells was not significantly different (p=0.9911), Data represent mean ± SD (n=6 per group). **(E)** Representative fixed-cell images of high-, mid-, and low-mt-Kaede-Hi-NESS– expressing cells and untransduced controls showing nucleoid morphology. TOM20 marks mitochondria (gray), and dsDNA marks nucleoids (green). Arrows indicate large nucleoid clumps. **(F)** mtDNA nucleoid number per cell was quantified from images acquired from confocal microscopy. One-way ANOVA and Post hoc Dunnett’s test showed that Mid-expressing cells had significantly fewer nucleoids compared to control (p = 0.0021), whereas High- and Low-expressors did not differ significantly from control (p = 0.6623 and p = 0.2455, respectively). Data represent mean ± SD with ≥145 cells analyzed per group. Colour of data point indicate different biological replicates total = 3. **(G)** mtDNA nucleoid number per cell was quantified from images acquired from confocal microscopy. One-way ANOVA and Post hoc Dunnett’s comparisons indicated that High-(p < 0.0001) and Mid-expressors (p = 0.0417) had significantly larger nucleoids compared to control, whereas low-expressors did not differ significantly (p = 0.2991). Data represent mean ± SD with ≥145 cells analyzed per group. Colour of data point indicates different biological replicates total = 3. **(H)** Representative Western blot showing complexes I–V in control (WT) and reporter-expressing cells; Vinculin served as a loading control. **(I)** Quantification of OXPHOS complexes normalized to loading control and expressed relative to untransduced control (WT). Two-way ANOVA showed no significant difference. Dunnett’s multiple comparisons test confirmed that none of the clones differed significantly from WT for any individual complex (all adjusted p > 0.25). Data are mean ± SD, n = 3 biological replicates.

To evaluate mtDNA maintenance or nucleoid organization, we first quantified mtDNA copy number by qPCR. Both the mid- and high-expressing clones showed a reduction in mtDNA copy number relative to untransduced control cells, whereas the low-expressing clone remained comparable to the control (Fig 2D). We next examined mtDNA nucleoid morphology in each clone (Fig 2E). The average number of nucleoids per cell was not significantly altered, except for the mid-expressing clone, which exhibited an approximately 20% decrease compared with untransduced control cells (Fig 2F). In contrast, analysis of the average nucleoid size revealed approximately 18% and 9% increase in the high- and mid-expressing clones, respectively, while the low-expressing clone remained comparable in size to the untransduced control cells (Fig 2G). This increase in average size is likely due to the appearance of some larger nucleoids in these cells (indicated by arrows in Fig 2E), which may reflect nucleoid clustering, likely driven by overexpression-related artifacts. To determine whether these changes affect or cause alterations to mitochondrial respiratory components, we examined OXPHOS complex abundance by immunoblotting. Average protein levels of complexes I–V were comparable across all clones and the untransduced control (Fig 2H-I).

Together, these results indicate higher expression of the mt-Kaede-HI-NESS reporter can impact mtDNA copy number and nucleoid sizes, but does not appear to substantially affect the steady-state levels or function of mitochondrial respiratory proteins. Nonetheless, it is possible to select for clones with lower expression levels of the reporter that do not impact mtDNA.

### Applications

Building on the successful development of a novel reporter that specifically labels mtDNA in both live and fixed cells, we next tested the ability of the mt-Kaede-Hi-NESS reporter to visualize mitochondrial nucleoid dynamics over time. While viral transduction provides stable cell lines that can be repeatedly studied, selection can be tedious and time-consuming. Thus, we compared mtDNA nucleoids in our stably transduced lentiviral mt-Kaede-HI-NESS cells with U2OS cells transiently transfected with the mt-Kaede-HI-NESS reporter (Fig 3A). While there are some differences in the distribution of mtDNA nucleoid sizes and numbers between stable and transient mt-Kaede-HI-NESS expression (Fig 3B-C), it is unclear whether the larger distribution of sizes is due to different expression levels or the stress of the transfection. We next wanted to test the ability of the mt-Kaede-HI-NESS reporter to visualize changes in mitochondrial nucleoids and to track dynamic events such as cytosolic and extracellular mtDNA release.

**Figure 3.**
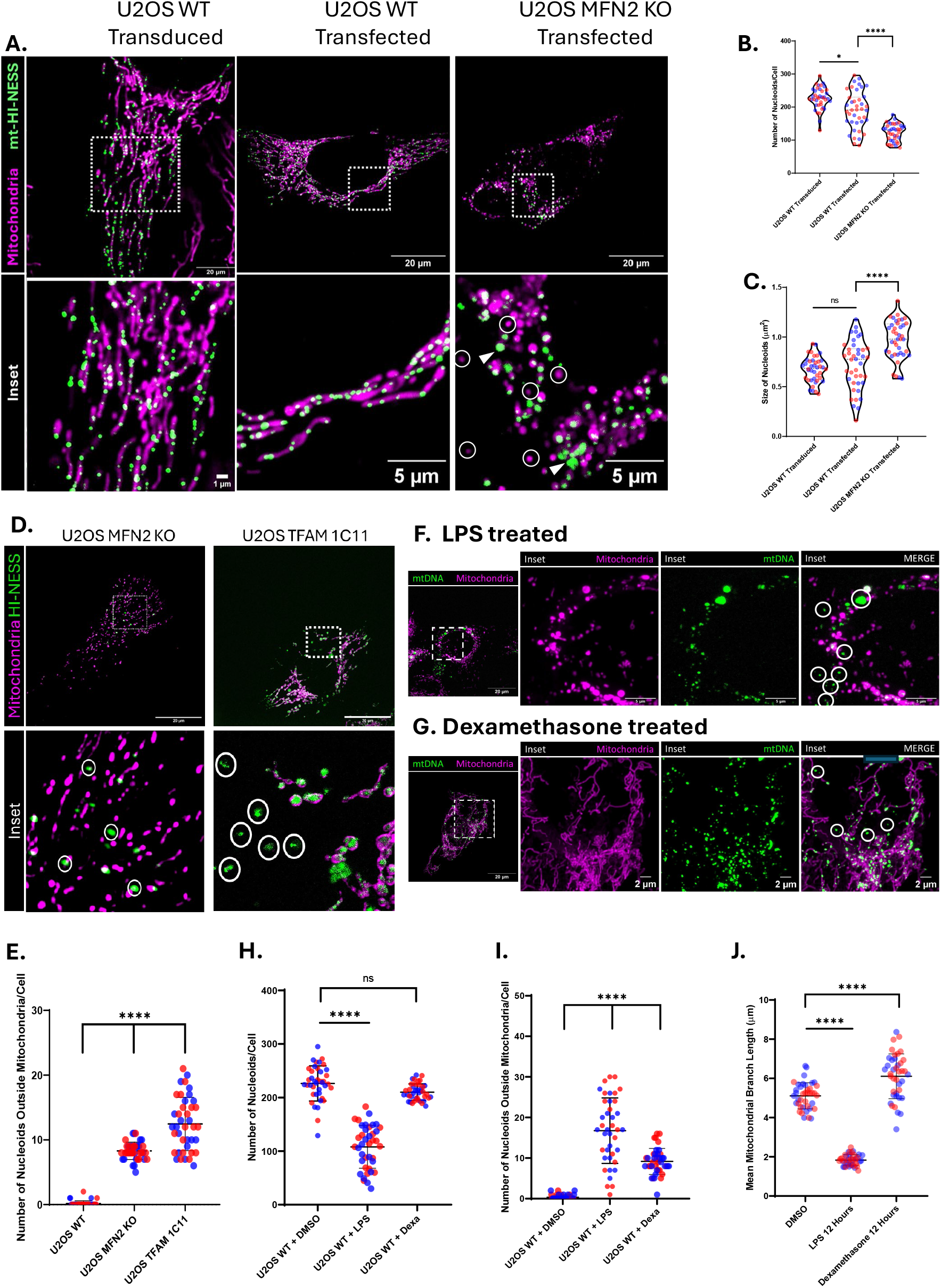
Using Mt-Kaede-HI-NESS to Visualize mtDNA Release in Models of Mitochondrial Dysfunction.. **(A)** Representative confocal images showing mitochondria (MitoTracker DeepRed) and mtDNA (mt-Kaede-HI-NESS) in transduced (WT U2OS) and transfected cell models (WT U2OS and U2OS MFN2 KO). **(B-C)** Quantitative analyses of **(B)** number and **(C)** size of mtDNA nucleoids in transfected vs transduced cell models (U2OS WT/MFN2 KO). Colours indicate biological replicates, violins indicate median and interquartile range, multiple unpaired t-tests. Stats: ns P > 0.05; *P ≤ 0.05; ****P ≤ 0.0001. **(D)** Representative confocal images showing mitochondria (MitoTracker Deep Red) and mtDNA (mt-Kaede-HI-NESS) in U2OS MFN2 knockout cells and U2OS TFAM 1C11 cells, circles indicate mtDNA not localized to the mitochondrial network. **(E)** Quantitative analysis of number of mtDNA puncta not localized to the mitochondrial network, colours indicate biological replicates, lines indicate mean +/-SD, two-way ANOVA. **(F-G)** Representative confocal images showing drug-induced extra-mitochondrial mtDNA, as shown by addition of **(F)** LPS and **(G)** Dexamethasone. **(H-G)** Quantitative analyses of drug-based models of mtDNA release, showing **(H)** number of nucleoids, **(I)** number of extra-mitochondrial nucleoids and **(J)** mean mitochondrial branch length. Colours indicate biological replicates, lines indicate mean +/-SD, two-way ANOVA. Stats: ns P > 0.05; ****P ≤ 0.0001.

#### Nucleoid dynamics and morphology

To examine if the mt-Kaede-HI-NESS reporter can be used to see changes in nucleoid dynamics, we used U2OS MFN2^-/-^ cells^54^, as loss of the mitochondrial fusion protein MFN2 leads to fragmented mitochondrial networks and altered mtDNA characteristics and distribution^11,55^. As expected, following transient transfection of mt-Kaede-HI-NESS into U2OS control or U2OS MFN2^-/-^, cells lacking MFN2 had nucleoids that were larger (shown with arrow) and fewer in number compared to control, as well as a subset of mitochondria fragments lacking nucleoids (circled) (Fig 3A-C).

#### Cytosolic mtDNA release

To see if the mt-Kaede-HI-NESS reporter could be used to visualize mtDNA that had escaped into the cytosol, we first used two established genetic models of mtDNA release, MFN2^-/-56^ and TFAM^-/+ 57,58^, both in U2OS cells. In both cases, following transient transfection, we were able to visualize and quantify cytosolic mtDNA using the mt-Kaede-HI-NESS reporter (Fig 3D-E). Next, we treated stably transduced mt-Kaede-HI-NESS U2OS cells with LPS (20ug/ml for 12 hours) or dexamethasone (200nM for 18 hours), two treatments that induce cytosolic mtDNA release ^59,60^. Again, both treatments lead to an increase in cytosolic mtDNA puncta (Fig 3F-J). Together these finding show that the mt-Kaede-HI-NESS reporter can be used to image cytosolic mtDNA release under a variety of conditions.

#### Extracellular mtDNA release

To examine the release of mtDNA into extracellular vesicles (EVs) we used our stably transduced mt-Kaede-HI-NESS cells and looked both basally, and following overnight treatment with LPS (2 μg/ml) in FBS-free medium to stimulate the formation of EVs^61^ (Fig 4). Following immunocapture of EVs with a CD13 antibody^61^ (Fig 4A,B), we were able to visualize the mt-Kaede-HI-NESS protein in a subset of EVs by confocal imaging, and quantify that LPS treatment increased the size of EVs, and the percentage of Kaede positive EVs (Fig 4C-H). Additionally, immunoblotting of the cellular and extracellular material following differential centrifugation showed the presence of the mt-Kaede-HI-NESS protein (probed with anti-HA) in the extracellular fraction, which was enriched upon LPS treatment (Fig 4J). Finally, quantitative PCR also confirmed the presence of mtDNA in the extracellular fraction and enrichment upon LPS treatment (Fig 4I). Collectively, this work demonstrates that the mt-Kaede-HI-NESS reporter can be used to monitor the release of mtDNA via EVs.

**Figure 4.**
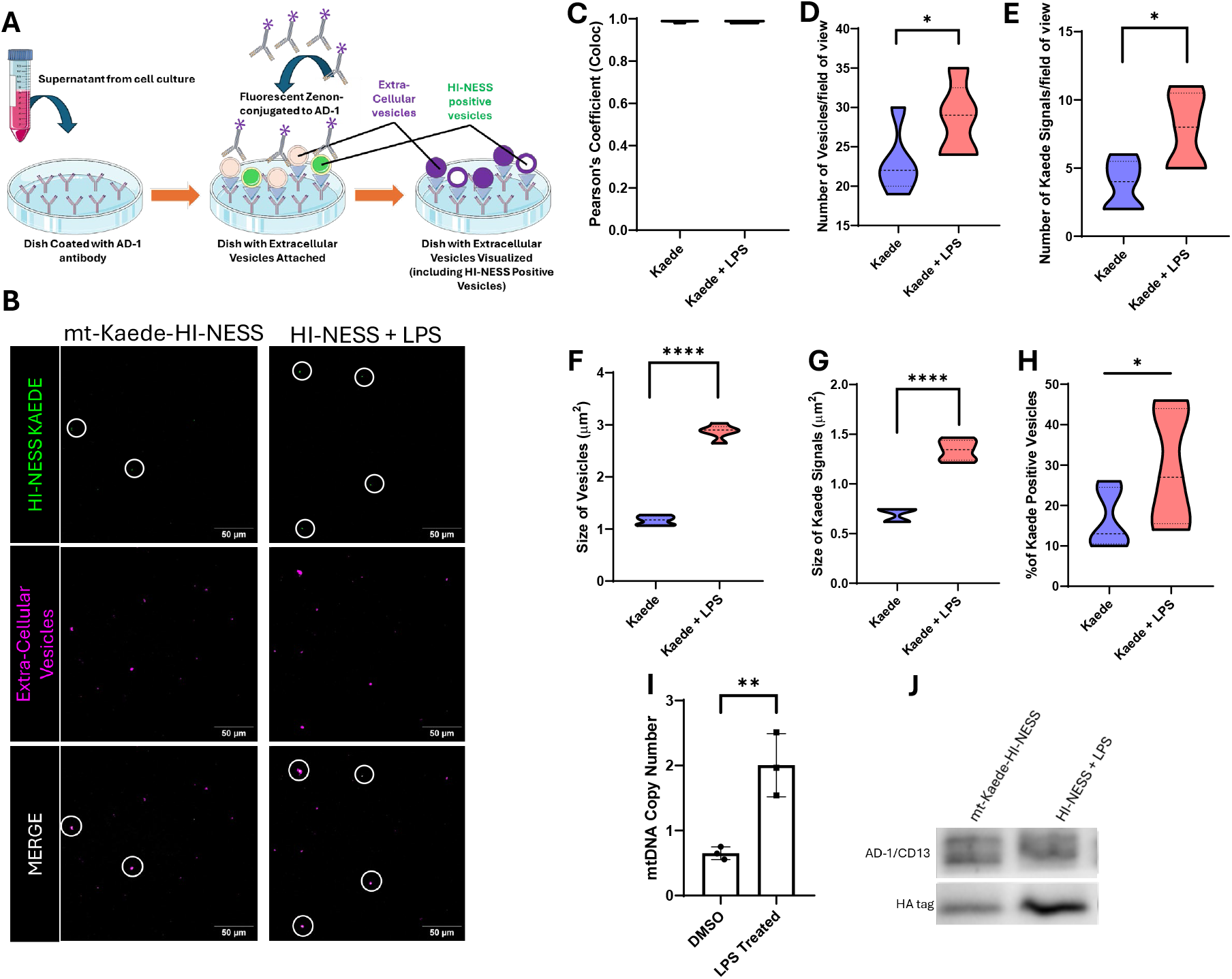
Extracellular mtDNA reported using mt-Kaede-HI-NESS. **A)** Schematic of the cell-free mtDNA detection assay. **B)** Representative confocal images showing mtDNA (mt-Kaede-HI- NESS) and extra-cellular vesicles (anti-CD13) in LPS treated and untreated cells, circles indicate presence of mtDNA. **C-H)** Quantification of extra-cellular mtDNA showing **(C)** Pearson’s co-localization coefficient between green and magenta signals, number of **(D)** extra-cellular vesicles and **(E)** extra cellular mtDNA puncta, size of **(F)** extra-cellular vesicles and **(G)** extra cellular mtDNA puncta and **(H)** percentage of extracellular vesicles that contain mtDNA. All violins show median and interquartile range, unpaired t-test. I) Relative mtDNA copy number in the extracellular fraction for LPS treated and untreated cells, bars indicate mean +/-SD, unpaired t-test. **J)** Representative bands from western blot analysis of the extracellular fraction for LPS treated and untreated cells, showing relative protein amounts of CD-13 and HA-tag in LPS treated/untreated cells. Stats: *P ≤ 0.05; **P ≤ 0.01; ****P ≤ 0.0001.

### Expanding the mt-HI-NESS colour palette

Having demonstrated the utility of mt-Kaede-HI-NESS to visualize nucleoids, we next explored whether this reporter could be adapted to other fluorescent proteins to broaden its utility for additional imaging applications. Thus, we generated several mt-Hi-NESS variants by replacing Kaede domain with different fluorescent proteins. To generate a red-emitting mt-HI-NESS, we used TagRFP (derived from *Entacmaea quadricolor*)^62^ (GenBank: ABR08320.1), which is established to function in the environment of the mitochondrial matrix. The adapted mt-TagRFP-Hi-NESS has a predicted full-length size of 40.6 kDa and a processed mitochondrial form of approximately 34–36 kDa (Fig S1).

To enable targeted induction of oxidative damage on mtDNA and to visualize the resulting mtDNA damage responses and dynamics, an additional reporter variant incorporating KillerRed, a fluorescent protein derived from *Anthoathecata*, was also generated. KillerRed is a genetically encoded photosensitizer that produces reactive oxygen species (ROS) upon light irradiation and induces cell damage when excited with green light (540–580 nm)^63^; GenBank: AAY40168.1). The mt-KillerRed-Hi-NESS protein has a predicted full-length molecular weight of ∼41.0 kDa and a mature mitochondrial form of ∼36.3 kDa (Fig S1).

To assess nucleoid labeling specificity for mt-TagRFP-Hi-NESS and mt-KillerRed-Hi-NESS reporters, both transiently transfected and stably transduced cells were analyzed. Unexpectedly, mt-TagRFP-Hi-NESS (Fig 5A) and mt-KillerRed-Hi-NESS (Fig 5B) reporters exhibited predominantly diffuse mitochondrial fluorescence with little to no discrete punctate labeling of nucleoids. In contrast to the robust punctate labeling observed with mt-Kaede-HI-NESS, this difference prompted investigation into the underlying cause of the diffuse mitochondrial signal. As the only difference between these reporters was the fluorescent protein backbone, we suspected an inherent fluorescent protein–specific property was responsible for the difference in mtDNA specific labeling. Notably, differences in oligomerization behavior of fluorescent represent a well-recognized variable^64^ that could impact mtDNA specificity of the mt-HI-NESS reporters through cooperative binding. In this regard, Kaede is a well-characterized tetrameric fluorescent protein^65^. In contrast, TagRFP was engineered to minimize oligomerization and is generally described as monomeric, although weak self-association has been reported in some contexts^62,66^. Moreover, KillerRed, is commonly described as monomeric or dimeric in cellular contexts^63,66^.

**Figure 5.**
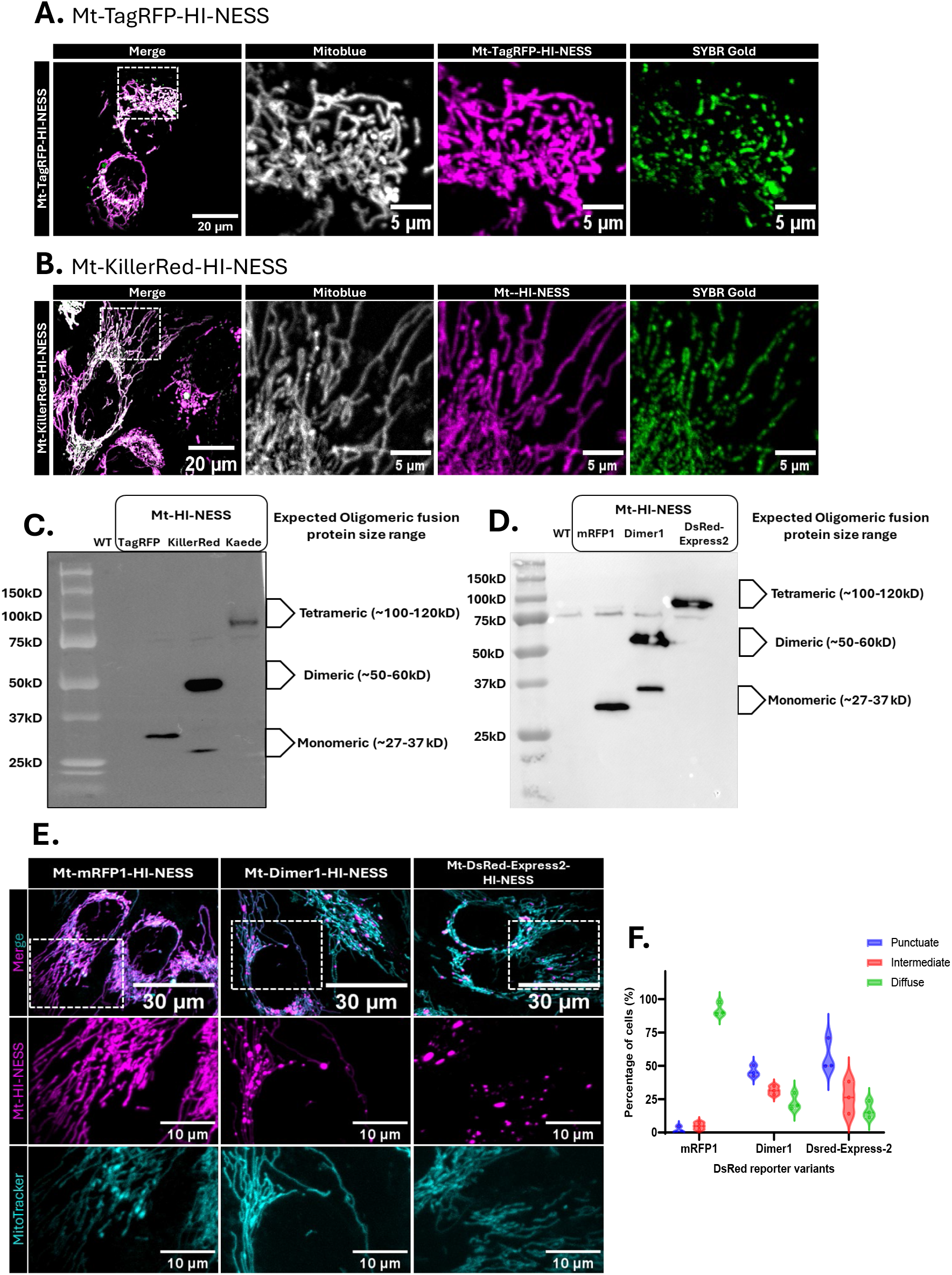
Fluorescent protein oligomerization influences reporter nucleoid-binding specificity. **(A)** Confocal live cell image of Mt-TagRFP-HI-NESS expressing cells (Magenta), mitochondria network labeled with Mitoblue (gray), and mtDNA stained with SYBR Gold (green). **(B)** Confocal live cell image of Mt-KillerRed-HI-NESS expressing cells (Magenta), mitochondria network labeled with Mitoblue (gray), and mtDNA stained with SYBR Gold (green). **(C)** Native (non-denaturing) PAGE Western blot showing the oligomerization states of TagRFP-, KillerRed-, and Kaede-based reporter constructs, detected by anti-HA immunoblotting. **(D)** Native (non-denaturing) PAGE Western blot showing the oligomerization states of mRFP1-, Dimer1-, and DsRed-based reporter constructs, detected by anti-HA immunoblotting. **(E)** Representative live cell images showing the DsRED variants-based reporters (mRFP1, Dimer1 and DsRed-Express2) (Mitochondria; Cyan, DsRED variant-HI-NESS: Magenta). **(F)** Visual scoring of reporter localization phenotypes. Cells expressing DsRed-variant–based reporters (mRFP1, Dimer1, and DsRed-Express2) were visually classified as exhibiting punctate, intermediate, or diffuse localization. The percentage of cells in each category was calculated based on the total number of cells analyzed per reporter (mRFP1, n = 420; Dimer1, n = 371; DsRed-Express2, n = 244).

#### Fluorescent protein oligomerization influences reporter nucleoid-binding specificity

To assess these oligomerization state of our mt-HI-NESS variants, we performed non-denaturing (native) PAGE followed by anti-HA immunoblotting (Fig 5C). As expected, mt-Kaede-Hi-NESS migrated as higher-order complexes consistent with its known tetrameric state. In contract, mt-KillerRed-Hi-NESS predominantly migrated as a dimeric species, but also exhibited a small secondary band in the monomeric range. Finally, mt-TagRFP-Hi-NESS migrated predominantly as a monomer (Fig 5C).

To systematically test the relationship between oligomerization state and reporter behavior, we generated a panel of mt-HI-NESS reporters using fluorescent proteins derived from DsRed, which have minimal amino acid differences (Fig S4) and well-defined oligomeric states: mRFP1 (monomer)^67^, Dimer1 (dimer)^67^, and DsRed-Express2 (tetramer)^68^. The nucleotide sequences, protein sequences, and vector maps for these reporters can be found in the supplemental material (Fig S1-S3).

The oligomeric state of these new reporters was confirmed by native PAGE followed by anti-HA immunoblotting. As expected, mt-mRFP1-Hi-NESS migrated predominantly as a monomer, whereas mt-Dimer1-Hi-NESS predominantly migrated as a dimer with a small additional band near the monomeric range, mirroring the pattern observed for mt-KillerRed-Hi-NESS. Finally, mt-DsRed-Express2-HI-NESS migrated as a higher-order tetrameric complex similar to the mt-Kaede-Hi-NESS reporter (Fig 5D).

Having confirmed the expected oligomeric state of these reporters, we then investigated their relative effectiveness for labelling mtDNA via confocal imaging and visual scoring. Notably, only the tetrameric reporters displayed strong punctate nucleoid localization, whereas monomeric and dimeric variants exhibited diffuse or weak labeling (Fig 5E-F;). This pattern suggests that tetrameric fluorescent proteins may benefit from cooperative interactions that increase effective binding affinity and residence time at nucleoids.

#### Double HI-NESS

As many commonly used fluorescent proteins are monomeric, we expect that they would not be amenable for use in the mt-HI-NESS reporter. To overcome this limitation, we engineered a “double Hi-NESS” construct for the monomeric TagRFP reporter, mt-TagRFP-N/C-HI-NESS, by adding an additional H-NS domain immediately after the K56 cleavage site in the MTS, such that one H-NS domain was at the N-terminus and the second was at the C-terminus (Fig 6A). To avoid repetitive sequences in the open reading frame that might have adverse effects on plasmid and/or viral propagation, we took advantage of the degeneracy in codon usage to change the nucleotide sequence without changing the encoded protein domain. To this end, the codon sequence for the N-terminal H-NS domain was ‘humanized’ by choosing codons more commonly used in the human genome. This mt-TagRFP-N/C-HI-NESS reporter has a predicted full-length size of 47 kDa, and a predicted processed size of 43.3 kDa (MitoFates) or 41 kDa (TargetP 2.0). The vector ID for mt-TagRFP-N/C-HI-NESS is VB230529-1292kdj, which can be used to retrieve detailed information about the vector of vectorbuilder.com. A vector map and the DNA sequence is provided in the supplemental material (Fig S2&3). When the mt-TagRFP-N/C-HI-NESS reporter was stably expressed in U2OS cells, confocal imaging revealed a punctate localization of the reporter within the mitochondrial network (Fig 6B). This marked improvement punctate signal compared to the diffuse Mt-TagRFP-HI-NESS signal (Fig 5A) was confirmed as nucleoid-specific by colocalization of the signal with SYBR Gold.

**Figure 6.**
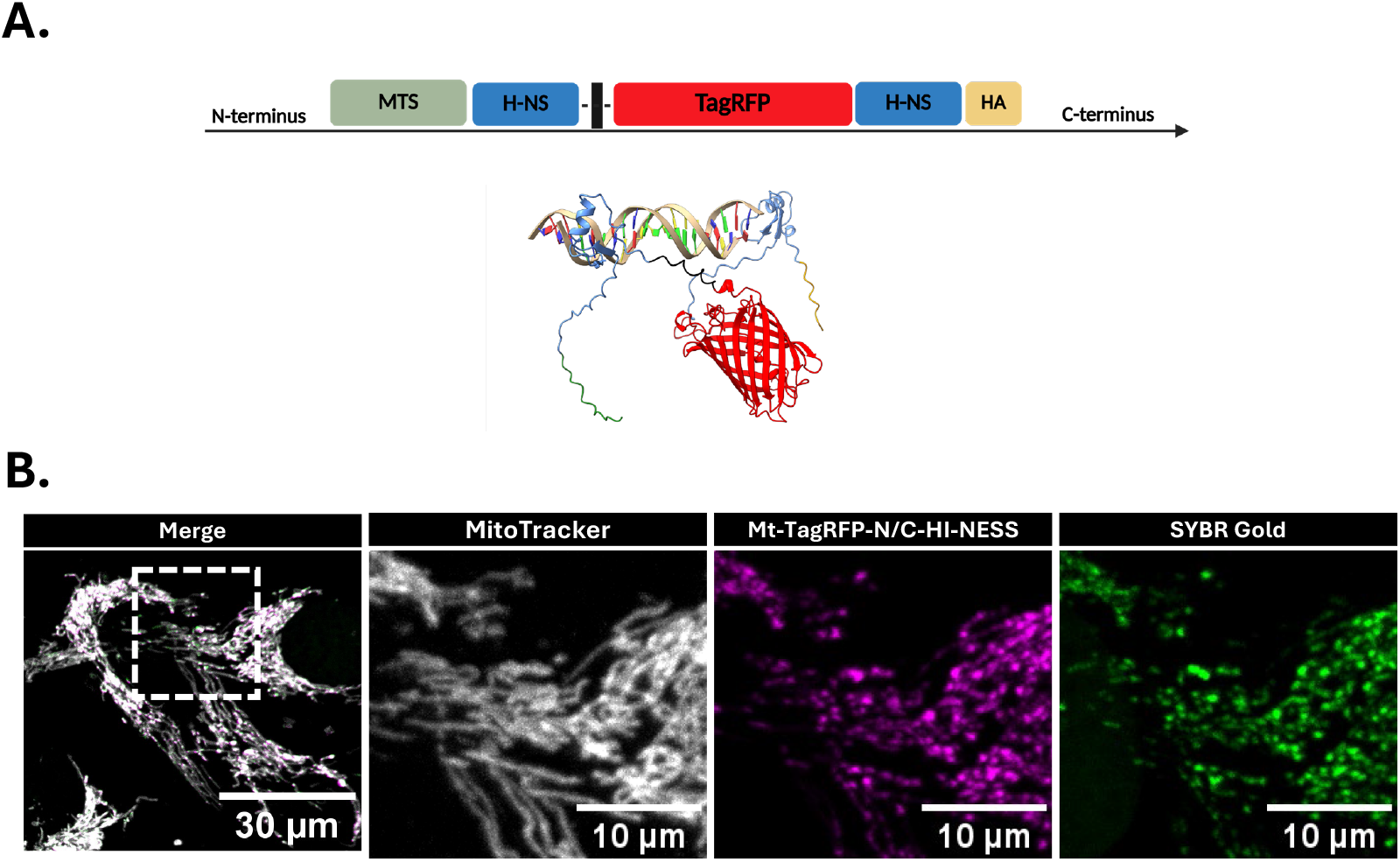
Double HI-NESS: **(A)**Domain schematics and **(B)** Confocal live cell images showing the improved binding capacity of the Double HI-NESS (Mt-TagRFP-N/C-HI-NESS).

### Nucleoid labelling in other systems

We also wanted to test the ability of the mt-Kaede-HI-NESS reporter to label mtDNA in other biological systems. While another report highlights the utility of mt-Kaede-HI-NESS reporter in plants (*Arabidopsis thaliana*) ^69^, here we also show its applicability in both yeast (*Saccharomyces cerevisiae*), and fruit fly (*Drosophila melanogaster*).

#### S. cerevisiae

To test whether the mt-Kaede-HI-NESS reporter can be adapted to visualize mtDNA in budding yeast, we constructed strains in which we integrated mt-Kaede-HI-NESS into the *URA3* locus, N-terminally tagged with a Su9 mitochondrial targeting sequence, and expressed from four different promoters chosen to cover a broad range of expression strengths^70^ (Fig 7A). In addition, each strain expressed Su9-mKate2 as a fluorescent reporter for the mitochondrial network^71^. With qPCR, we found that when expressed from the *RPL25* promoter, i.e. the strongest promoter tested, the mt-Kaede-HI-NESS reporter leads to a significantly reduced mtDNA copy number per cell during exponential growth on SCD medium (synthetic complete medium with 2% glucose as carbon source) (Fig 7B). By contrast, cells expressing the reporter from the *MLC1, CUP1*, or *ACT1* promoters have similar mtDNA copy numbers (Fig 7B), cell volumes (Fig S5A) and bud fractions (Fig S5B) compared to wild-type cells, and do not show any obvious growth defects. We also verified that the strains grow on respiratory media with 2% glycerol and 1% ethanol as carbon sources, in which functional mtDNA is essential.

**Figure 7.**
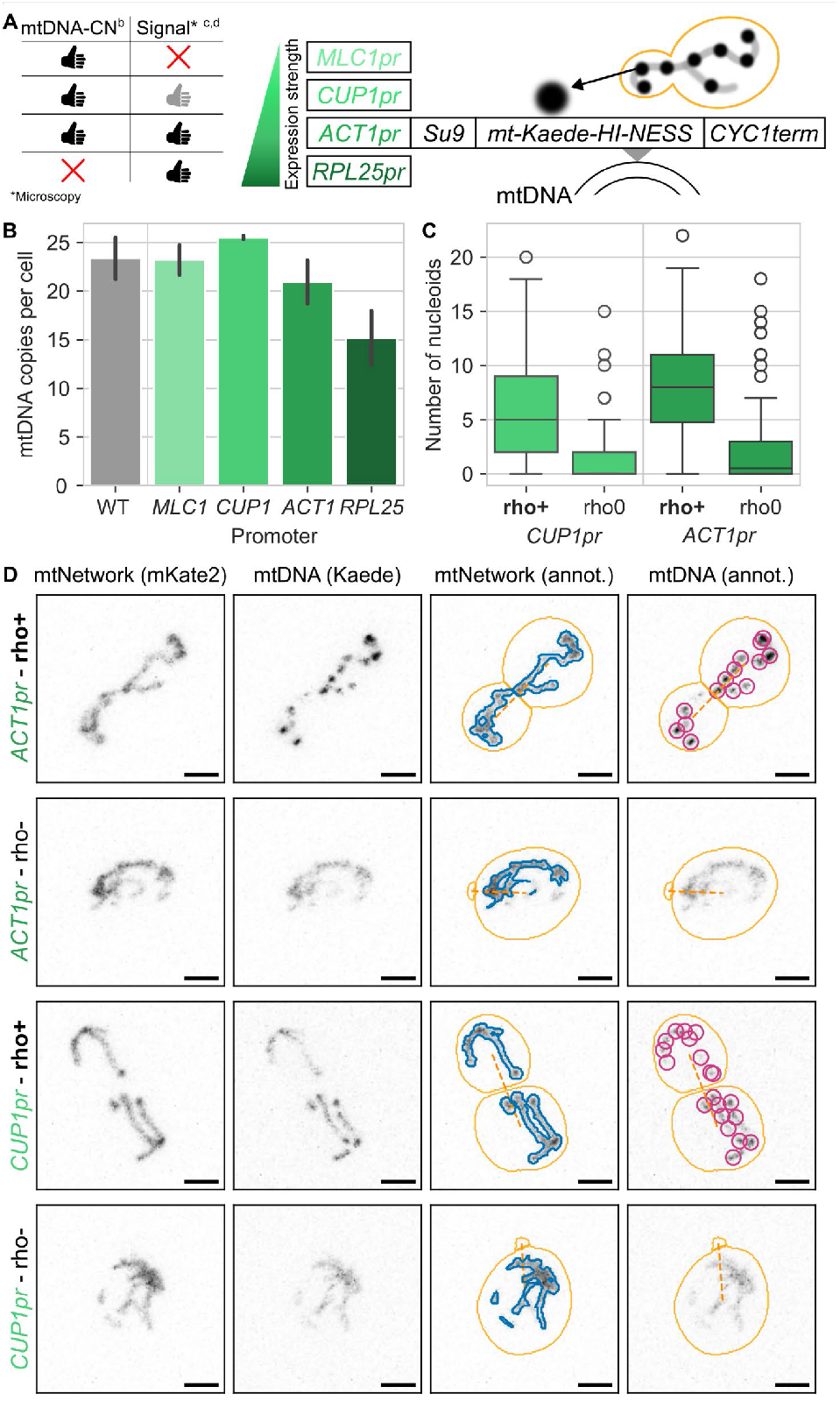
Mt-Kaede-HI-NESS in *S. cerevisiae*. **(A)** Schematic representation of the experimental setup. mt-Kaede-HI-NESS fused to a Su9 mitochondrial targeting sequence was expressed in budding yeast from promoters with varying strength. **(B)** mtDNA copies per cell as determined by qPCR for strains expressing the mt-Kaede-HI-NESS reporter from different promoters as indicated. **(C)** Number of automatically detected high-confidence spots (nucleoids, see methods) for budding yeast cells expressing mt-Kaede-HI-NESS from the *CUP1* or *ACT1* promoters, and corresponding *mip1Δ*a cells. **(D)** Corresponding representative confocal images (maximum projections) as well as automated network segmentation and detected nucleoids. In addition to mt-Kaede-HI-NESS, all strains express mKate2 targeted to the mitochondrial matrix for visualization of the mitochondrial network. Cell outlines and dashed lines connecting buds with their mother cells are shown in yellow. Automatic segmentation of the mitochondrial network is shown in blue. Automatically detected high-confidence spots (nucleoids) are shown in pink. In **C** and **D**, excitation intensities used to image mt-Kaede-HI-NESS were adjusted depending on the promoter to account for different expression levels. Scale bars denote 2 μm.

With live-cell confocal microscopy, we then found that when the reporter was expressed from the *MLC1* promoter, although clearly localized to the mitochondrial network, the signal was very weak and barely above the background (Fig S5C-D). However, when expressed from the *CUP1* or *ACT1* promoters, we observed clear spot-like signal located in the mitochondrial network, as expected for mtDNA containing nucleoids (Fig 7D, Fig. S5C-D). In the case of the *ACT1* promoter, the signal was cleaner compared to the *CUP1* promoter (Fig S5C-D), and required lower excitation intensities (Fig 7D). For all promoters, the mitochondrial network volume is very similar to the control strain without mt-Kaede-HI-NESS reporter (Fig S6E). Next, we used SpotMAX^72^ for automated spot counting, and found that cells contained about 8 resolvable spots in the case of the *ACT1* promoter, which is consistent with what is expected based on similar experiments performed using a LacO-LacI based reporter system, the current gold-standard for mtDNA live-cell imaging in budding yeast^71,73^ (Fig 7C,D). In the case of the *CUP1* promoter, we only detected about 5 clear spots per cell, likely due to the lower signal-to-noise ratio. To further verify that the observed spots correspond to mt-Kaede-HI-NESS bound to mtDNA, we made use of the fact that mtDNA is not essential for budding yeast growth on fermentable media. We deleted the mitochondrial DNA polymerase *MIP1*, resulting in the loss of mtDNA (Fig S5F). Indeed, in *mip1Δ* cells, the mt-Kaede-HI-NESS signal is delocalized throughout the mitochondrial network and does not localize in spots (Fig 7C-D). Taken together, mt-Kaede-HI-NESS can be used as a fluorescent reporter for mtDNA in live budding yeast cells. Expression from the strong *ACT1* promoter provides a high signal-to-noise ratio even at lower excitation intensities well suited for live-cell imaging, without any obvious perturbations of mtDNA and cell function.

#### D. melanogaster

To test the mt-Kaede-HI-NESS reporter in an animal model, we took advantage of *Drosophila* melanogaster and the UAS expression system. We first generated a line carrying the mt-Kaede-HI-NESS open reading frame under control the UAS promoter/enhancer. The UAS-mt-Kaede-HI-NESS reporter transgenic line was crossed to the daughterless-Gal4 line to drive ubiquitous expression of the UAS-mt-Kaede-HI-NESS. Following the isolation of fat body from larvae, we were able to image mt-Kaede-HI-NESS signal, which localized within the mitochondrial network as expected, suggesting the reporter also works in *Drosophila* (Fig 8). However, it should be noted that the larvae did not pupariate, possibly due to a negative impact on normal development caused by high expression of the mt-Kaede-HI-NESS protein. Future efforts will need to be made to regulate the expression of mt-Kaede-HI-NESS, which can be accomplished using different weaker expressing driver lines.

**Figure 8.**
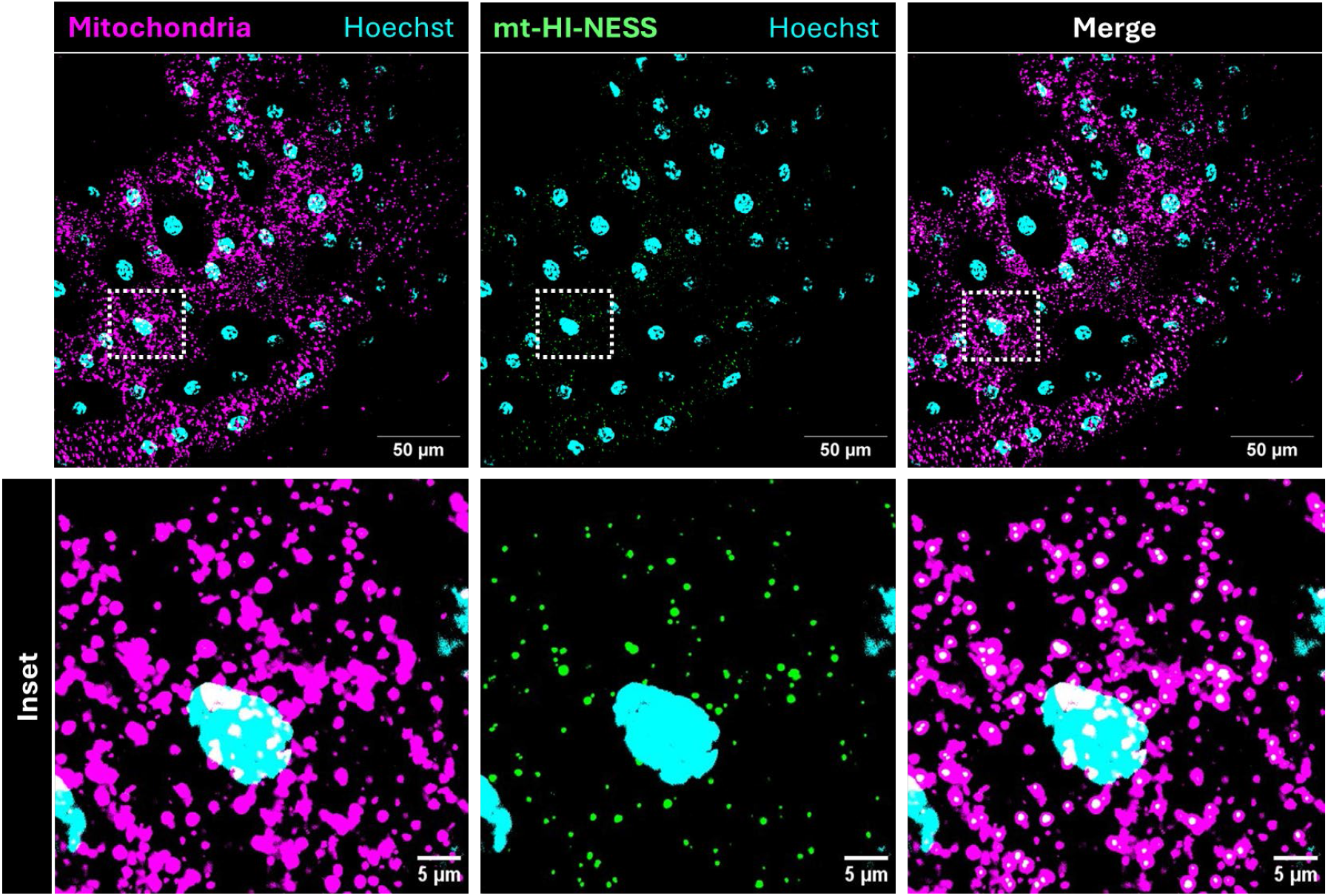
Mt-Kaede-HI-NESS in *D. melanogaster* larval fat body. Representative confocal images of the larval fat body tissue from dau-Gal4 crossed to UAS-Kaede mt-HI-NESS. The Kaede mt-HI-NESS (green) is localized within the mitochondrial network (magenta).

## Discussion

In this study, we developed, optimized, and validated mt-Hi-NESS reporters that enables specific labeling of mtDNA nucleoids. Building on the original HI-NESS genetic reporter for DNA labeling^37^, we generated a mitochondrially targeted version suitable for live-cell imaging of mtDNA. Using mt-Kaede-Hi-NESS, we demonstrated that this reporter can be broadly applied to examine nucleoid morphology and dynamics, as well as to monitor both cytosolic and extracellular mtDNA release events.

As with other exogenously expressed proteins, high expression levels of mt-Hi-NESS can introduce artefacts or perturb mitochondrial homeostasis. Consistent with this potential complication, our analysis revealed that elevated reporter expression can alter mtDNA copy number and nucleoid morphology. Based on these findings, we recommend only using cells expressing low levels of the reporter for analysis. As a rapid and reliable indicator of appropriate expression levels cells, imaged cells should be screened to confirm normal nucleoid number and morphology without evidence of clustering or clumping. While the reporter can be used following transient transfection, where a range of expression levels can be observed, we recommend using lentiviral transduction to generate stable cell lines where single-cell clones are established. In single cell clones, mtDNA copy number can also be assessed by qPCR to screen for clones with mtDNA levels equal to untransduced control cells, as these clones will maintain normal mitochondrial physiology and represent optimal lines for downstream imaging and analysis.

With this optimized expression framework established, we next sought to expand the color palette of mt-Hi-NESS to broaden its applicability across diverse imaging applications. To this end, we generated reporter variants fused to TagRFP and KillerRed. Surprisingly, both TagRFP- and KillerRed-based reporters exhibited diffuse mitochondrial labeling rather than discrete nucleoid localization. Through comparative analysis of DsRed-derived monomeric, dimeric, and tetrameric variants we showed that only tetrameric fluorescent proteins reliably produced punctate nucleoid labeling. To overcome this challenge and enable the use of monomeric, we engineered a “double-Hi-NESS” architecture by introducing a second H-NS-binding domain at the N-terminus. This modification substantially improved nucleoid binding for the reporter with monomeric TagRFP, thereby expanding the flexibility of the mt-Hi-NESS platform and enabling compatibility with a wider range of fluorophores and imaging modalities while avoiding reliance on tetrameric backbones.

The development of additional mt-HI-NESS variants to take advantage of the different applications of various fluorescent proteins will further expand the applications of the mt-HI-NESS reporters. However, when using the mt-HI-NESS reporters, it will be important to consider the specific fluorescent protein used, as well as to consider the possible effects of overexpression. As we have seen in budding yeast and flies, the issue of overexpression is also relevant in other organisms, but this limitation should be able to be mitigated using various genetic strategies. Nonetheless, that experimental conditions will likely need to be optimized and determined for each reporter in each model. Despite this potential limitation, we have shown that the mt-Kaede-HI-NESS can be used to study a variety of nucleoid dynamics including changes in size, number and distribution, as well as both cytosolic and extracellular release of mtDNA in live cell imaging experiments. While our current here efforts have focused on U2OS cells, the reporter is also applicable to other human cell types. Beyond human cells, the mt-HI-NESS reporter can also be adapted to label mtDNA in other organisms including *A. thaliana*^69^, S. cerevisiae, and D. melanogaster, and we expect it will be broadly applicable to several additional organisms.

Ultimately, these reporters will allow researchers to address questions about how nucleoids are distributed throughout the mitochondrial network, how mtDNA replication is coordinated with mitochondrial fission and fusion events, and to study the release of mtDNA into the cytosol or its endosomal/vesicle-mediated transport to autophagosomes for degradation^58,74,75^. Better tools to study this processes will be important, as mtDNA release and dynamics are increasingly recognized to be important in the context of human diseases^76–78^.

## Methods

### Cell culture

HEK-293 and U2OS cells were cultured in Dulbecco’s modified Eagle’s medium (DMEM) (Gibco, 11965092) supplemented with 10% fetal bovine serum (FBS) (Fisher, 12483020). All cells were maintained at 37 °C and 5% CO2 incubator. The U2OS TFAM-/+ 1C11 line described previously^58^, was a gift from Dr. Laura Newman (University of Virginia) and Dr. Gerald Shadel (Salk Institute). U2OS MFN2 knockout cells were kindly gifted by Dr. Edward Fon (McGill University)^54^. For the mtDNA release studies, we added 20 μg/ml LPS for 12 hours (InvivoGen, tlrl-pb5lps) or 200 nM Dexamethasone (Sigma-Aldrich, D4902) for 18 hours to cells. Cells were subsequently imaged for mtDNA localization. Extra-mitochondrial DNA was analyzed using the mitoQC plugin on FIJI^79^, while the mitochondrial network was quantified using MiNA^80^.

### Protein Structure Modeling

To model the predicted structures of our novel chimeric mt-HI-NESS proteins, we used Alphafold2^81,82^ (https://colab.research.google.com/github/sokrypton/ColabFold/blob/main/bet_a/AlphaFold2_advanced.ipynb), and the 3D structures were visualized using Chimera software^83^. To model predicted interactions between our mt-HI-NESS proteins and mitochondrial DNA (mtDNA) we used the online application HADDOCK (YYY)^84^. The 3D co-ordinates of the mtDNA mitochondrial light strand promotor sequence (5’TTAACAGTCACCCCCCAACTAA3’) were obtained from a previously reported crystal structure^85^. A range of active site residues YSYVDENGETKTWTGQGRTP was provided to the server based on previous reports^86^ and all other parameters were set to default settings within the HADDOCK web application (https://wenmr.science.uu.nl/haddock2.4/). The complex with the highest binding energy score was used to generate the figures, which were visualized using the Chimera software.

### HI-NESS plasmids

The mt-HI-NESS reporters were synthesized by VectorBuilder (Chicago, IL, USA) and cloned into a mammalian gene expression lentiviral vector following the hPGK Promoter. The vector contains ampicillin and puromycin resistance markers. Vector maps and DNA sequences are provided in the supplemental material, and the vector IDs, which can be used to retrieve detailed information about the vector of vectorbuilder.com can be found in (Fig. S1)

### Lentivirus Production

HEK 293T were cells seeded into 100 mm dishes and incubated until they reached approximately 80% confluency. The plasmids containing mt-HI-NESS constructs (2µg) were transfected into HEK cells along with the packaging (psPAX2; 1.5 µg) and envelope (pMD2.G; 0.5 µg) plasmids using either FuGENE 6 or Lipofectamine 3000 transfection reagent (both from Invitrogen, Thermo Fisher Scientific, Burlington, Ontario, Canada) according to the manufacturer protocol. The medium was changed after cells were incubated at 37 °C and 5% CO2 for about 16 hrs. The culture medium containing lentivirus particles were collected over 48-96 hrs post-transfection. Lenti-X concentrator from Takara bio (Cat. Nos. 631231 & 631232) was used to according to manufacturer protocol to concentrate the viral particle.

### Generation of stable cell lines

U2OS cells were seeded in a 6-well plate (50,000 cells/well) and cultured to reach 70% confluency (1-2 days). Fresh medium containing 8 µg/mL polybrene was added to cells, along with 100µL of containing concentrated lentivirus particles, for a total of 2 mL. Infected cells were incubated at 37 °C and 5% CO2 overnight at which point fresh culture medium without polybrene was added. After 24 hrs, the media was replaced again, this time containing puromycin (2 µg/mL) to select for stably transduced cells. Cells were grown for two weeks of selection, with puromycin-containing media changed every 2-3 days. To establish a genetically homogeneous population of cells, individual cells were obtained by serial dilution and single colonies picked. Cells that were not selected with puromycin are sorted using Fluorescent Activated Cell Sorting.

### Fluorescent Activated Cell Sorting

Cells were sorted using a Sony SH800 cell sorter. Briefly, trypsinized, washed cells were suspended at a concentration of 1X10^6^ cells/mL in the growth medium and filtered using a 40 µm cell strainer (VWR: 10199-655). Positive cells were then sorted based on fluorescent intensity in the GFP, or mCherry channel into 15 mL collection tubes using a sterile 100 µm chip (Sony: LE-C3210).

### Flow cytometry

Median fluorescent intensity (MFI) of mt-HI-NESS cells was acquired using a Becton-Dickson FACS CANTO cytometer supported by the BD FACS Diva software (BD Biosciences). Briefly, U2OS cells were washed with PBS, trypsinized and re-suspended in flow buffer (PBS with 1% BSA) prior to staining with the LIVE/DEAD™ Fixable Near IR (Invitrogen, L34992) kit per manufacturer’s instructions. Mt-HI-NESS signal was detected using a 488 nm laser with the FITC detector and live/dead signal was detected using a 633 nm laser with the APC-Cy7 detector. Live-single cells were gated based on FSC, SSC and low APC-Cy7 fluorescence from which MFI of mt-HI-NESS was reported from the FITC channel from three independent experiments.

### Sample preparation for live-cell imaging

Unless otherwise indicated, all reagents and cell culture media components were obtained from ThermoFisher Scientific (Markham, Ontario, Canada). Transduced cells expressing different variants of the HI-NESS constructs were cultured in 35 mm dishes with a 1.5 glass coverslip bottom (Cellvis, D35-20-1.5-N). Imaging was performed either 1 or 2 days after seeding the cells in the dishes. For some experiments, an additional extrinsic fluorescent label was also applied to the cells (e.g., SYBR Gold (S11494), Mitoblue BioTracker 405 (SCT135), MitoTracker Green FM (M7514) or MitoTracker Deep Red FM (M22426)). Stock solutions of each probe were prepared in sterile anhydrous DMSO (D12345) (stock concentrations: SYBR Gold 1:1000; MitoTracker Green 1 mM; MitoTracker Deep Red 100 uM). To label the cells, media was aspirated, and fresh and pre-warmed media containing 1:1000 dilutions of SYBR Gold, MitoTracker Green, or MitoTracker Deep Red stock solutions were added. Cells were incubated for 1 hr, washed once with sterile PBS, and fresh media was then added to the wells. Cells were then imaged on an LCI system stage (Quorum Technologies, Guelph, Ontario, Canada), where temperature, humidity, and pH (5% CO2) were maintained.

### Immunofluorescence sample preparation (Cell fixing)

Approximately 120,000 cells were seeded and grown overnight on circular 12mm #1.5 coverslips Cat #72230-10 in 24 well plate (Cat.# MBC.TC.F.24). Media was aspirated and cells were washed once with 0.5mL of pre-warmed PBS followed by fixation using 0.5ml of pre-warmed 4% paraformaldehyde (Electron Microscopy Sciences Cat: 15710) for 12-15 mins at 37°C. After fixation, cells were washed 3 times with 0.5mL of PBS, followed by quenching using 0.5mL of 50 mM NH_4_Cl for 15 min with three more PBS washes.

### Immunofluorescence

Fixed cells were permeabilized and blocked in a one-step reaction with 0.3% Triton-X in PBS with 10% Serum (FBS) for 1hr at room temperature. Primary antibodies were diluted in 0.1% Triton-X in PBS with 5% Serum (FBS) and incubated overnight at 4°C. Secondary antibodies were diluted in 0.1% Triton-X in PBS with 5% Serum (FBS) and incubated for 1hr at room temperature. Cells were then washed in PBS 3 times and mounted onto glass slides with DAKO fluorescent mounting media (S302380-2). Primary and secondary antibody data is found in the supplementary materials.

### Confocal imaging

Z-stack confocal images were acquired using an Olympus IX83 inverted spinning disk confocal microscope equipped with SpinSR10 system operated by CellSens Dimension software. The confocal system was equipped with lasers emitting at 405 nm (Ch1), 488 nm (Ch2), 561 nm (Ch3), and 640 nm (Ch4), with an ORCA-Flash4.0 V3 Digital CMOS camera (Hamamatsu Photonics K.K.). Spectral separation was effected using 89902 ET - 405/488/561/647nm Laser Quad Band Set (Chroma) and FWHM band pass can be found in the table below. Images were acquired at 20% laser power and with an exposure of 200ms at 14fps for all channels. One or more channels were selected, depending on the fluorophores used in the imaging experiment.

|  | Laser | Band Name | Center Wavelength (nm) | Bandwidth (nm) |
| --- | --- | --- | --- | --- |
| Ch1 | 405 | DAPI | 447 | 60 |
| Ch2 | 488 | GFP | 520 | 35 |
| Ch3 | 561 | RED | 617 | 73 |
| Ch4 | 640 | Cy5 | 684 | 40 |

### Nucleoid Number and Size Analysis

Nucleoid number and size were quantified in FIJI/ImageJ (doi:10.1038/nmeth.2019). For each field of view, TOM20 and dsDNA channels were merged and a maximum-intensity projection was generated. Whole-cell ROIs were manually delineated using the TOM20 channel to define the mitochondrial network and cell boundary (ROIs were drawn to avoid nuclear regions). ROIs were applied to the dsDNA channel, and nucleoid-like objects were segmented using Auto Local Thresholding (Phansalkar method; radius = 15). Thresholded images were converted to binary masks and analyzed using the built-in Analyze Particles function with an area filter of **0.05–2.5 µm**^**2**^ (image calibrated) and circularity **0.30–1.00**. The same segmentation parameters were applied to all images for reproducibility. Batch processing was performed using a custom ImageJ macro (Supp. data), which also saved ROI sets and binary masks for auditability.

### Western blotting

Cell pellets (∼2,000,000 cells) were collected after trypsinization, washed with phosphate buffered saline (PBS) and lysed with RIPA buffer (ThermoFisherScientific, 89900) containing protease inhibitors. The cell lysates (20-50 μg of proteins) were resolved on SDS-PAGE gels (12%) and transferred onto PVDF Membranes in 25mM Tris, 192mM glycine and 20% (vol/vol) methanol at 30V overnight at 4°C. After blocking with 5% nonfat milk in TBST (25mM Tris, pH7.4, 150mM NaCl, 0.05% Tween-20), the blots were washed and incubated at RT for 2-3hrs with the following primary antibodies at the indicated dilutions; OxPhos antibody cocktail (Abcam, ab110411; 1:1000 dilution), anti-Actin (Sigma, A5316; 1:1000), anti-HSP60 (Cell Signaling Technology; 1:1000), anti-VDAC1 (Abcam, ab14734; 1:1000), anti-HA tag (Santa Cruz Biotechnology, sc-7392; 1:1000), HA-Tag (C29F4) Rabbit Monoclonal Antibody #3724 1:1000, and CD13 (homemade, clone AD-1^61^, diluted to 1 μg/ml).

Appropriate horseradish peroxidase (HRP)-conjugated secondary antibodies were used as follows: goat anti-rabbit IgG HRP linked Antibody (Cell Signaling Technology, 7074S, 1:2000), or goat anti-mouse IgG HRP (Santa Cruz Biotechnology, sc-2055;1:2000). After washing with TBST (4x) the blots were developed with SuperSignal West Femto enhanced chemiluminescence substrate (Pierce Chemical, Rockford, IL). The signal was detected and quantified with a chemiluminescence imaging analyzer (LAS3000 mini, Fujifilm).

### Quantification of mtDNA copy number

Total DNA was isolated from cell pellets (∼1,500,000 cells) using the E.Z.N.A. Tissue DNA Kit, Omega Bio-tek (VWR Scientific, D6905-03), according to the manufacturer’s protocol. Isolated DNA (50-100 ng) was used for quantitative PCR (QPCR) to determine mtDNA copy number, as reported previously^16^. Primers used are as bellow: MT forward: CACCCAAGAACAGGGTTTGT; Reverse: TGGCCATGGGTATGTTGTTAA; 18S forward: TAGAGGGACAAGTGGCGTTC reverse:CGCTGAGCCAGTCAGTGT. Briefly, mtDNA and 18S DNA were amplified using PowerUp SYBR Green Master Mix (Thermo Fisher Scientific, A25742) in the QuantStudio 6 Flex Real-Time PCR system (Thermo Fisher Scientific) machine, and the delta delta Ct method was used to determine mtDNA copy number relative to the 18S. Reactions were performed in triplicate and mtDNA copy number analysis was performed on six independent biological replicates.

### Extracellular vesicles

For analysis of extracellular vesicles, cells were treated with 2 ug/ml LPS overnight, in serum-free media. For western blot and qPCR analysis pellets were extracted after ultracentrifugation at 100,000 g. For immunocapture and subsequent imaging, 20 mm live cell imaging dishes were coated overnight at 4 degrees with a homemade monoclonal antibody against CD13 (clone AD-1)^61^. After washing with PBS, the cell media supernatant (centrifuged for 2000 g for 20 minutes) was added to the antibody coated dish and incubated overnight at 4 degrees. The dishes were subsequently washed with PBS and the CD13(AD-1) mouse antibody prelabelled using Zenon antibody labelling kit Alexa Fluor 568 (Thermo Fisher Scientific, Z25006) was added to the dish. The coverslips were then imaged with confocal microscopy. Colocalization was performed using the JaCOP plugin on FIJI^87^.

### Yeast-specific methods

All strains were derived from W303, constructed using standard lithium acetate transformation, and verified by sequencing. Yeast strains used in this study are listed in Supplementary Table 1.

### Microscopy

After inoculation in SCD (synthetic complete medium with 2% glucose), cells were grown for at least 24 hours while maintaining OD below 1 through appropriate dilutions. Coverslips (µ-Slide 8 Well, ibi-Treat, Ibidi) were covered with 200 μL concanavalin A (1 mg/mL), incubated for 5-10 min, and washed twice with double-distilled water. 1 mL of the cell culture was sonicated and 200 μL were transferred to a well. After about 5 min, the supernatant was removed, and cells were washed twice with filtered SCD. Finally, 200 μL filtered SCD was added to cover the wells.

Live cells were then imaged with a Zeiss LSM 800 confocal microscope, using a Plan-Apochromat 63x/1.4 NA Oil DIC objective and obtaining z-stacks of 46 slices with 0.22 μm increments. mKate2 was imaged with an excitation wavelength of 561 nm, detecting emission between 615 and 700 nm. mt-Kaede-HI-NESS was imaged with an excitation wavelength of 488lZnm and detection between 410 and 546lZnm. Bright-field images were obtained with the transmitted light detector (T-PMT).

### Image analysis

Image analysis was performed using Cell-ACDC^88^ and SpotMAX ^72^. First, raw microscopy files were converted to TIFF using the Bioformats library^89^. Using the brightfield channel, cells were segmented with YeaZ v1.0.3^90^ embedded in Cell-ACDC. Segmentations were then carefully inspected and corrected where necessary, and mother-bud pairs were annotated.

To count the number of nucleoids and quantify the mitochondrial network volume, we used the software tool SpotMAX. The analysis is fully automatic and it was previously described^72^. The mitochondrial network volume was quantified by segmenting the mKate2 channel in 3D and summing the positive voxels. The analysis steps to segment the mitochondrial network are the following: 1) 3D Gaussian filter with sigma = 0.75, 2) 3D Sato filter to enhance the tubular structure with sigmas = (1.0, 2.0)^91^, 3) automatic thresholding of the filtered image using the Yen algorithm^92^, 4) connected-component labelling and removal of objects with less than 10 voxels volume (artefacts). To count the number of spots, we used SpotMAX with the Li algorithm^93^for semantic segmentation and local peak detection. Note that we distinguish between all detected spots (shown in Fig. S6) and high-confidence spots (shown in Fig. 8). The high-confidence spots are those spots that pass the SpotMAX quality control consisting of two tests: a Welch’s t-test and a minimum Glass’ effect size between the Kaede and mKate2 normalised intensities within a spheroid with user-selected size centered at each spot. The spheroid radii were determined through visual inspection and they are 0.453 μm in the xy-directions (twice the Abbe’s diffraction limit) and 1 μm in the z-direction. The valid spots are those where the t-statistic of the t-test is positive, the p-value is below 0.025, and the Glass’s effect size is greater than 1.5. More details about this strategy can be found as published previously^72^.

The signal-to-noise ratio (SNR) shown in Fig. S6C was calculated with the following formula:

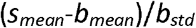

Here, *s*_mean_ is the mean of the raw pixel intensities inside the resolution-limited volume centered at each spot, and *b*_mean_ and *b*_std_ are the mean and the standard deviation of the background pixels, respectively. The background pixels are those inside the segmented mitochondrial network and outside the spots’ masks. The spots’ masks are constructed by placing a spheroid (with the same radii described above) on each detected spot (all spots).

### mtDNA copy number quantification with qPCR

mtDNA copy numbers were determined largely following the strategy described previously^71^. After inoculation in SCD, cells were grown for at least 22 hours to a final OD between 0.1 and 0.5, always maintaining OD below 1 through appropriate dilutions. For each culture, 1 mL was sonicated and used to measure cell volume at a range of 10 to 328 fL using a Coulter counter (Beckman Coulter, Z2 Particle Counter), and another 1 mL was sonicated and used to determine bud fractions through visual inspection with a microscope (at least 200 cells were counted per replicate). For DNA extraction, 10 mL culture was centrifuged at 3434 g for 5 mins. The pellet was washed with 1 mL double-distilled water and centrifuged again for 1 min at 11600 g. The supernatant was removed, and pellets were frozen in liquid nitrogen and stored at-20°C.

DNA was extracted by phenol-chloroform-isoamyl alcohol (PCI) extraction. DNA pellets were resuspended in 200 µL DNA extraction buffer (2% Triton X-100, 1% SDS, 100 mM NaCl, 10 mM TRIS, 1 mM EDTA; pH = 8.0), and then 200 µL PCI and glass beads were added. Cells were mechanically disrupted by vortexing for 30 seconds at 3000 oscillations per minute (Mini-BeadBeater 24, 230 V, BioSpec Products). After centrifugation at 11600 g for 5 min, the aqueous phase was collected, and DNA was precipitated by mixing with 500 µL 100% ethanol and centrifugation at 11600 g for 5 min. Next, the pellet was washed with 700 µL 70% ethanol, and after complete removal and evaporation of the alcohol resuspended in 50 µL nuclease-free water. RNA residues were removed by 30 min incubation with RNase A (DNase-free; 1 mg/mL) at 37°C. PCI extraction was then repeated as described above. Finally, the DNA was resuspended in 50 µL nuclease-free water and stored at-20°C until further use.

To determine mtDNA concentrations, qPCRs were performed in technical triplicates using a LightCycler 480 Multiwell Plate 96 (Roche), using 1 ng DNA per replicate. A DNA-binding fluorescent dye (BioRad, SsoAdvanced Universal SYBR Green Supermix) and specific primers (Supplementary Table S2) for two nuclear DNA genes (ACT1 and MRX6) and two mtDNA genes (COX2 and COX3) were used. Single technical replicates were excluded from further analysis when the standard deviation between the three replicates was above 0.5. For each gene, Cq values of the remaining replicates were then averaged, and DNA concentrations were calculated using a linear fit to a gene-dependent calibration curve^71^. Concentrations of nuclear DNA and mtDNA were then calculated by averaging the concentrations of the respective genes. The ratio of mtDNA and nuclear DNA concentration was then used together with the average nuclear DNA copy number per cell as estimated from the bud fractions to calculate the average mtDNA copy number per cell^71^.

### *Drosophila*-specific methods

#### Drosophila husbandry

Drosophila stocks and experimental larvae were raised at room temperature on food consisting of 770 g Torula yeast, 1600 g cornmeal, 675 g sucrose, 2340 g D-glucose, 150 g agar, and 240 mL of an acid mixture (propionic acid/phosphoric acid) for every 34 L of water.

#### Drosophila stocks and genetics

For Drosophila experiments, da-Gal4 (BDSC #95282) line was crossed to UAS Kaede-HI-NESS (generated for this study). Hatched first instar larvae of the F1 progeny were then transferred into vials with food (50 larvae per vial). Experiments were conducted on larvae at 120 hrs after egg-laying (AEL).

#### Kaede HI-NESS stock generation

For Drosophila melanogaster studies, Kaede HI-NESS (from this paper) was cloned into a pDestination UAS-plasmid and purified plasmid was microinjected into flies carrying attP2 3L insertion sites to generate stable transgenic flies (injections were carried out by GENOMEPROLAB, Montreal, Canada).

#### Tissue Dissection, MitoTracker staining and Microscopy

Larvae (120hrs AEL) were inverted in Schneider’s Drosophila medium (Gibco, 21720024) to expose the fat body. Inverted carcasses were incubated in in Schneider’s media containing MitoTracker Deep Red (1:1000; Thermo Fisher Scientific, M22426) and Hoechst (1:5000; Thermo Fisher Scientific, 62249) for 30 minutes. Inverted larvae were then washed three times with schneider’s media and fat body tissues were then immediately mounted in Vectasheild (Vector laboratories, H1000) mounting media on a glass coverslip and subsequently imaged.

## Acknowledgements & Funding

We would like to thank Dr. Laura Newman (University of Virginia), Dr. Gerald Shadel (Salk Institute), and Dr. Edward Fon (McGill University) for sharing CRISPR edited U2OS cell lines used in the studies. Work was supported by funds provided by a Natural Sciences and Engineering Research Council of Canada (NSERC) Discovery grant (TES) and a Crohn’s & Colitis Canada Innovations grant (TES & PC). JOD was supported by an HBI International Graduate Recruitment Scholarship. AJ was supported by a President’s Doctoral Recruitment Scholarship in Transdisciplinary Research through the University of Calgary. We also thank the Cumming Medical Research Fund and the Snyder Institute for support of the imaging-based experiments. The funding sources were not involved in the study design, data collection and analysis, the writing of the report, nor in the decision to submit.

## Author contributions

JOD designed, performed, and analyzed experiments, prepared figures, and wrote the manuscript. JD designed, performed, analyzed experiments, and prepared figures. LS designed, performed, and analyzed experiments, prepared figures, and edited the manuscript. MZ designed, performed, and analyzed experiments, and prepared figures. FS performed and analyzed experiments, prepared figures, and edited the manuscript. DB designed, performed, and analyzed experiments. FP performed image analysis, prepared figures and edited the manuscript. AB performed and analyzed experiments, and edited the manuscript. AS designed, performed, and analyzed experiments. AJ performed experiments and prepared figures. AM performed experiments. CB performed and analyzed experiments, and edited the manuscript. SG supervised the study. KMS designed performed and analyzed experiments, supervised the study, wrote portions of the manuscript. PC conceived and designed the study, designed and analyzed experiments, supervised the study, prepared figures and wrote and revised the manuscript. TES conceived and designed the study, analyzed experiments, prepared figures, supervised the study, and wrote and revised the manuscript.

**Figure 2. Using Mt-Kaede-HI-NESS to Visualize mtDNA Release in Models of Mitochondrial Dysfunction.. (A)** Representative confocal images showing mitochondria (MitoTracker DeepRed) and mtDNA (mt-Kaede-HI-NESS) in transduced (WT U2OS) and transfected cell models (WT U2OS and U2OS MFN2 KO). **(B-C)** Quantitative analyses of **(B)** number and **(C)** size of mtDNA nucleoids in transfected vs transduced cell models (U2OS WT/MFN2 KO). Colours indicate biological replicates, violins indicate median and interquartile range, multiple unpaired t-tests. Stats: ns P > 0.05; *P ≤ 0.05; ****P ≤ 0.0001. **(D)** Representative confocal images showing mitochondria (MitoTracker Deep Red) and mtDNA (mt-Kaede-HI-NESS) in U2OS MFN2 knockout cells and U2OS TFAM 1C11 cells, circles indicate mtDNA not localized to the mitochondrial network. **(E)** Quantitative analysis of number of mtDNA puncta not localized to the mitochondrial network, colours indicate biological replicates, lines indicate mean +/-SD, two-way ANOVA. **(F-G)** Representative confocal images showing drug-induced extra-mitochondrial mtDNA, as shown by addition of **(F)** LPS and **(G)** Dexamethasone. **(H-G)** Quantitative analyses of drug-based models of mtDNA release, showing (H) number of nucleoids, **(I)** number of extra-mitochondrial nucleoids and **(J)** mean mitochondrial branch length. Colours indicate biological replicates, lines indicate mean +/-SD, two-way ANOVA. Stats: ns P > 0.05; ****P ≤ 0.0001.

**Figure 3. Extracellular mtDNA reported using mt-Kaede-HI-NESS. (A)** Schematic of the cell-free mtDNA detection assay. **(B)** Representative confocal images showing mtDNA (mt-Kaede-HI-NESS) and extra-cellular vesicles (anti-CD13) in LPS treated and untreated cells, circles indicate presence of mtDNA. C-H) Quantification of extra-cellular mtDNA showing. **(C)** Pearson’s co-localization coefficient between green and magenta signals, number of **(D)** extra-cellular vesicles and **(E)** extra cellular mtDNA puncta, size of **(F)** extra-cellular vesicles and **(G)** extra cellular mtDNA puncta and **(H)** percentage of extracellular vesicles that contain mtDNA. All violins show median and interquartile range, unpaired t-test. **(I)** Relative mtDNA copy number in the extracellular fraction for LPS treated and untreated cells, bars indicate mean +/-SD, unpaired t-test. **(J)** Representative bands from western blot analysis of the extracellular fraction for LPS treated and untreated cells, showing relative protein amounts of CD-13 and HA-tag in LPS treated/untreated cells. Stats: *P ≤ 0.05; **P ≤ 0.01; ****P ≤ 0.0001.

## References

1. Anderson S, Bankier AT, Barrell BG, et al. Sequence and organization of the human mitochondrial genome. Nature. 1981;290(5806):457–465. doi:10.1038/290457a0

2. Kaufman BA, Durisic N, Mativetsky JM, et al. The Mitochondrial Transcription Factor TFAM Coordinates the Assembly of Multiple DNA Molecules into Nucleoid-like Structures. Mol Biol Cell. 2007;18(9):3225–3236. doi:10.1091/mbc.E07-05-0404

3. Parisi MA, Clayton DA. Similarity of human mitochondrial transcription factor 1 to high mobility group proteins. Science. 1991;252(5008):965–969. doi:10.1126/science.2035027

4. Kukat C, Wurm CA, Spåhr H, Falkenberg M, Larsson NG, Jakobs S. Super-resolution microscopy reveals that mammalian mitochondrial nucleoids have a uniform size and frequently contain a single copy of mtDNA. Proc Natl Acad Sci U S A. 2011;108(33):13534–13539. doi:10.1073/pnas.1109263108

5. Nicholls TJ, Gustafsson CM. Separating and Segregating the Human Mitochondrial Genome. Trends Biochem Sci. 2018;43(11):869–881. doi:10.1016/j.tibs.2018.08.007

6. Chapman J, Ng YS, Nicholls TJ. The Maintenance of Mitochondrial DNA Integrity and Dynamics by Mitochondrial Membranes. Life Basel Switz. 2020;10(9). doi:10.3390/life10090164

7. Sabouny R, Shutt TE. The role of mitochondrial dynamics in mtDNA maintenance. J Cell Sci. 2021;134(24):jcs258944. doi:10.1242/jcs.258944

8. Semadhi MP, Mulyaty D, Halimah E, Levita J. Healthy mitochondrial DNA in balanced mitochondrial dynamics: A potential marker for neuro⍰aging prediction (Review). Biomed Rep. 2023;19(3):64. doi:10.3892/br.2023.1646

9. Lewis SC, Uchiyama LF, Nunnari J. ER-mitochondria contacts couple mtDNA synthesis with mitochondrial division in human cells. Science. 2016;353(6296):aaf5549. doi:10.1126/science.aaf5549

10. Ilamathi HS, Benhammouda S, Lounas A, et al. Contact sites between endoplasmic reticulum sheets and mitochondria regulate mitochondrial DNA replication and segregation. iScience. 2023;26(7):107180. doi:10.1016/j.isci.2023.107180

11. Silva Ramos E, Motori E, Bruser C, et al. Mitochondrial fusion is required for regulation of mitochondrial DNA replication. PLoS Genet. 2019;15(6):e1008085. doi:10.1371/journal.pgen.1008085

12. Ishihara T, Ban-Ishihara R, Maeda M, et al. Dynamics of mitochondrial DNA nucleoids regulated by mitochondrial fission is essential for maintenance of homogeneously active mitochondria during neonatal heart development. Mol Cell Biol. 2015;35(1):211–223. doi:10.1128/MCB.01054-14

13. Ban-Ishihara R, Ishihara T, Sasaki N, Mihara K, Ishihara N. Dynamics of nucleoid structure regulated by mitochondrial fission contributes to cristae reformation and release of cytochrome c. Proc Natl Acad Sci U S A. 2013;110(29):11863–11868. doi:10.1073/pnas.1301951110

14. Ilamathi HS, Ouellet M, Sabouny R, et al. A new automated tool to quantify nucleoid distribution within mitochondrial networks. Sci Rep. 2021;11(1):22755. doi:10.1038/s41598-021-01987-9

15. Ishihara T, Ban-Ishihara R, Ota A, Ishihara N. Mitochondrial nucleoid trafficking regulated by the inner-membrane AAA-ATPase ATAD3A modulates respiratory complex formation. Proc Natl Acad Sci U S A. 2022;119(47):e2210730119. doi:10.1073/pnas.2210730119

16. Sabouny R, Wong R, Lee-Glover L, et al. Characterization of the C584R variant in the mtDNA depletion syndrome gene FBXL4, reveals a novel role for FBXL4 as a regulator of mitochondrial fusion. Biochim Biophys Acta Mol Basis Dis. 2019;1865(11):165536. doi:10.1016/j.bbadis.2019.165536

17. Donkervoort S, Sabouny R, Yun P, et al. MSTO1 mutations cause mtDNA depletion, manifesting as muscular dystrophy with cerebellar involvement. Acta Neuropathol (Berl). 2019;138(6):1013–1031. doi:10.1007/s00401-019-02059-z

18. Chen H, McCaffery JM, Chan DC. Mitochondrial fusion protects against neurodegeneration in the cerebellum. Cell. 2007;130(3):548–562. doi:10.1016/j.cell.2007.06.026

19. Newman LE, Shadel GS. Mitochondrial DNA Release in Innate Immune Signaling. Annu Rev Biochem. Published online March 31, 2023. doi:10.1146/annurev-biochem-032620-104401

20. Caicedo A, Benavides-Almeida A, Haro-Vinueza A, et al. Decoding the nature and complexity of extracellular mtDNA: Types and implications for health and disease. Mitochondrion. 2024;75:101848. doi:10.1016/j.mito.2024.101848

21. Prole DL, Chinnery PF, Jones NS. Visualizing, quantifying and manipulating mitochondrial DNA in vivo. J Biol Chem. Published online 15 2020. doi:10.1074/jbc.REV120.015101

22. Ashley N, Harris D, Poulton J. Detection of mitochondrial DNA depletion in living human cells using PicoGreen staining. Exp Cell Res. 2005;303(2):432–446. doi:10.1016/j.yexcr.2004.10.013

23. Bereiter-Hahn J, Vöth M. Distribution and dynamics of mitochondrial nucleoids in animal cells in culture. Exp Biol Online. 1996;1(4):1–17. doi:10.1007/s00898-996-0004-1

24. Jevtic V, Kindle P, Avilov SV. SYBR Gold dye enables preferential labelling of mitochondrial nucleoids and their time-lapse imaging by structured illumination microscopy. PloS One. 2018;13(9):e0203956. doi:10.1371/journal.pone.0203956

25. Jevtic V, Kindle P, Avilov SV. Specific Labeling of Mitochondrial Nucleoids for Time-lapse Structured Illumination Microscopy. J Vis Exp JoVE. 2020;(160). doi:10.3791/60003

26. He J, Mao CC, Reyes A, et al. The AAA+ protein ATAD3 has displacement loop binding properties and is involved in mitochondrial nucleoid organization. J Cell Biol. 2007;176(2):141–146. doi:10.1083/jcb.200609158

27. Alexeyev M. TFAM in mtDNA Homeostasis: Open Questions. DNA. 2023;3(3):134–136. doi:10.3390/dna3030011

28. Rajala N, Gerhold JM, Martinsson P, Klymov A, Spelbrink JN. Replication factors transiently associate with mtDNA at the mitochondrial inner membrane to facilitate replication. Nucleic Acids Res. 2014;42(2):952–967. doi:10.1093/nar/gkt988

29. Ekstrand MI, Falkenberg M, Rantanen A, et al. Mitochondrial transcription factor A regulates mtDNA copy number in mammals. Hum Mol Genet. 2004;13(9):935–944. doi:10.1093/hmg/ddh109

30. Bonekamp NA, Jiang M, Motori E, et al. High levels of TFAM repress mammalian mitochondrial DNA transcription in vivo. Life Sci Alliance. 2021;4(11). doi:10.26508/lsa.202101034

31. Ikeda M, Ide T, Fujino T, et al. Overexpression of TFAM or twinkle increases mtDNA copy number and facilitates cardioprotection associated with limited mitochondrial oxidative stress. PloS One. 2015;10(3):e0119687. doi:10.1371/journal.pone.0119687

32. Ylikallio E, Tyynismaa H, Tsutsui H, Ide T, Suomalainen A. High mitochondrial DNA copy number has detrimental effects in mice. Hum Mol Genet. 2010;19(13):2695–2705. doi:10.1093/hmg/ddq163

33. Sumitani M, Kasashima K, Ohta E, Kang D, Endo H. Association of a novel mitochondrial protein M19 with mitochondrial nucleoids. J Biochem (Tokyo). 2009;146(5):725–732. doi:10.1093/jb/mvp118

34. Zhang J, Campbell RE, Ting AY, Tsien RY. Creating new fluorescent probes for cell biology. Nat Rev Mol Cell Biol. 2002;3(12):12. doi:10.1038/nrm976

35. Cranfill PJ, Sell BR, Baird MA, et al. Quantitative Assessment of Fluorescent Proteins. Nat Methods. 2016;13(7):557–562. doi:10.1038/nmeth.3891

36. Dairaghi DJ, Shadel GS, Clayton DA. Human mitochondrial transcription factor A and promoter spacing integrity are required for transcription initiation. Biochim Biophys Acta. 1995;1271(1):127–134. doi:10.1016/0925-4439(95)00019-z

37. Kremer LS, Gao G, Rigoni G, et al. Tissue-specific responses to TFAM and mtDNA copy number manipulation in prematurely ageing mice. eLife. 2025;14:RP104461. doi:10.7554/eLife.104461

38. Rashid FZM, Mahlandt E, van der Vaart M, et al. HI-NESS: a family of genetically encoded DNA labels based on a bacterial nucleoid-associated protein. Nucleic Acids Res. 2022;50(2):e10. doi:10.1093/nar/gkab993

39. Shindo H, Iwaki T, Ieda R, et al. Solution structure of the DNA binding domain of a nucleoid-associated protein, H-NS, from Escherichia coli. FEBS Lett. 1995;360(2):125–131. doi:10.1016/0014-5793(95)00079-o

40. Gordon BRG, Li Y, Cote A, et al. Structural basis for recognition of AT-rich DNA by unrelated xenogeneic silencing proteins. Proc Natl Acad Sci U S A. 2011;108(26):10690–10695. doi:10.1073/pnas.1102544108

41. Qin L, Erkelens AM, Ben Bdira F, Dame RT. The architects of bacterial DNA bridges: a structurally and functionally conserved family of proteins. Open Biol. 2019;9(12):190223. doi:10.1098/rsob.190223

42. Shutt TE, Gray MW. Bacteriophage origins of mitochondrial replication and transcription proteins. Trends Genet. 2006;22(2):90–95. doi:10.1016/j.tig.2005.11.007

43. Shutt TE, Gray MW. Homologs of mitochondrial transcription factor B, sparsely distributed within the eukaryotic radiation, are likely derived from the dimethyladenosine methyltransferase of the mitochondrial endosymbiont. Mol Biol Evol. 2006;23(6):1169–1179. doi:10.1093/molbev/msk001

44. Carrodeguas JA, Kobayashi R, Lim SE, Copeland WC, Bogenhagen DF. The accessory subunit of Xenopus laevis mitochondrial DNA polymerase gamma increases processivity of the catalytic subunit of human DNA polymerase gamma and is related to class II aminoacyl-tRNA synthetases. Mol Cell Biol. 1999;19(6):4039–4046. doi:10.1128/MCB.19.6.4039

45. Carrodeguas JA, Bogenhagen DF. Protein sequences conserved in prokaryotic aminoacyl-tRNA synthetases are important for the activity of the processivity factor of human mitochondrial DNA polymerase. Nucleic Acids Res. 2000;28(5):1237–1244.

46. Forkink M, Smeitink JAM, Brock R, Willems PHGM, Koopman WJH. Detection and manipulation of mitochondrial reactive oxygen species in mammalian cells. Biochim Biophys Acta BBA - Bioenerg. 2010;1797(6):1034–1044. doi:10.1016/j.bbabio.2010.01.022

47. Filippin L, Abad MC, Gastaldello S, Magalhães PJ, Sandonà D, Pozzan T. Improved strategies for the delivery of GFP-based Ca2+ sensors into the mitochondrial matrix. Cell Calcium. 2005;37(2):129–136. doi:10.1016/j.ceca.2004.08.002

48. Olenych SG, Claxton NS, Ottenberg GK, Davidson MW. The fluorescent protein color palette. Curr Protoc Cell Biol. 2007;Chapter 21:Unit 21.5. doi:10.1002/0471143030.cb2105s36

49. Ali SS, Beckett E, Bae SJ, Navarre WW. The 5.5 Protein of Phage T7 Inhibits H-NS through Interactions with the Central Oligomerization Domain⍰. J Bacteriol. 2011;193(18):4881–4892. doi:10.1128/JB.05198-11

50. Jumper J, Evans R, Pritzel A, et al. Highly accurate protein structure prediction with AlphaFold. Nature. 2021;596(7873):7873. doi:10.1038/s41586-021-03819-2

51. Wiedemann N, Pfanner N. Mitochondrial Machineries for Protein Import and Assembly. Annu Rev Biochem. 2017;86:685–714. doi:10.1146/annurev-biochem-060815-014352

52. Almagro Armenteros JJ, Salvatore M, Emanuelsson O, et al. Detecting sequence signals in targeting peptides using deep learning. Life Sci Alliance. 2019;2(5):e201900429. doi:10.26508/lsa.201900429

53. Fukasawa Y, Tsuji J, Fu SC, Tomii K, Horton P, Imai K. MitoFates: improved prediction of mitochondrial targeting sequences and their cleavage sites. Mol Cell Proteomics MCP. 2015;14(4):1113–1126. doi:10.1074/mcp.M114.043083

54. Vranas M, Lu Y, Rasool S, et al. Selective localization of Mfn2 near PINK1 enables its preferential ubiquitination by Parkin on mitochondria. Open Biol. 2022;12(1):210255. doi:10.1098/rsob.210255

55. Chen H, Vermulst M, Wang YE, et al. Mitochondrial fusion is required for mtDNA stability in skeletal muscle and tolerance of mtDNA mutations. Cell. 2010;141(2):280–289. doi:10.1016/j.cell.2010.02.026

56. Irazoki A, Gordaliza-Alaguero I, Frank E, et al. Disruption of mitochondrial dynamics triggers muscle inflammation through interorganellar contacts and mitochondrial DNA mislocation. Nat Commun. 2023;14(1):108. doi:10.1038/s41467-022-35732-1

57. West AP, Khoury-Hanold W, Staron M, et al. Mitochondrial DNA stress primes the antiviral innate immune response. Nature. 2015;520(7548):553–557. doi:10.1038/nature14156

58. Newman LE, Weiser Novak S, Rojas GR, et al. Mitochondrial DNA replication stress triggers a pro-inflammatory endosomal pathway of nucleoid disposal. Nat Cell Biol. Published online February 8, 2024. doi:10.1038/s41556-023-01343-1

59. Nguyen M, Collier JJ, Ignatenko O, et al. Parkinson’s genes orchestrate pyroptosis through selective trafficking of mtDNA to leaky lysosomes. bioRxiv. Preprint posted online September 12, 2023:2023.09.11.557213. doi:10.1101/2023.09.11.557213

60. Trumpff C, Marsland AL, Basualto-Alarcón C, et al. Acute psychological stress increases serum circulating cell-free mitochondrial DNA. Psychoneuroendocrinology. 2019;106:268–276. doi:10.1016/j.psyneuen.2019.03.026

61. Deng JT, Hoylaerts MF, Nouwen EJ, De Broe ME, Van Hoof VO. Purification of circulating liver plasma membrane fragments using a monoclonal antileucine aminopeptidase antibody. Hepatol Baltim Md. 1996;23(3):445–454. doi:10.1002/hep.510230308

62. Merzlyak EM, Goedhart J, Shcherbo D, et al. Bright monomeric red fluorescent protein with an extended fluorescence lifetime. Nat Methods. 2007;4(7):555–557. doi:10.1038/nmeth1062

63. Bulina ME, Chudakov DM, Britanova OV, et al. A genetically encoded photosensitizer. Nat Biotechnol. 2006;24(1):1. doi:10.1038/nbt1175

64. Costantini LM, Fossati M, Francolini M, Snapp EL. Assessing the tendency of fluorescent proteins to oligomerize under physiologic conditions. Traffic. 2012;13(5):643–649. doi:10.1111/j.1600-0854.2012.01336.x

65. Ando R, Hama H, Yamamoto-Hino M, Mizuno H, Miyawaki A. An optical marker based on the UV-induced green-to-red photoconversion of a fluorescent protein. Proc Natl Acad Sci U S A. 2002;99(20):12651–12656. doi:10.1073/pnas.202320599

66. Lambert TJ. FPbase: a community-editable fluorescent protein database. Nat Methods. 2019;16(4):277–278. doi:10.1038/s41592-019-0352-8

67. Campbell RE, Tour O, Palmer AE, et al. A monomeric red fluorescent protein. Proc Natl Acad Sci U S A. 2002;99(12):7877–7882. doi:10.1073/pnas.082243699

68. Strack RL, Strongin DE, Bhattacharyya D, et al. A noncytotoxic DsRed variant for whole-cell labeling. Nat Methods. 2008;5(11):955–957. doi:10.1038/nmeth.1264

69. Chustecki JM, Schneider AQ, Faber MH, Christensen AC. Running on empty: Mitochondria without mtDNA exhibit differential motility and connectivity. bioRxiv. Preprint posted online August 28, 2024:2024.08.27.609985. doi:10.1101/2024.08.27.609985

70. Keren L, Zackay O, Lotan-Pompan M, et al. Promoters maintain their relative activity levels under different growth conditions. Mol Syst Biol. 2013;9(1):701. doi:10.1038/msb.2013.59

71. Seel A, Padovani F, Mayer M, et al. Regulation with cell size ensures mitochondrial DNA homeostasis during cell growth. Nat Struct Mol Biol. Published online September 7, 2023. doi:10.1038/s41594-023-01091-8

72. Padovani F, Čavka I, Neves ARR, et al. SpotMAX: a generalist framework for multidimensional automatic spot detection and quantification. bioRxiv. Preprint posted online October 23, 2024:2024.10.22.619610. doi:10.1101/2024.10.22.619610

73. Osman C, Noriega TR, Okreglak V, Fung JC, Walter P. Integrity of the yeast mitochondrial genome, but not its distribution and inheritance, relies on mitochondrial fission and fusion. Proc Natl Acad Sci U S A. 2015;112(9):E947–956. doi:10.1073/pnas.1501737112

74. Zecchini V, Paupe V, Herranz-Montoya I, et al. Fumarate induces vesicular release of mtDNA to drive innate immunity. Nature. 2023;615(7952):7952. doi:10.1038/s41586-023-05770-w

75. Sen A, Kallabis S, Gaedke F, et al. Mitochondrial membrane proteins and VPS35 orchestrate selective removal of mtDNA. Nat Commun. 2022;13(1):6704. doi:10.1038/s41467-022-34205-9

76. Al Khatib I, Deng J, Lei Y, et al. Activation of the cGAS-STING innate immune response in cells with deficient mitochondrial topoisomerase TOP1MT. Hum Mol Genet. Published online May 2, 2023:ddad062. doi:10.1093/hmg/ddad062

77. Zaman M, Sharma G, Almutawa W, et al. The MFN2 Q367H variant from a patient with lateonset distal myopathy reveals a novel pathomechanism connected to mtDNA-mediated inflammation. Life Sci Alliance. 2025;Accepted:2024.06.20.24309123. doi:10.1101/2024.06.20.24309123

78. VanPortfliet JJ, Chute C, Lei Y, Shutt TE, West AP. Mitochondrial DNA release and sensing in innate immune responses. Hum Mol Genet. 2024;33(R1):R80–R91. doi:10.1093/hmg/ddae031

79. Montava-Garriga L, Singh F, Ball G, Ganley IG. Semi-automated quantitation of mitophagy in cells and tissues. Mech Ageing Dev. 2020;185:111196. doi:10.1016/j.mad.2019.111196

80. Valente AJ, Maddalena LA, Robb EL, Moradi F, Stuart JA. A simple ImageJ macro tool for analyzing mitochondrial network morphology in mammalian cell culture. Acta Histochem. 2017;119(3):315–326. doi:10.1016/j.acthis.2017.03.001

81. Jumper J, Evans R, Pritzel A, et al. Highly accurate protein structure prediction with AlphaFold. Nature. 2021;596(7873):7873. doi:10.1038/s41586-021-03819-2

82. Mirdita M, Schütze K, Moriwaki Y, Heo L, Ovchinnikov S, Steinegger M. ColabFold: making protein folding accessible to all. Nat Methods. 2022;19(6):6. doi:10.1038/s41592-022-01488-1

83. Pettersen EF, Goddard TD, Huang CC. UCSF Chimera—a visualization system for exploratory research and analysis. J Comput Chem. 2004;25:1605–1612.

84. van Zundert GCP, Rodrigues JPGLM, Trellet M, et al. The HADDOCK2.2 Web Server: User-Friendly Integrative Modeling of Biomolecular Complexes. J Mol Biol. 2016;428(4):720–725. doi:10.1016/j.jmb.2015.09.014

85. Choi WS, Garcia-Diaz M. A minimal motif for sequence recognition by mitochondrial transcription factor A (TFAM). Nucleic Acids Res. 2022;50(1):322–332. doi:10.1093/nar/gkab1230

86. Rosselli-Murai LK, Sforça ML, Sassonia RC, et al. Structural characterization of the H-NS protein from Xylella fastidiosa and its interaction with DNA. Arch Biochem Biophys. 2012;526(1):22–28. doi:10.1016/j.abb.2012.06.007

87. Bolte S, Cordelières FP. A guided tour into subcellular colocalization analysis in light microscopy. J Microsc. 2006;224(Pt 3):213–232. doi:10.1111/j.1365-2818.2006.01706.x

88. Padovani F, Mairhörmann B, Falter-Braun P, Lengefeld J, Schmoller KM. Segmentation, tracking and cell cycle analysis of live-cell imaging data with Cell-ACDC. BMC Biol. 2022;20(1):174. doi:10.1186/s12915-022-01372-6

89. Linkert M, Rueden CT, Allan C, et al. Metadata matters: access to image data in the real world. J Cell Biol. 2010;189(5):777–782. doi:10.1083/jcb.201004104

90. Dietler N, Minder M, Gligorovski V, et al. A convolutional neural network segments yeast microscopy images with high accuracy. Nat Commun. 2020;11(1):5723. doi:10.1038/s41467-020-19557-4

91. Sato Y, Nakajima S, Shiraga N, et al. Three-dimensional multi-scale line filter for segmentation and visualization of curvilinear structures in medical images. Med Image Anal. 1998;2(2):143–168. doi:10.1016/s1361-8415(98)80009-1

92. Yen JC, Chang FJ, Chang S. A new criterion for automatic multilevel thresholding. IEEE Trans Image Process. 1995;4(3):370–378. doi:10.1109/83.366472

93. Li CH, Lee CK. Minimum cross entropy thresholding. Pattern Recognit. 1993;26(4):617–625. doi:10.1016/0031-3203(93)90115-D

