## Supplemental Information for "A novel genetic fluorescent reporter to visualize mitochondrial nucleoids"

**Supplemental Figure 1:** Nucleotide and protein sequences of mt-HI-NESS constructs. The text is colour coded according to the protein domains as follows: green indicates the double Cox8a mitochondrial targeting sequence; black indicates the linker sequence; red indicates the fluorescent protein open reading frame; blue indicates the H-NS DNA binding domain; yellow indicates the HA tag. Bolded protein residues in the protein sequences indicate predicted cleavage sites of the mitochondrial targeting sequence.

>**Mt-Kaede-HI-NESS cDNA**

ATGTCCGTCCTGACGCCGCTGCTGCTGCGGGGCTTGACAGGCTCGGCCCGGCGGCTCCCAGTGCCGCGCGCCAAGATCCATTCGTTGGGGGATCTGTCCGTCCTGACGCCGCTGCTGCTGCGGGGCTTGACAGGCTCGGCCCGGCGGCTCCCAGTGCCGCGCGCCAAGATCCATTCGTTGGGGGATCCACCGGTCGCCACCatgagtctgattaaaccagaaatgaagatcaagctgcttatggaaggcaatgtaaacgggcaccagtttgttattgagggagatggaaaaggccatccttttgagggaaaacagagtatggaccttgtagtcaaagaaggcgcacctctcccttttgcctacgatatcttgacaacagcattccattatggtaacagggtttttgctaaatacccagaccatataccagactacttcaagcagtcgtttcccaaagggttttcttgggagcgaagcctgatgttcgaggacgggggcgtttgcatcgctacaaatgacataacactgaaaggagacactttttttaacaaagttcgatttgatggcgtaaactttcccccaaatggtcctgttatgcagaagaagactctgaaatgggaggcatccactgagaaaatgtatttgcgtgatggagtgttgacgggcgatattaccatggctctgctgcttaaaggagatgtccattaccgatgtgacttcagaactacttacaaatctaggcaggagggtgtcaagttgccaggatatcactttgtcgatcactgcatcagcatattgaggcatgacaaagactacaacgaggttaagctgtatgagcatgctgttgcccattctggattgccggacaacgtcaagGctgccgttaaatctggcaccaaagctaaacgtgctcagcgtccggcaaaatatagctacgttgacgaaaacggcgaaactaaaacctggactggccaaggccgtactccagctgtaatcaaaaaagcaatggatgagcaaggtaaatccctcgacgatttcctgatcaagcaaTACCCATACGATGTTCCAGATTACGCTtaa

**>Mt-Kaede-HI-NESS Protein**

MSVLTPLLLRGLTGSARRLPVPRAKIHSLGDLSVLTPLLLRGLTGSARRLPVPRAKIHSLGDPPVATMSLIKPEMKIKLLMEGNVNGHQFVIEGDGKGHPFEGKQSMDLVVKEGAPLPFAYDILTTAFHYGNRVFAKYPDHIPDYFKQSFPKGFSWERSLMFEDGGVCIATNDITLKGDTFFNKVRFDGVNFPPNGPVMQKKTLKWEASTEKMYLRDGVLTGDITMALLLKGDVHYRCDFRTTYKSRQEGVKLPGYHFVDHCISILRHDKDYNEVKLYEHAVAHSGLPDNVKAAVKSGTKAKRAQRPAKYSYVDENGETKTWTGQGRTPAVIKKAMDEQGKSLDDFLIKQYPYDVPDYA*

**>Mt-TagRFP-HI-NESS cDNA**

ATGTCCGTCCTGACGCCGCTGCTGCTGCGGGGCTTGACAGGCTCGGCCCGGCGGCTCCCAGTGCCGCGCGCCAAGATCCATTCGTTGGGGGATCTGTCCGTCCTGACGCCGCTGCTGCTGCGGGGCTTGACAGGCTCGGCCCGGCGGCTCCCAGTGCCGCGCGCCAAGATCCATTCGTTGGGGGATCCACCGGTCGCCACCATGAGCGAGCTGATTAAGGAGAACATGCACATGAAGCTGTACATGGAGGGCACCGTGAACAACCACCACTTCAAGTGCACATCCGAGGGCGAAGGCAAGCCCTACGAGGGCACCCAGACCATGAGAATCAAGGTGGTCGAGGGCGGCCCTCTCCCCTTCGCCTTCGACATCCTGGCTACCAGCTTCATGTACGGCAGCAGAACCTTCATCAACCACACCCAGGGCATCCCCGACTTCTTTAAGCAGTCCTTCCCTGAGGGCTTCACATGGGAGAGAGTCACCACATACGAAGACGGGGGCGTGCTGACCGCTACCCAGGACACCAGCCTCCAGGACGGCTGCCTCATCTACAACGTCAAGATCAGAGGGGTGAACTTCCCATCCAACGGCCCTGTGATGCAGAAGAAAACACTCGGCTGGGAGGCCAACACCGAGATGCTGTACCCCGCTGACGGCGGCCTGGAAGGCAGAAGCGACATGGCCCTGAAGCTCGTGGGCGGGGGCCACCTGATCTGCAACTTCAAGACCACATACAGATCCAAGAAACCCGCTAAGAACCTCAAGATGCCCGGCGTCTACTATGTGGACCACAGACTGGAAAGAATCAAGGAGGCCGACAAAGAGACCTACGTCGAGCAGCACGAGGTGGCTGTGGCCAGATACTGCGACCTCCCTAGCAAACTGGGGCACAAGGctgccgttaaatctggcaccaaagctaaacgtgctcagcgtccggcaaaatatagctacgttgacgaaaacggcgaaactaaaacctggactggccaaggccgtactccagctgtaatcaaaaaagcaatggatgagcaaggtaaatccctcgacgatttcctgatcaagcaaTACCCATACGATGTTCCAGATTACGCTtaa

**>Mt-TagRFP-HI-NESS Protein**

MSVLTPLLLRGLTGSARRLPVPRAKIHSLGDLSVLTPLLLRG**L**TGSARRLPVPRA**K**IHSLGDPPVATMSELIKENMHMKLYMEGTVNNHHFKCTSEGEGKPYEGTQTMRIKVVEGGPLPFAFDILATSFMYGSRTFINHTQGIPDFFKQSFPEGFTWERVTTYEDGGVLTATQDTSLQDGCLIYNVKIRGVNFPSNGPVMQKKTLGWEANTEMLYPADGGLEGRSDMALKLVGGGHLICNFKTTYRSKKPAKNLKMPGVYYVDHRLERIKEADKETYVEQHEVAVARYCDLPSKLGHKAAVKSGTKAKRAQRPAKYSYVDENGETKTWTGQGRTPAVIKKAMDEQGKSLDDFLIKQYPYDVPDYA*

**>Mt-KillerRed-HI-NESS cDNA**

ATGTCCGTCCTGACGCCGCTGCTGCTGCGGGGCTTGACAGGCTCGGCCCGGCGGCTCCCAGTGCCGCGCGCCAAGATCCATTCGTTGGGGGATCTGTCCGTCCTGACGCCGCTGCTGCTGCGGGGCTTGACAGGCTCGGCCCGGCGGCTCCCAGTGCCGCGCGCCAAGATCCATTCGTTGGGGGATCCACCGGTCGCCACCATGGGTTCAGAGGGCGGCCCCGCCCTGTTCCAGAGCGACATGACCTTCAAAATCTTCATCGACGGCGAGGTGAACGGCCAGAAGTTCACCATCGTGGCCGACGGCAGCAGCAAGTTCCCCCACGGCGACTTCAACGTGCACGCCGTGTGCGAGACCGGCAAGCTGCCCATGAGCTGGAAGCCCATCTGCCACCTGATCCAGTACGGCGAGCCCTTCTTCGCCCGCTACCCCGACGGCATCAGCCATTTCGCCCAGGAGTGCTTCCCCGAGGGCCTGAGCATCGACCGCACCGTGCGCTTCGAGAACGACGGCACCATGACCAGCCACcACACCTACGAGCTGGACGACACCTGCGTGGTGAGCCGCATCACCGTGAACTGCGACGGCTTCCAGCCCGACGGCCCCATCATGCGCGACCAGCTGGTGGACATCCTGCCCAACGAGACCCACATGTTCCCCCACGGCCCCAACGCCGTGCGCCAGCTGGCCTTCATCGGCTTCACCACCGCCGACGGCGGCCTGATGATGGGCCACTTCGACAGCAAGATGACCTTCAACGGCAGCCGCGCCATCGAGATCCCCGGCCCACACTTCGTGACCATCATCACCAAGCAGATGAGGGACACCAGCGACAAGCGCGACCACGTGTGCCAGCGCGAGGTGGCCTACGCCCACAGCGTGCCCCGCATCACCAGCGCCATCGGtagcgacgaggatGctgccgttaaatctggcaccaaagctaaacgtgctcagcgtccggcaaaatatagctacgttgacgaaaacggcgaaactaaaacctggactggccaaggccgtactccagctgtaatcaaaaaagcaatggatgagcaaggtaaatccctcgacgatttcctgatcaagcaaTACCCATACGATGTTCCAGATTACGCTtaa

**>Mt-KillerRed-HI-NESS Protein**

MSVLTPLLLRGLTGSARRLPVPRAKIHSLGDLSVLTPLLLRG**L**TGSARRLPVPRA**K**IHSLGDPPVATMGSEGGPALFQSDMTFKIFIDGEVNGQKFTIVADGSSKFPHGDFNVHAVCETGKLPMSWKPICHLIQYGEPFFARYPDGISHFAQECFPEGLSIDRTVRFENDGTMTSHHTYELDDTCVVSRITVNCDGFQPDGPIMRDQLVDILPNETHMFPHGPNAVRQLAFIGFTTADGGLMMGHFDSKMTFNGSRAIEIPGPHFVTIITKQMRDTSDKRDHVCQREVAYAHSVPRITSAIGSDEDAAVKSGTKAKRAQRPAKYSYVDENGETKTWTGQGRTPAVIKKAMDEQGKSLDDFLIKQYPYDVPDYA*

**>Mt-mRFP1-HI-NESS cDNA**

ATGTCCGTCCTGACGCCGCTGCTGCTGCGGGGCTTGACAGGCTCGGCCCGGCGGCTCCCAGTGCCGCGCGCCAAGATCCATTCGTTGGGGGATCTGTCCGTCCTGACGCCGCTGCTGCTGCGGGGCTTGACAGGCTCGGCCCGGCGGCTCCCAGTGCCGCGCGCCAAGATCCATTCGTTGGGGGATCCACCGGTCGCCACCATGGCCTCCTCCGAGGACGTCATCAAGGAGTTCATGCGCTTCAAGGTGCGCATGGAGGGCTCCGTGAACGGCCACGAGTTCGAGATCGAGGGCGAGGGCGAGGGCCGCCCCTACGAGGGCACCCAGACCGCCAAGCTGAAGGTGACCAAGGGCGGCCCCCTGCCCTTCGCCTGGGACATCCTGTCCCCTCAGTTCCAGTACGGCTCCAAGGCCTACGTGAAGCACCCCGCCGACATCCCCGACTACTTGAAGCTGTCCTTCCCCGAGGGCTTCAAGTGGGAGCGCGTGATGAACTTCGAGGACGGCGGCGTGGTGACCGTGACCCAGGACTCCTCCCTGCAGGACGGCGAGTTCATCTACAAGGTGAAGCTGCGCGGCACCAACTTCCCCTCCGACGGCCCCGTAATGCAGAAGAAGACCATGGGCTGGGAGGCCTCCACCGAGCGGATGTACCCCGAGGACGGCGCCCTGAAGGGCGAGATCAAGATGAGGCTGAAGCTGAAGGACGGCGGCCACTACGACGCCGAGGTCAAGACCACCTACATGGCCAAGAAGCCCGTGCAGCTGCCCGGCGCCTACAAGACCGACATCAAGCTGGACATCACCTCCCACAACGAGGACTACACCATCGTGGAACAGTACGAGCGCGCCGAGGGCCGCCACTCCACCGGCGCCGCTGCCGTTAAATCTGGCACCAAAGCTAAACGTGCTCAGCGTCCGGCAAAATATAGCTACGTTGACGAAAACGGCGAAACTAAAACCTGGACTGGCCAAGGCCGTACTCCAGCTGTAATCAAAAAAGCAATGGATGAGCAAGGTAAATCCCTCGACGATTTCCTGATCAAGCAATACCCATACGATGTTCCAGATTACGCTTAA

**>Mt-mRFP1-HI-NESS Protein**

MSVLTPLLLRGLTGSARRLPVPRAKIHSLGDLSVLTPLLLRG**L**TGSARRLPVPRAKIHSLGDPPVATMASSEDVIKEFMRFKVRMEGSVNGHEFEIEGEGEGRPYEGTQTAKLKVTKGGPLPFAWDILSPQFQYGSKAYVKHPADIPDYLKLSFPEGFKWERVMNFEDGGVVTVTQDSSLQDGEFIYKVKLRGTNFPSDGPVMQKKTMGWEASTERMYPEDGALKGEIKMRLKLKDGGHYDAEVKTTYMAKKPVQLPGAYKTDIKLDITSHNEDYTIVEQYERAEGRHSTGAAAVKSGTKAKRAQRPAKYSYVDENGETKTWTGQGRTPAVIKKAMDEQGKSLDDFLIKQYPYDVPDYA*

**>Mt-Dimer1-HI-NESS cDNA**

ATGTCCGTCCTGACGCCGCTGCTGCTGCGGGGCTTGACAGGCTCGGCCCGGCGGCTCCCAGTGCCGCGCGCCAAGATCCATTCGTTGGGGGATCTGTCCGTCCTGACGCCGCTGCTGCTGCGGGGCTTGACAGGCTCGGCCCGGCGGCTCCCAGTGCCGCGCGCCAAGATCCATTCGTTGGGGGATCCACCGGTCGCCACCATGGTGGCCTCCTCCGAGGACGTCATCAAAGAGTTCATGCGCTTCAAGGTGCGCATGGAGGGCTCCGTGAACGGCCACGAGTTCGAGATCGAGGGCGAGGGCGAGGGCCGCCCCTACGAGGGCACCCAGACCGCCAAGCTGAAGGTGACCAAGGGCGGCCCCCTGCCCTTCGCCTGGGACATCCTGTCCCCCCAGTTCCAGTACGGCTCCAAGGTGTACGTGAAGCACCCCGCCGACATCCCCGACTACAAGAAGCTGTCCTTCCCCGAGGGCTTCAAGTGGGAGCGCGTGATGAACTTCGAGGACGGCGGCGTGGTGACCGTGACCCAGGACTCCTCCCTGCAGGACGGCTGCTTTATCTACAAGGTGAAGTTCCGCGGCGTGAACTTCCCCTCCGACGGCCCCGTAATGCAGAAGAAGACCATGGGCTGGGAGGCCTCCACCGAGCGCCTGTACCCCCGCGACGGCGTGCTGAAGGGCGAGATCCACCAGGCCCTGAAGCTGAAGGACGGCGGCCACTACCTGGTGGAGTTCAAGACCATCTACATGGCCAAGAAGCCCGTGCAGCTGCCCGGCTACTACTACGTGGACACCAAGCTGGACATCACCTCCCACAACGAGGACTACACCATCGTGGAACAGTACGAGCGCGCCGAGGGCCGCCACCACCTGTTCCTGGCTGCCGTTAAATCTGGCACCAAAGCTAAACGTGCTCAGCGTCCGGCAAAATATAGCTACGTTGACGAAAACGGCGAAACTAAAACCTGGACTGGCCAAGGCCGTACTCCAGCTGTAATCAAAAAAGCAATGGATGAGCAAGGTAAATCCCTCGACGATTTCCTGATCAAGCAATACCCATACGATGTTCCAGATTACGCTTAA

**>Mt-Dimer1-HI-NESS Protein**

MSVLTPLLLRGLTGSARRLPVPRAKIHSLGDLSVLTPLLLRG**L**TGSARRLPVPRAKIHSLGDPPVATMVASSEDVIKEFMRFKVRMEGSVNGHEFEIEGEGEGRPYEGTQTAKLKVTKGGPLPFAWDILSPQFQYGSKVYVKHPADIPDYKKLSFPEGFKWERVMNFEDGGVVTVTQDSSLQDGCFIYKVKFRGVNFPSDGPVMQKKTMGWEASTERLYPRDGVLKGEIHQALKLKDGGHYLVEFKTIYMAKKPVQLPGYYYVDTKLDITSHNEDYTIVEQYERAEGRHHLFLAAVKSGTKAKRAQRPAKYSYVDENGETKTWTGQGRTPAVIKKAMDEQGKSLDDFLIKQYPYDVPDYA*

**>Mt-DsRED-Express2-HI-NESS cDNA**

ATGTCCGTCCTGACGCCGCTGCTGCTGCGGGGCTTGACAGGCTCGGCCCGGCGGCTCCCAGTGCCGCGCGCCAAGATCCATTCGTTGGGGGATCTGTCCGTCCTGACGCCGCTGCTGCTGCGGGGCTTGACAGGCTCGGCCCGGCGGCTCCCAGTGCCGCGCGCCAAGATCCATTCGTTGGGGGATCCACCGGTCGCCACCATGGATAGCACTGAGAACGTCATCAAGCCCTTCATGCGCTTCAAGGTGCACATGGAGGGCTCCGTGAACGGCCACGAGTTCGAGATCGAGGGCGAGGGCGAGGGCAAGCCCTACGAGGGCACCCAGACCGCCAAGCTGCAGGTGACCAAGGGCGGCCCCCTGCCCTTCGCCTGGGACATCCTGTCCCCCCAGTTCCAGTACGGCTCCAAGGTGTACGTGAAGCACCCCGCCGACATCCCCGACTACAAGAAGCTGTCCTTCCCCGAGGGCTTCAAGTGGGAGCGCGTGATGAACTTCGAGGACGGCGGCGTGGTGACCGTGACCCAGGACTCCTCCCTGCAGGACGGCACCTTCATCTACCACGTGAAGTTCATCGGCGTGAACTTCCCCTCCGACGGCCCCGTAATGCAGAAGAAGACTCTGGGCTGGGAGCCCTCCACCGAGCGCCTGTACCCCCGCGACGGCGTGCTGAAGGGCGAGATCCACAAGGCGCTGAAGCTGAAGGGCGGCGGCCACTACCTGGTGGAGTTCAAGTCAATCTACATGGCCAAGAAGCCCGTGAAGCTGCCCGGCTACTACTACGTGGACTCCAAGCTGGACATCACCTCCCACAACGAGGACTACACCGTGGTGGAGCAGTACGAGCGCGCCGAGGCCCGCCACCACCTGTTCCAGGCTGCCGTTAAATCTGGCACCAAAGCTAAACGTGCTCAGCGTCCGGCAAAATATAGCTACGTTGACGAAAACGGCGAAACTAAAACCTGGACTGGCCAAGGCCGTACTCCAGCTGTAATCAAAAAAGCAATGGATGAGCAAGGTAAATCCCTCGACGATTTCCTGATCAAGCAATACCCATACGATGTTCCAGATTACGCTTAA

**>Mt-DsRED-Express2-HI-NESS Protein**

MSVLTPLLLRGLTGSARRLPVPRAKIHSLGDLSVLTPLLLRG**L**TGSARRLPVPRAKIHSLGDPPVATMDSTENVIKPFMRFKVHMEGSVNGHEFEIEGEGEGKPYEGTQTAKLQVTKGGPLPFAWDILSPQFQYGSKVYVKHPADIPDYKKLSFPEGFKWERVMNFEDGGVVTVTQDSSLQDGTFIYHVKFIGVNFPSDGPVMQKKTLGWEPSTERLYPRDGVLKGEIHKALKLKGGGHYLVEFKSIYMAKKPVKLPGYYYVDSKLDITSHNEDYTVVEQYERAEARHHLFQAAVKSGTKAKRAQRPAKYSYVDENGETKTWTGQGRTPAVIKKAMDEQGKSLDDFLIKQYPYDVPDYA*

**>mt-TagRFP-N/C-HI-NESS cDNA**

ATGTCCGTCCTGACGCCGCTGCTGCTGCGGGGCTTGACAGGCTCGGCCCGGCGGCTCCCAGTGCCGCGCGCCAAGATCCATTCGTTGGGGGATCTGTCCGTCCTGACGCCGCTGCTGCTGCGGGGCTTGACAGGCTCGGCCCGGCGGCTCCCAGTGCCGCGCGCCAAGgcCgcTgtGaaGtcCggAacAaaGgcCaaGAgAgcCcaAAgAccCgcCaaGtaCTCTtaTgtGgaTgaGaaTggAgaGacCaaGacAtggacCggTcaGggTAgAacAccTgcAgtGatTaaGaaGgcTatggaCgaAcaGggAaaGAGcctGgaTgaCttTctCatTaaAcaGATCCATTCGTTGGGGGATCCACCGGTCGCCACCATGAGCGAGCTGATTAAGGAGAACATGCACATGAAGCTGTACATGGAGGGCACCGTGAACAACCACCACTTCAAGTGCACATCCGAGGGCGAAGGCAAGCCCTACGAGGGCACCCAGACCATGAGAATCAAGGTGGTCGAGGGCGGCCCTCTCCCCTTCGCCTTCGACATCCTGGCTACCAGCTTCATGTACGGCAGCAGAACCTTCATCAACCACACCCAGGGCATCCCCGACTTCTTTAAGCAGTCCTTCCCTGAGGGCTTCACATGGGAGAGAGTCACCACATACGAAGACGGGGGCGTGCTGACCGCTACCCAGGACACCAGCCTCCAGGACGGCTGCCTCATCTACAACGTCAAGATCAGAGGGGTGAACTTCCCATCCAACGGCCCTGTGATGCAGAAGAAAACACTCGGCTGGGAGGCCAACACCGAGATGCTGTACCCCGCTGACGGCGGCCTGGAAGGCAGAAGCGACATGGCCCTGAAGCTCGTGGGCGGGGGCCACCTGATCTGCAACTTCAAGACCACATACAGATCCAAGAAACCCGCTAAGAACCTCAAGATGCCCGGCGTCTACTATGTGGACCACAGACTGGAAAGAATCAAGGAGGCCGACAAAGAGACCTACGTCGAGCAGCACGAGGTGGCTGTGGCCAGATACTGCGACCTCCCTAGCAAACTGGGGCACAAGgctgccgttaaatctggcaccaaagctaaacgtgctcagcgtccggcaaaatatagctacgttgacgaaaacggcgaaactaaaacctggactggccaaggccgtactccagctgtaatcaaaaaagcaatggatgagcaaggtaaatccctcgacgatttcctgatcaagcaaTACCCATACGATGTTCCAGATTACGCTtaa

**>mt-TagRFP-N/C-HI-NESS Protein**

MSVLTPLLLRGLTGSARRLPVPRAKIHSLGDLSVLTPLLLRG**L**TGSARRLPVPRAKAAVKSGTKAKRAQRPAKYSYVDENGETKTWTGQGRTPAVIKKAMDEQGKSLDDFLIKQIHSLGDPPVATMSELIKENMHMKLYMEGTVNNHHFKCTSEGEGKPYEGTQTMRIKVVEGGPLPFAFDILATSFMYGSRTFINHTQGIPDFFKQSFPEGFTWERVTTYEDGGVLTATQDTSLQDGCLIYNVKIRGVNFPSNGPVMQKKTLGWEANTEMLYPADGGLEGRSDMALKLVGGGHLICNFKTTYRSKKPAKNLKMPGVYYVDHRLERIKEADKETYVEQHEVAVARYCDLPSKLGHKAAVKSGTKAKRAQRPAKYSYVDENGETKTWTGQGRTPAVIKKAMDEQGKSLDDFLIKQYPYDVPDYA*

**Supplementary Figure 2:** Vector Map of the plasmids used in this study.

**Mt-Kaede-HI-NESS**

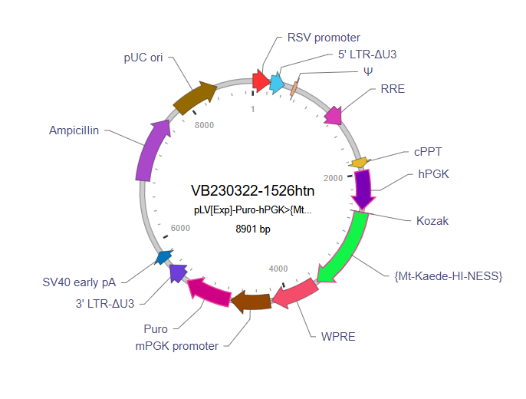

**Mt-TagRFP-HI-NESS**

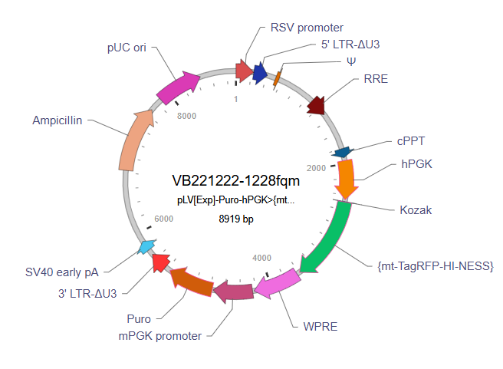

**Mt-KillerRed-HI-NESS**

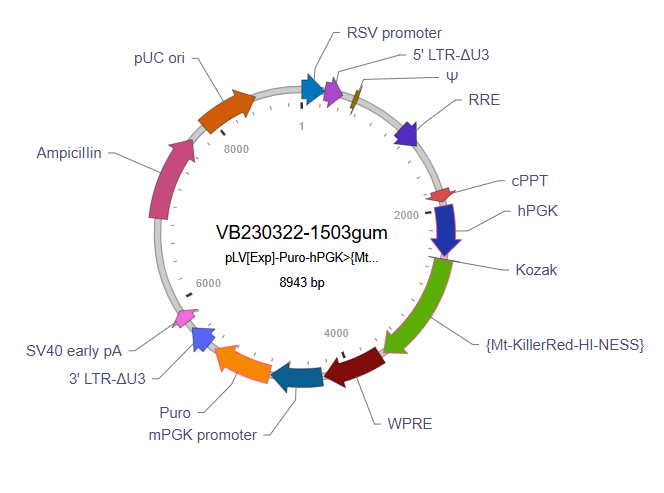

**Mt-mRFP1-HI-NESS**

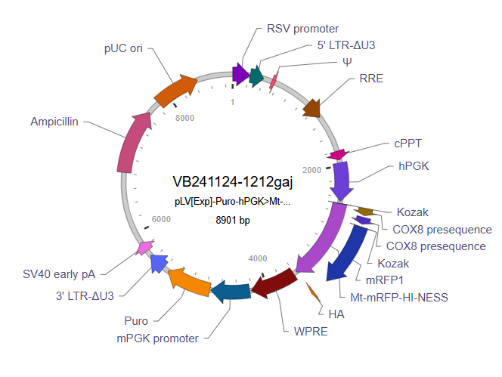

**Mt-Dimer1-HI-NESS**

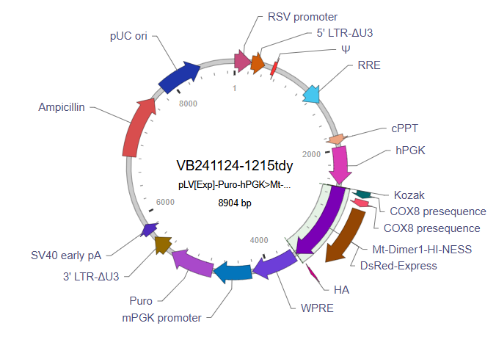

**Mt-DsRED-Express2-HI-NESS**

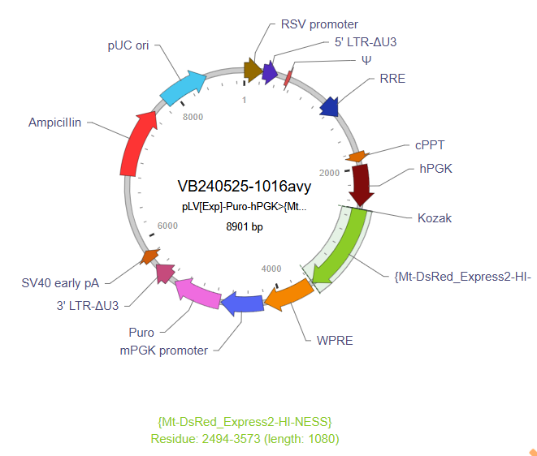

**Mt-TagRFP-N/C-HI-NESS (Double-HI-NESS)**

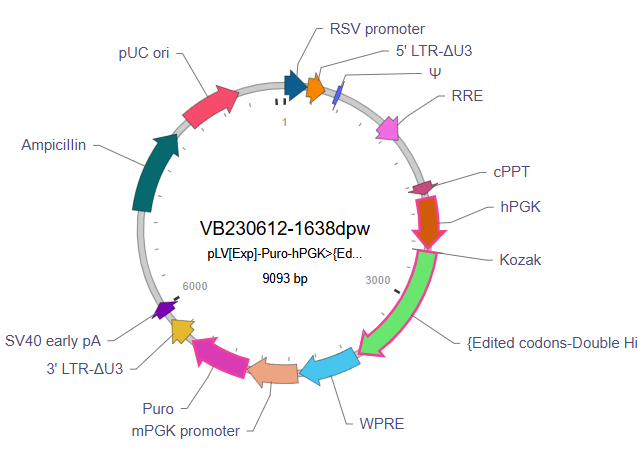

**Supplementary Figure 3:** Nucleotide sequences of the plasmids used in this study. The indicated vector IDs can be used to retrieve detailed information about the vector of vectorbuilder.com.

**> Mt-Kaede-HI-NESS - VB230322-1526htn - pLV[Exp]-Puro-hPGK>{Mt-Kaede-HI-NESS}**

AATGTAGTCTTATGCAATACTCTTGTAGTCTTGCAACATGGTAACGATGAGTTAGCAACATGCCTTACAAGGAGAGAAAAAGCACCGTGCATGCCGATTGGTGGAAGTAAGGTGGTACGATCGTGCCTTATTAGGAAGGCAACAGACGGGTCTGACATGGATTGGACGAACCACTGAATTGCCGCATTGCAGAGATATTGTATTTAAGTGCCTAGCTCGATACATAAACGGGTCTCTCTGGTTAGACCAGATCTGAGCCTGGGAGCTCTCTGGCTAACTAGGGAACCCACTGCTTAAGCCTCAATAAAGCTTGCCTTGAGTGCTTCAAGTAGTGTGTGCCCGTCTGTTGTGTGACTCTGGTAACTAGAGATCCCTCAGACCCTTTTAGTCAGTGTGGAAAATCTCTAGCAGTGGCGCCCGAACAGGGACTTGAAAGCGAAAGGGAAACCAGAGGAGCTCTCTCGACGCAGGACTCGGCTTGCTGAAGCGCGCACGGCAAGAGGCGAGGGGCGGCGACTGGTGAGTACGCCAAAAATTTTGACTAGCGGAGGCTAGAAGGAGAGAGATGGGTGCGAGAGCGTCAGTATTAAGCGGGGGAGAATTAGATCGCGATGGGAAAAAATTCGGTTAAGGCCAGGGGGAAAGAAAAAATATAAATTAAAACATATAGTATGGGCAAGCAGGGAGCTAGAACGATTCGCAGTTAATCCTGGCCTGTTAGAAACATCAGAAGGCTGTAGACAAATACTGGGACAGCTACAACCATCCCTTCAGACAGGATCAGAAGAACTTAGATCATTATATAATACAGTAGCAACCCTCTATTGTGTGCATCAAAGGATAGAGATAAAAGACACCAAGGAAGCTTTAGACAAGATAGAGGAAGAGCAAAACAAAAGTAAGACCACCGCACAGCAAGCGGCCGCTGATCTTCAGACCTGGAGGAGGAGATATGAGGGACAATTGGAGAAGTGAATTATATAAATATAAAGTAGTAAAAATTGAACCATTAGGAGTAGCACCCACCAAGGCAAAGAGAAGAGTGGTGCAGAGAGAAAAAAGAGCAGTGGGAATAGGAGCTTTGTTCCTTGGGTTCTTGGGAGCAGCAGGAAGCACTATGGGCGCAGCGTCAATGACGCTGACGGTACAGGCCAGACAATTATTGTCTGGTATAGTGCAGCAGCAGAACAATTTGCTGAGGGCTATTGAGGCGCAACAGCATCTGTTGCAACTCACAGTCTGGGGCATCAAGCAGCTCCAGGCAAGAATCCTGGCTGTGGAAAGATACCTAAAGGATCAACAGCTCCTGGGGATTTGGGGTTGCTCTGGAAAACTCATTTGCACCACTGCTGTGCCTTGGAATGCTAGTTGGAGTAATAAATCTCTGGAACAGATTTGGAATCACACGACCTGGATGGAGTGGGACAGAGAAATTAACAATTACACAAGCTTAATACACTCCTTAATTGAAGAATCGCAAAACCAGCAAGAAAAGAATGAACAAGAATTATTGGAATTAGATAAATGGGCAAGTTTGTGGAATTGGTTTAACATAACAAATTGGCTGTGGTATATAAAATTATTCATAATGATAGTAGGAGGCTTGGTAGGTTTAAGAATAGTTTTTGCTGTACTTTCTATAGTGAATAGAGTTAGGCAGGGATATTCACCATTATCGTTTCAGACCCACCTCCCAACCCCGAGGGGACCCGACAGGCCCGAAGGAATAGAAGAAGAAGGTGGAGAGAGAGACAGAGACAGATCCATTCGATTAGTGAACGGATCTCGACGGTATCGCTAGCTTTTAAAAGAAAAGGGGGGATTGGGGGGTACAGTGCAGGGGAAAGAATAGTAGACATAATAGCAACAGACATACAAACTAAAGAATTACAAAAACAAATTACAAAAATTCAAAATTTTACTAGTGATTATCGGATCAACTTTGTATAGAAAAGTTGGGGTTGCGCCTTTTCCAAGGCAGCCCTGGGTTTGCGCAGGGACGCGGCTGCTCTGGGCGTGGTTCCGGGAAACGCAGCGGCGCCGACCCTGGGTCTCGCACATTCTTCACGTCCGTTCGCAGCGTCACCCGGATCTTCGCCGCTACCCTTGTGGGCCCCCCGGCGACGCTTCCTGCTCCGCCCCTAAGTCGGGAAGGTTCCTTGCGGTTCGCGGCGTGCCGGACGTGACAAACGGAAGCCGCACGTCTCACTAGTACCCTCGCAGACGGACAGCGCCAGGGAGCAATGGCAGCGCGCCGACCGCGATGGGCTGTGGCCAATAGCGGCTGCTCAGCAGGGCGCGCCGAGAGCAGCGGCCGGGAAGGGGCGGTGCGGGAGGCGGGGTGTGGGGCGGTAGTGTGGGCCCTGTTCCTGCCCGCGCGGTGTTCCGCATTCTGCAAGCCTCCGGAGCGCACGTCGGCAGTCGGCTCCCTCGTTGACCGAATCACCGACCTCTCTCCCCAGGCAAGTTTGTACAAAAAAGCAGGCTGCCACCATGTCCGTCCTGACGCCGCTGCTGCTGCGGGGCTTGACAGGCTCGGCCCGGCGGCTCCCAGTGCCGCGCGCCAAGATCCATTCGTTGGGGGATCTGTCCGTCCTGACGCCGCTGCTGCTGCGGGGCTTGACAGGCTCGGCCCGGCGGCTCCCAGTGCCGCGCGCCAAGATCCATTCGTTGGGGGATCCACCGGTCGCCACCATGAGTCTGATTAAACCAGAAATGAAGATCAAGCTGCTTATGGAAGGCAATGTAAACGGGCACCAGTTTGTTATTGAGGGAGATGGAAAAGGCCATCCTTTTGAGGGAAAACAGAGTATGGACCTTGTAGTCAAAGAAGGCGCACCTCTCCCTTTTGCCTACGATATCTTGACAACAGCATTCCATTATGGTAACAGGGTTTTTGCTAAATACCCAGACCATATACCAGACTACTTCAAGCAGTCGTTTCCCAAAGGGTTTTCTTGGGAGCGAAGCCTGATGTTCGAGGACGGGGGCGTTTGCATCGCTACAAATGACATAACACTGAAAGGAGACACTTTTTTTAACAAAGTTCGATTTGATGGCGTAAACTTTCCCCCAAATGGTCCTGTTATGCAGAAGAAGACTCTGAAATGGGAGGCATCCACTGAGAAAATGTATTTGCGTGATGGAGTGTTGACGGGCGATATTACCATGGCTCTGCTGCTTAAAGGAGATGTCCATTACCGATGTGACTTCAGAACTACTTACAAATCTAGGCAGGAGGGTGTCAAGTTGCCAGGATATCACTTTGTCGATCACTGCATCAGCATATTGAGGCATGACAAAGACTACAACGAGGTTAAGCTGTATGAGCATGCTGTTGCCCATTCTGGATTGCCGGACAACGTCAAGGCTGCCGTTAAATCTGGCACCAAAGCTAAACGTGCTCAGCGTCCGGCAAAATATAGCTACGTTGACGAAAACGGCGAAACTAAAACCTGGACTGGCCAAGGCCGTACTCCAGCTGTAATCAAAAAAGCAATGGATGAGCAAGGTAAATCCCTCGACGATTTCCTGATCAAGCAATACCCATACGATGTTCCAGATTACGCTTAAACCCAGCTTTCTTGTACAAAGTGGTGATAATCGAATTCCGATAATCAACCTCTGGATTACAAAATTTGTGAAAGATTGACTGGTATTCTTAACTATGTTGCTCCTTTTACGCTATGTGGATACGCTGCTTTAATGCCTTTGTATCATGCTATTGCTTCCCGTATGGCTTTCATTTTCTCCTCCTTGTATAAATCCTGGTTGCTGTCTCTTTATGAGGAGTTGTGGCCCGTTGTCAGGCAACGTGGCGTGGTGTGCACTGTGTTTGCTGACGCAACCCCCACTGGTTGGGGCATTGCCACCACCTGTCAGCTCCTTTCCGGGACTTTCGCTTTCCCCCTCCCTATTGCCACGGCGGAACTCATCGCCGCCTGCCTTGCCCGCTGCTGGACAGGGGCTCGGCTGTTGGGCACTGACAATTCCGTGGTGTTGTCGGGGAAGCTGACGTCCTTTCCATGGCTGCTCGCCTGTGTTGCCACCTGGATTCTGCGCGGGACGTCCTTCTGCTACGTCCCTTCGGCCCTCAATCCAGCGGACCTTCCTTCCCGCGGCCTGCTGCCGGCTCTGCGGCCTCTTCCGCGTCTTCGCCTTCGCCCTCAGACGAGTCGGATCTCCCTTTGGGCCGCCTCCCCGCATCGGGAATTCCCGCGGTTCGAATTCTACCGGGTAGGGGAGGCGCTTTTCCCAAGGCAGTCTGGAGCATGCGCTTTAGCAGCCCCGCTGGGCACTTGGCGCTACACAAGTGGCCTCTGGCCTCGCACACATTCCACATCCACCGGTAGGCGCCAACCGGCTCCGTTCTTTGGTGGCCCCTTCGCGCCACCTTCTACTCCTCCCCTAGTCAGGAAGTTCCCCCCCGCCCCGCAGCTCGCGTCGTGCAGGACGTGACAAATGGAAGTAGCACGTCTCACTAGTCTCGTGCAGATGGACAGCACCGCTGAGCAATGGAAGCGGGTAGGCCTTTGGGGCAGCGGCCAATAGCAGCTTTGCTCCTTCGCTTTCTGGGCTCAGAGGCTGGGAAGGGGTGGGTCCGGGGGCGGGCTCAGGGGCGGGCTCAGGGGCGGGGCGGGCGCCCGAAGGTCCTCCGGAGGCCCGGCATTCTGCACGCTTCAAAAGCGCACGTCTGCCGCGCTGTTCTCCTCTTCCTCATCTCCGGGCCTTTCGACCTCACGTGGCCACCATGACCGAGTACAAGCCCACGGTGCGCCTCGCCACCCGCGACGACGTCCCCAGGGCCGTACGCACCCTCGCCGCCGCGTTCGCCGACTACCCCGCCACGCGCCACACCGTCGATCCGGACCGCCACATCGAGCGGGTCACCGAGCTGCAAGAACTCTTCCTCACGCGCGTCGGGCTCGACATCGGCAAGGTGTGGGTCGCGGACGACGGCGCCGCGGTGGCGGTCTGGACCACGCCGGAGAGCGTCGAAGCGGGGGCGGTGTTCGCCGAGATCGGCCCGCGCATGGCCGAGTTGAGCGGTTCCCGGCTGGCCGCGCAGCAACAGATGGAAGGCCTCCTGGCGCCGCACCGGCCCAAGGAGCCCGCGTGGTTCCTGGCCACCGTCGGCGTCTCGCCCGACCACCAGGGCAAGGGTCTGGGCAGCGCCGTCGTGCTCCCCGGAGTGGAGGCGGCCGAGCGCGCCGGGGTGCCCGCCTTCCTGGAGACCTCCGCGCCCCGCAACCTCCCCTTCTACGAGCGGCTCGGCTTCACCGTCACCGCCGACGTCGAGGTGCCCGAAGGACCGCGCACCTGGTGCATGACCCGCAAGCCCGGTGCCTGAGGTACCTTTAAGACCAATGACTTACAAGGCAGCTGTAGATCTTAGCCACTTTTTAAAAGAAAAGGGGGGACTGGAAGGGCTAATTCACTCCCAACGAAGACAAGATCTGCTTTTTGCTTGTACTGGGTCTCTCTGGTTAGACCAGATCTGAGCCTGGGAGCTCTCTGGCTAACTAGGGAACCCACTGCTTAAGCCTCAATAAAGCTTGCCTTGAGTGCTTCAAGTAGTGTGTGCCCGTCTGTTGTGTGACTCTGGTAACTAGAGATCCCTCAGACCCTTTTAGTCAGTGTGGAAAATCTCTAGCAGTAGTAGTTCATGTCATCTTATTATTCAGTATTTATAACTTGCAAAGAAATGAATATCAGAGAGTGAGAGGAACTTGTTTATTGCAGCTTATAATGGTTACAAATAAAGCAATAGCATCACAAATTTCACAAATAAAGCATTTTTTTCACTGCATTCTAGTTGTGGTTTGTCCAAACTCATCAATGTATCTTATCATGTCTGGCTCTAGCTATCCCGCCCCTAACTCCGCCCATCCCGCCCCTAACTCCGCCCAGTTCCGCCCATTCTCCGCCCCATGGCTGACTAATTTTTTTTATTTATGCAGAGGCCGAGGCCGCCTCGGCCTCTGAGCTATTCCAGAAGTAGTGAGGAGGCTTTTTTGGAGGCCTAGGGACGTACCCAATTCGCCCTATAGTGAGTCGTATTACGCGCGCTCACTGGCCGTCGTTTTACAACGTCGTGACTGGGAAAACCCTGGCGTTACCCAACTTAATCGCCTTGCAGCACATCCCCCTTTCGCCAGCTGGCGTAATAGCGAAGAGGCCCGCACCGATCGCCCTTCCCAACAGTTGCGCAGCCTGAATGGCGAATGGGACGCGCCCTGTAGCGGCGCATTAAGCGCGGCGGGTGTGGTGGTTACGCGCAGCGTGACCGCTACACTTGCCAGCGCCCTAGCGCCCGCTCCTTTCGCTTTCTTCCCTTCCTTTCTCGCCACGTTCGCCGGCTTTCCCCGTCAAGCTCTAAATCGGGGGCTCCCTTTAGGGTTCCGATTTAGTGCTTTACGGCACCTCGACCCCAAAAAACTTGATTAGGGTGATGGTTCACGTAGTGGGCCATCGCCCTGATAGACGGTTTTTCGCCCTTTGACGTTGGAGTCCACGTTCTTTAATAGTGGACTCTTGTTCCAAACTGGAACAACACTCAACCCTATCTCGGTCTATTCTTTTGATTTATAAGGGATTTTGCCGATTTCGGCCTATTGGTTAAAAAATGAGCTGATTTAACAAAAATTTAACGCGAATTTTAACAAAATATTAACGCTTACAATTTAGGTGGCACTTTTCGGGGAAATGTGCGCGGAACCCCTATTTGTTTATTTTTCTAAATACATTCAAATATGTATCCGCTCATGAGACAATAACCCTGATAAATGCTTCAATAATATTGAAAAAGGAAGAGTATGAGTATTCAACATTTCCGTGTCGCCCTTATTCCCTTTTTTGCGGCATTTTGCCTTCCTGTTTTTGCTCACCCAGAAACGCTGGTGAAAGTAAAAGATGCTGAAGATCAGTTGGGTGCACGAGTGGGTTACATCGAACTGGATCTCAACAGCGGTAAGATCCTTGAGAGTTTTCGCCCCGAAGAACGTTTTCCAATGATGAGCACTTTTAAAGTTCTGCTATGTGGCGCGGTATTATCCCGTATTGACGCCGGGCAAGAGCAACTCGGTCGCCGCATACACTATTCTCAGAATGACTTGGTTGAGTACTCACCAGTCACAGAAAAGCATCTTACGGATGGCATGACAGTAAGAGAATTATGCAGTGCTGCCATAACCATGAGTGATAACACTGCGGCCAACTTACTTCTGACAACGATCGGAGGACCGAAGGAGCTAACCGCTTTTTTGCACAACATGGGGGATCATGTAACTCGCCTTGATCGTTGGGAACCGGAGCTGAATGAAGCCATACCAAACGACGAGCGTGACACCACGATGCCTGTAGCAATGGCAACAACGTTGCGCAAACTATTAACTGGCGAACTACTTACTCTAGCTTCCCGGCAACAATTAATAGACTGGATGGAGGCGGATAAAGTTGCAGGACCACTTCTGCGCTCGGCCCTTCCGGCTGGCTGGTTTATTGCTGATAAATCTGGAGCCGGTGAGCGTGGGTCTCGCGGTATCATTGCAGCACTGGGGCCAGATGGTAAGCCCTCCCGTATCGTAGTTATCTACACGACGGGGAGTCAGGCAACTATGGATGAACGAAATAGACAGATCGCTGAGATAGGTGCCTCACTGATTAAGCATTGGTAACTGTCAGACCAAGTTTACTCATATATACTTTAGATTGATTTAAAACTTCATTTTTAATTTAAAAGGATCTAGGTGAAGATCCTTTTTGATAATCTCATGACCAAAATCCCTTAACGTGAGTTTTCGTTCCACTGAGCGTCAGACCCCGTAGAAAAGATCAAAGGATCTTCTTGAGATCCTTTTTTTCTGCGCGTAATCTGCTGCTTGCAAACAAAAAAACCACCGCTACCAGCGGTGGTTTGTTTGCCGGATCAAGAGCTACCAACTCTTTTTCCGAAGGTAACTGGCTTCAGCAGAGCGCAGATACCAAATACTGTTCTTCTAGTGTAGCCGTAGTTAGGCCACCACTTCAAGAACTCTGTAGCACCGCCTACATACCTCGCTCTGCTAATCCTGTTACCAGTGGCTGCTGCCAGTGGCGATAAGTCGTGTCTTACCGGGTTGGACTCAAGACGATAGTTACCGGATAAGGCGCAGCGGTCGGGCTGAACGGGGGGTTCGTGCACACAGCCCAGCTTGGAGCGAACGACCTACACCGAACTGAGATACCTACAGCGTGAGCTATGAGAAAGCGCCACGCTTCCCGAAGAGAGAAAGGCGGACAGGTATCCGGTAAGCGGCAGGGTCGGAACAGGAGAGCGCACGAGGGAGCTTCCAGGGGGAAACGCCTGGTATCTTTATAGTCCTGTCGGGTTTCGCCACCTCTGACTTGAGCGTCGATTTTTGTGATGCTCGTCAGGGGGGCGGAGCCTATGGAAAAACGCCAGCAACGCGGCCTTTTTACGGTTCCTGGCCTTTTGCTGGCCTTTTGCTCACATGTTCTTTCCTGCGTTATCCCCTGATTCTGTGGATAACCGTATTACCGCCTTTGAGTGAGCTGATACCGCTCGCCGCAGCCGAACGACCGAGCGCAGCGAGTCAGTGAGCGAGGAAGCGGAAGAGCGCCCAATACGCAAACCGCCTCTCCCCGCGCGTTGGCCGATTCATTAATGCAGCTGGCACGACAGGTTTCCCGACTGGAAAGCGGGCAGTGAGCGCAACGCAATTAATGTGAGTTAGCTCACTCATTAGGCACCCCAGGCTTTACACTTTATGCTTCCGGCTCGTATGTTGTGTGGAATTGTGAGCGGATAACAATTTCACACAGGAAACAGCTATGACCATGATTACGCCAAGCGCGCAATTAACCCTCACTAAAGGGAACAAAAGCTGGAGCTGCAAGCTT

**> Mt-TagRFP-HI-NESS - VB221222-1228fqm - pLV[Exp]-Puro-hPGK>{mt-TagRFP-HI-NESS}**

AATGTAGTCTTATGCAATACTCTTGTAGTCTTGCAACATGGTAACGATGAGTTAGCAACATGCCTTACAAGGAGAGAAAAAGCACCGTGCATGCCGATTGGTGGAAGTAAGGTGGTACGATCGTGCCTTATTAGGAAGGCAACAGACGGGTCTGACATGGATTGGACGAACCACTGAATTGCCGCATTGCAGAGATATTGTATTTAAGTGCCTAGCTCGATACATAAACGGGTCTCTCTGGTTAGACCAGATCTGAGCCTGGGAGCTCTCTGGCTAACTAGGGAACCCACTGCTTAAGCCTCAATAAAGCTTGCCTTGAGTGCTTCAAGTAGTGTGTGCCCGTCTGTTGTGTGACTCTGGTAACTAGAGATCCCTCAGACCCTTTTAGTCAGTGTGGAAAATCTCTAGCAGTGGCGCCCGAACAGGGACTTGAAAGCGAAAGGGAAACCAGAGGAGCTCTCTCGACGCAGGACTCGGCTTGCTGAAGCGCGCACGGCAAGAGGCGAGGGGCGGCGACTGGTGAGTACGCCAAAAATTTTGACTAGCGGAGGCTAGAAGGAGAGAGATGGGTGCGAGAGCGTCAGTATTAAGCGGGGGAGAATTAGATCGCGATGGGAAAAAATTCGGTTAAGGCCAGGGGGAAAGAAAAAATATAAATTAAAACATATAGTATGGGCAAGCAGGGAGCTAGAACGATTCGCAGTTAATCCTGGCCTGTTAGAAACATCAGAAGGCTGTAGACAAATACTGGGACAGCTACAACCATCCCTTCAGACAGGATCAGAAGAACTTAGATCATTATATAATACAGTAGCAACCCTCTATTGTGTGCATCAAAGGATAGAGATAAAAGACACCAAGGAAGCTTTAGACAAGATAGAGGAAGAGCAAAACAAAAGTAAGACCACCGCACAGCAAGCGGCCGCTGATCTTCAGACCTGGAGGAGGAGATATGAGGGACAATTGGAGAAGTGAATTATATAAATATAAAGTAGTAAAAATTGAACCATTAGGAGTAGCACCCACCAAGGCAAAGAGAAGAGTGGTGCAGAGAGAAAAAAGAGCAGTGGGAATAGGAGCTTTGTTCCTTGGGTTCTTGGGAGCAGCAGGAAGCACTATGGGCGCAGCGTCAATGACGCTGACGGTACAGGCCAGACAATTATTGTCTGGTATAGTGCAGCAGCAGAACAATTTGCTGAGGGCTATTGAGGCGCAACAGCATCTGTTGCAACTCACAGTCTGGGGCATCAAGCAGCTCCAGGCAAGAATCCTGGCTGTGGAAAGATACCTAAAGGATCAACAGCTCCTGGGGATTTGGGGTTGCTCTGGAAAACTCATTTGCACCACTGCTGTGCCTTGGAATGCTAGTTGGAGTAATAAATCTCTGGAACAGATTTGGAATCACACGACCTGGATGGAGTGGGACAGAGAAATTAACAATTACACAAGCTTAATACACTCCTTAATTGAAGAATCGCAAAACCAGCAAGAAAAGAATGAACAAGAATTATTGGAATTAGATAAATGGGCAAGTTTGTGGAATTGGTTTAACATAACAAATTGGCTGTGGTATATAAAATTATTCATAATGATAGTAGGAGGCTTGGTAGGTTTAAGAATAGTTTTTGCTGTACTTTCTATAGTGAATAGAGTTAGGCAGGGATATTCACCATTATCGTTTCAGACCCACCTCCCAACCCCGAGGGGACCCGACAGGCCCGAAGGAATAGAAGAAGAAGGTGGAGAGAGAGACAGAGACAGATCCATTCGATTAGTGAACGGATCTCGACGGTATCGCTAGCTTTTAAAAGAAAAGGGGGGATTGGGGGGTACAGTGCAGGGGAAAGAATAGTAGACATAATAGCAACAGACATACAAACTAAAGAATTACAAAAACAAATTACAAAAATTCAAAATTTTACTAGTGATTATCGGATCAACTTTGTATAGAAAAGTTGGGGTTGCGCCTTTTCCAAGGCAGCCCTGGGTTTGCGCAGGGACGCGGCTGCTCTGGGCGTGGTTCCGGGAAACGCAGCGGCGCCGACCCTGGGTCTCGCACATTCTTCACGTCCGTTCGCAGCGTCACCCGGATCTTCGCCGCTACCCTTGTGGGCCCCCCGGCGACGCTTCCTGCTCCGCCCCTAAGTCGGGAAGGTTCCTTGCGGTTCGCGGCGTGCCGGACGTGACAAACGGAAGCCGCACGTCTCACTAGTACCCTCGCAGACGGACAGCGCCAGGGAGCAATGGCAGCGCGCCGACCGCGATGGGCTGTGGCCAATAGCGGCTGCTCAGCAGGGCGCGCCGAGAGCAGCGGCCGGGAAGGGGCGGTGCGGGAGGCGGGGTGTGGGGCGGTAGTGTGGGCCCTGTTCCTGCCCGCGCGGTGTTCCGCATTCTGCAAGCCTCCGGAGCGCACGTCGGCAGTCGGCTCCCTCGTTGACCGAATCACCGACCTCTCTCCCCAGGCAAGTTTGTACAAAAAAGCAGGCTGCCACCATGTCCGTCCTGACGCCGCTGCTGCTGCGGGGCTTGACAGGCTCGGCCCGGCGGCTCCCAGTGCCGCGCGCCAAGATCCATTCGTTGGGGGATCTGTCCGTCCTGACGCCGCTGCTGCTGCGGGGCTTGACAGGCTCGGCCCGGCGGCTCCCAGTGCCGCGCGCCAAGATCCATTCGTTGGGGGATCCACCGGTCGCCACCATGAGCGAGCTGATTAAGGAGAACATGCACATGAAGCTGTACATGGAGGGCACCGTGAACAACCACCACTTCAAGTGCACATCCGAGGGCGAAGGCAAGCCCTACGAGGGCACCCAGACCATGAGAATCAAGGTGGTCGAGGGCGGCCCTCTCCCCTTCGCCTTCGACATCCTGGCTACCAGCTTCATGTACGGCAGCAGAACCTTCATCAACCACACCCAGGGCATCCCCGACTTCTTTAAGCAGTCCTTCCCTGAGGGCTTCACATGGGAGAGAGTCACCACATACGAAGACGGGGGCGTGCTGACCGCTACCCAGGACACCAGCCTCCAGGACGGCTGCCTCATCTACAACGTCAAGATCAGAGGGGTGAACTTCCCATCCAACGGCCCTGTGATGCAGAAGAAAACACTCGGCTGGGAGGCCAACACCGAGATGCTGTACCCCGCTGACGGCGGCCTGGAAGGCAGAAGCGACATGGCCCTGAAGCTCGTGGGCGGGGGCCACCTGATCTGCAACTTCAAGACCACATACAGATCCAAGAAACCCGCTAAGAACCTCAAGATGCCCGGCGTCTACTATGTGGACCACAGACTGGAAAGAATCAAGGAGGCCGACAAAGAGACCTACGTCGAGCAGCACGAGGTGGCTGTGGCCAGATACTGCGACCTCCCTAGCAAACTGGGGCACAAGGCTGCCGTTAAATCTGGCACCAAAGCTAAACGTGCTCAGCGTCCGGCAAAATATAGCTACGTTGACGAAAACGGCGAAACTAAAACCTGGACTGGCCAAGGCCGTACTCCAGCTGTAATCAAAAAAGCAATGGATGAGCAAGGTAAATCCCTCGACGATTTCCTGATCAAGCAATACCCATACGATGTTCCAGATTACGCTTAAACCCAGCTTTCTTGTACAAAGTGGTGATAATCGAATTCCGATAATCAACCTCTGGATTACAAAATTTGTGAAAGATTGACTGGTATTCTTAACTATGTTGCTCCTTTTACGCTATGTGGATACGCTGCTTTAATGCCTTTGTATCATGCTATTGCTTCCCGTATGGCTTTCATTTTCTCCTCCTTGTATAAATCCTGGTTGCTGTCTCTTTATGAGGAGTTGTGGCCCGTTGTCAGGCAACGTGGCGTGGTGTGCACTGTGTTTGCTGACGCAACCCCCACTGGTTGGGGCATTGCCACCACCTGTCAGCTCCTTTCCGGGACTTTCGCTTTCCCCCTCCCTATTGCCACGGCGGAACTCATCGCCGCCTGCCTTGCCCGCTGCTGGACAGGGGCTCGGCTGTTGGGCACTGACAATTCCGTGGTGTTGTCGGGGAAGCTGACGTCCTTTCCATGGCTGCTCGCCTGTGTTGCCACCTGGATTCTGCGCGGGACGTCCTTCTGCTACGTCCCTTCGGCCCTCAATCCAGCGGACCTTCCTTCCCGCGGCCTGCTGCCGGCTCTGCGGCCTCTTCCGCGTCTTCGCCTTCGCCCTCAGACGAGTCGGATCTCCCTTTGGGCCGCCTCCCCGCATCGGGAATTCCCGCGGTTCGAATTCTACCGGGTAGGGGAGGCGCTTTTCCCAAGGCAGTCTGGAGCATGCGCTTTAGCAGCCCCGCTGGGCACTTGGCGCTACACAAGTGGCCTCTGGCCTCGCACACATTCCACATCCACCGGTAGGCGCCAACCGGCTCCGTTCTTTGGTGGCCCCTTCGCGCCACCTTCTACTCCTCCCCTAGTCAGGAAGTTCCCCCCCGCCCCGCAGCTCGCGTCGTGCAGGACGTGACAAATGGAAGTAGCACGTCTCACTAGTCTCGTGCAGATGGACAGCACCGCTGAGCAATGGAAGCGGGTAGGCCTTTGGGGCAGCGGCCAATAGCAGCTTTGCTCCTTCGCTTTCTGGGCTCAGAGGCTGGGAAGGGGTGGGTCCGGGGGCGGGCTCAGGGGCGGGCTCAGGGGCGGGGCGGGCGCCCGAAGGTCCTCCGGAGGCCCGGCATTCTGCACGCTTCAAAAGCGCACGTCTGCCGCGCTGTTCTCCTCTTCCTCATCTCCGGGCCTTTCGACCTCACGTGGCCACCATGACCGAGTACAAGCCCACGGTGCGCCTCGCCACCCGCGACGACGTCCCCAGGGCCGTACGCACCCTCGCCGCCGCGTTCGCCGACTACCCCGCCACGCGCCACACCGTCGATCCGGACCGCCACATCGAGCGGGTCACCGAGCTGCAAGAACTCTTCCTCACGCGCGTCGGGCTCGACATCGGCAAGGTGTGGGTCGCGGACGACGGCGCCGCGGTGGCGGTCTGGACCACGCCGGAGAGCGTCGAAGCGGGGGCGGTGTTCGCCGAGATCGGCCCGCGCATGGCCGAGTTGAGCGGTTCCCGGCTGGCCGCGCAGCAACAGATGGAAGGCCTCCTGGCGCCGCACCGGCCCAAGGAGCCCGCGTGGTTCCTGGCCACCGTCGGCGTCTCGCCCGACCACCAGGGCAAGGGTCTGGGCAGCGCCGTCGTGCTCCCCGGAGTGGAGGCGGCCGAGCGCGCCGGGGTGCCCGCCTTCCTGGAGACCTCCGCGCCCCGCAACCTCCCCTTCTACGAGCGGCTCGGCTTCACCGTCACCGCCGACGTCGAGGTGCCCGAAGGACCGCGCACCTGGTGCATGACCCGCAAGCCCGGTGCCTGAGGTACCTTTAAGACCAATGACTTACAAGGCAGCTGTAGATCTTAGCCACTTTTTAAAAGAAAAGGGGGGACTGGAAGGGCTAATTCACTCCCAACGAAGACAAGATCTGCTTTTTGCTTGTACTGGGTCTCTCTGGTTAGACCAGATCTGAGCCTGGGAGCTCTCTGGCTAACTAGGGAACCCACTGCTTAAGCCTCAATAAAGCTTGCCTTGAGTGCTTCAAGTAGTGTGTGCCCGTCTGTTGTGTGACTCTGGTAACTAGAGATCCCTCAGACCCTTTTAGTCAGTGTGGAAAATCTCTAGCAGTAGTAGTTCATGTCATCTTATTATTCAGTATTTATAACTTGCAAAGAAATGAATATCAGAGAGTGAGAGGAACTTGTTTATTGCAGCTTATAATGGTTACAAATAAAGCAATAGCATCACAAATTTCACAAATAAAGCATTTTTTTCACTGCATTCTAGTTGTGGTTTGTCCAAACTCATCAATGTATCTTATCATGTCTGGCTCTAGCTATCCCGCCCCTAACTCCGCCCATCCCGCCCCTAACTCCGCCCAGTTCCGCCCATTCTCCGCCCCATGGCTGACTAATTTTTTTTATTTATGCAGAGGCCGAGGCCGCCTCGGCCTCTGAGCTATTCCAGAAGTAGTGAGGAGGCTTTTTTGGAGGCCTAGGGACGTACCCAATTCGCCCTATAGTGAGTCGTATTACGCGCGCTCACTGGCCGTCGTTTTACAACGTCGTGACTGGGAAAACCCTGGCGTTACCCAACTTAATCGCCTTGCAGCACATCCCCCTTTCGCCAGCTGGCGTAATAGCGAAGAGGCCCGCACCGATCGCCCTTCCCAACAGTTGCGCAGCCTGAATGGCGAATGGGACGCGCCCTGTAGCGGCGCATTAAGCGCGGCGGGTGTGGTGGTTACGCGCAGCGTGACCGCTACACTTGCCAGCGCCCTAGCGCCCGCTCCTTTCGCTTTCTTCCCTTCCTTTCTCGCCACGTTCGCCGGCTTTCCCCGTCAAGCTCTAAATCGGGGGCTCCCTTTAGGGTTCCGATTTAGTGCTTTACGGCACCTCGACCCCAAAAAACTTGATTAGGGTGATGGTTCACGTAGTGGGCCATCGCCCTGATAGACGGTTTTTCGCCCTTTGACGTTGGAGTCCACGTTCTTTAATAGTGGACTCTTGTTCCAAACTGGAACAACACTCAACCCTATCTCGGTCTATTCTTTTGATTTATAAGGGATTTTGCCGATTTCGGCCTATTGGTTAAAAAATGAGCTGATTTAACAAAAATTTAACGCGAATTTTAACAAAATATTAACGCTTACAATTTAGGTGGCACTTTTCGGGGAAATGTGCGCGGAACCCCTATTTGTTTATTTTTCTAAATACATTCAAATATGTATCCGCTCATGAGACAATAACCCTGATAAATGCTTCAATAATATTGAAAAAGGAAGAGTATGAGTATTCAACATTTCCGTGTCGCCCTTATTCCCTTTTTTGCGGCATTTTGCCTTCCTGTTTTTGCTCACCCAGAAACGCTGGTGAAAGTAAAAGATGCTGAAGATCAGTTGGGTGCACGAGTGGGTTACATCGAACTGGATCTCAACAGCGGTAAGATCCTTGAGAGTTTTCGCCCCGAAGAACGTTTTCCAATGATGAGCACTTTTAAAGTTCTGCTATGTGGCGCGGTATTATCCCGTATTGACGCCGGGCAAGAGCAACTCGGTCGCCGCATACACTATTCTCAGAATGACTTGGTTGAGTACTCACCAGTCACAGAAAAGCATCTTACGGATGGCATGACAGTAAGAGAATTATGCAGTGCTGCCATAACCATGAGTGATAACACTGCGGCCAACTTACTTCTGACAACGATCGGAGGACCGAAGGAGCTAACCGCTTTTTTGCACAACATGGGGGATCATGTAACTCGCCTTGATCGTTGGGAACCGGAGCTGAATGAAGCCATACCAAACGACGAGCGTGACACCACGATGCCTGTAGCAATGGCAACAACGTTGCGCAAACTATTAACTGGCGAACTACTTACTCTAGCTTCCCGGCAACAATTAATAGACTGGATGGAGGCGGATAAAGTTGCAGGACCACTTCTGCGCTCGGCCCTTCCGGCTGGCTGGTTTATTGCTGATAAATCTGGAGCCGGTGAGCGTGGGTCTCGCGGTATCATTGCAGCACTGGGGCCAGATGGTAAGCCCTCCCGTATCGTAGTTATCTACACGACGGGGAGTCAGGCAACTATGGATGAACGAAATAGACAGATCGCTGAGATAGGTGCCTCACTGATTAAGCATTGGTAACTGTCAGACCAAGTTTACTCATATATACTTTAGATTGATTTAAAACTTCATTTTTAATTTAAAAGGATCTAGGTGAAGATCCTTTTTGATAATCTCATGACCAAAATCCCTTAACGTGAGTTTTCGTTCCACTGAGCGTCAGACCCCGTAGAAAAGATCAAAGGATCTTCTTGAGATCCTTTTTTTCTGCGCGTAATCTGCTGCTTGCAAACAAAAAAACCACCGCTACCAGCGGTGGTTTGTTTGCCGGATCAAGAGCTACCAACTCTTTTTCCGAAGGTAACTGGCTTCAGCAGAGCGCAGATACCAAATACTGTTCTTCTAGTGTAGCCGTAGTTAGGCCACCACTTCAAGAACTCTGTAGCACCGCCTACATACCTCGCTCTGCTAATCCTGTTACCAGTGGCTGCTGCCAGTGGCGATAAGTCGTGTCTTACCGGGTTGGACTCAAGACGATAGTTACCGGATAAGGCGCAGCGGTCGGGCTGAACGGGGGGTTCGTGCACACAGCCCAGCTTGGAGCGAACGACCTACACCGAACTGAGATACCTACAGCGTGAGCTATGAGAAAGCGCCACGCTTCCCGAAGAGAGAAAGGCGGACAGGTATCCGGTAAGCGGCAGGGTCGGAACAGGAGAGCGCACGAGGGAGCTTCCAGGGGGAAACGCCTGGTATCTTTATAGTCCTGTCGGGTTTCGCCACCTCTGACTTGAGCGTCGATTTTTGTGATGCTCGTCAGGGGGGCGGAGCCTATGGAAAAACGCCAGCAACGCGGCCTTTTTACGGTTCCTGGCCTTTTGCTGGCCTTTTGCTCACATGTTCTTTCCTGCGTTATCCCCTGATTCTGTGGATAACCGTATTACCGCCTTTGAGTGAGCTGATACCGCTCGCCGCAGCCGAACGACCGAGCGCAGCGAGTCAGTGAGCGAGGAAGCGGAAGAGCGCCCAATACGCAAACCGCCTCTCCCCGCGCGTTGGCCGATTCATTAATGCAGCTGGCACGACAGGTTTCCCGACTGGAAAGCGGGCAGTGAGCGCAACGCAATTAATGTGAGTTAGCTCACTCATTAGGCACCCCAGGCTTTACACTTTATGCTTCCGGCTCGTATGTTGTGTGGAATTGTGAGCGGATAACAATTTCACACAGGAAACAGCTATGACCATGATTACGCCAAGCGCGCAATTAACCCTCACTAAAGGGAACAAAAGCTGGAGCTGCAAGCTT

**> Mt-KillerRed-HI-NESS - VB230322-1503gum - pLV[Exp]-Puro-hPGK>{Mt-KillerRed-HI-NESS}**

AATGTAGTCTTATGCAATACTCTTGTAGTCTTGCAACATGGTAACGATGAGTTAGCAACATGCCTTACAAGGAGAGAAAAAGCACCGTGCATGCCGATTGGTGGAAGTAAGGTGGTACGATCGTGCCTTATTAGGAAGGCAACAGACGGGTCTGACATGGATTGGACGAACCACTGAATTGCCGCATTGCAGAGATATTGTATTTAAGTGCCTAGCTCGATACATAAACGGGTCTCTCTGGTTAGACCAGATCTGAGCCTGGGAGCTCTCTGGCTAACTAGGGAACCCACTGCTTAAGCCTCAATAAAGCTTGCCTTGAGTGCTTCAAGTAGTGTGTGCCCGTCTGTTGTGTGACTCTGGTAACTAGAGATCCCTCAGACCCTTTTAGTCAGTGTGGAAAATCTCTAGCAGTGGCGCCCGAACAGGGACTTGAAAGCGAAAGGGAAACCAGAGGAGCTCTCTCGACGCAGGACTCGGCTTGCTGAAGCGCGCACGGCAAGAGGCGAGGGGCGGCGACTGGTGAGTACGCCAAAAATTTTGACTAGCGGAGGCTAGAAGGAGAGAGATGGGTGCGAGAGCGTCAGTATTAAGCGGGGGAGAATTAGATCGCGATGGGAAAAAATTCGGTTAAGGCCAGGGGGAAAGAAAAAATATAAATTAAAACATATAGTATGGGCAAGCAGGGAGCTAGAACGATTCGCAGTTAATCCTGGCCTGTTAGAAACATCAGAAGGCTGTAGACAAATACTGGGACAGCTACAACCATCCCTTCAGACAGGATCAGAAGAACTTAGATCATTATATAATACAGTAGCAACCCTCTATTGTGTGCATCAAAGGATAGAGATAAAAGACACCAAGGAAGCTTTAGACAAGATAGAGGAAGAGCAAAACAAAAGTAAGACCACCGCACAGCAAGCGGCCGCTGATCTTCAGACCTGGAGGAGGAGATATGAGGGACAATTGGAGAAGTGAATTATATAAATATAAAGTAGTAAAAATTGAACCATTAGGAGTAGCACCCACCAAGGCAAAGAGAAGAGTGGTGCAGAGAGAAAAAAGAGCAGTGGGAATAGGAGCTTTGTTCCTTGGGTTCTTGGGAGCAGCAGGAAGCACTATGGGCGCAGCGTCAATGACGCTGACGGTACAGGCCAGACAATTATTGTCTGGTATAGTGCAGCAGCAGAACAATTTGCTGAGGGCTATTGAGGCGCAACAGCATCTGTTGCAACTCACAGTCTGGGGCATCAAGCAGCTCCAGGCAAGAATCCTGGCTGTGGAAAGATACCTAAAGGATCAACAGCTCCTGGGGATTTGGGGTTGCTCTGGAAAACTCATTTGCACCACTGCTGTGCCTTGGAATGCTAGTTGGAGTAATAAATCTCTGGAACAGATTTGGAATCACACGACCTGGATGGAGTGGGACAGAGAAATTAACAATTACACAAGCTTAATACACTCCTTAATTGAAGAATCGCAAAACCAGCAAGAAAAGAATGAACAAGAATTATTGGAATTAGATAAATGGGCAAGTTTGTGGAATTGGTTTAACATAACAAATTGGCTGTGGTATATAAAATTATTCATAATGATAGTAGGAGGCTTGGTAGGTTTAAGAATAGTTTTTGCTGTACTTTCTATAGTGAATAGAGTTAGGCAGGGATATTCACCATTATCGTTTCAGACCCACCTCCCAACCCCGAGGGGACCCGACAGGCCCGAAGGAATAGAAGAAGAAGGTGGAGAGAGAGACAGAGACAGATCCATTCGATTAGTGAACGGATCTCGACGGTATCGCTAGCTTTTAAAAGAAAAGGGGGGATTGGGGGGTACAGTGCAGGGGAAAGAATAGTAGACATAATAGCAACAGACATACAAACTAAAGAATTACAAAAACAAATTACAAAAATTCAAAATTTTACTAGTGATTATCGGATCAACTTTGTATAGAAAAGTTGGGGTTGCGCCTTTTCCAAGGCAGCCCTGGGTTTGCGCAGGGACGCGGCTGCTCTGGGCGTGGTTCCGGGAAACGCAGCGGCGCCGACCCTGGGTCTCGCACATTCTTCACGTCCGTTCGCAGCGTCACCCGGATCTTCGCCGCTACCCTTGTGGGCCCCCCGGCGACGCTTCCTGCTCCGCCCCTAAGTCGGGAAGGTTCCTTGCGGTTCGCGGCGTGCCGGACGTGACAAACGGAAGCCGCACGTCTCACTAGTACCCTCGCAGACGGACAGCGCCAGGGAGCAATGGCAGCGCGCCGACCGCGATGGGCTGTGGCCAATAGCGGCTGCTCAGCAGGGCGCGCCGAGAGCAGCGGCCGGGAAGGGGCGGTGCGGGAGGCGGGGTGTGGGGCGGTAGTGTGGGCCCTGTTCCTGCCCGCGCGGTGTTCCGCATTCTGCAAGCCTCCGGAGCGCACGTCGGCAGTCGGCTCCCTCGTTGACCGAATCACCGACCTCTCTCCCCAGGCAAGTTTGTACAAAAAAGCAGGCTGCCACCATGTCCGTCCTGACGCCGCTGCTGCTGCGGGGCTTGACAGGCTCGGCCCGGCGGCTCCCAGTGCCGCGCGCCAAGATCCATTCGTTGGGGGATCTGTCCGTCCTGACGCCGCTGCTGCTGCGGGGCTTGACAGGCTCGGCCCGGCGGCTCCCAGTGCCGCGCGCCAAGATCCATTCGTTGGGGGATCCACCGGTCGCCACCATGGGTTCAGAGGGCGGCCCCGCCCTGTTCCAGAGCGACATGACCTTCAAAATCTTCATCGACGGCGAGGTGAACGGCCAGAAGTTCACCATCGTGGCCGACGGCAGCAGCAAGTTCCCCCACGGCGACTTCAACGTGCACGCCGTGTGCGAGACCGGCAAGCTGCCCATGAGCTGGAAGCCCATCTGCCACCTGATCCAGTACGGCGAGCCCTTCTTCGCCCGCTACCCCGACGGCATCAGCCATTTCGCCCAGGAGTGCTTCCCCGAGGGCCTGAGCATCGACCGCACCGTGCGCTTCGAGAACGACGGCACCATGACCAGCCACCACACCTACGAGCTGGACGACACCTGCGTGGTGAGCCGCATCACCGTGAACTGCGACGGCTTCCAGCCCGACGGCCCCATCATGCGCGACCAGCTGGTGGACATCCTGCCCAACGAGACCCACATGTTCCCCCACGGCCCCAACGCCGTGCGCCAGCTGGCCTTCATCGGCTTCACCACCGCCGACGGCGGCCTGATGATGGGCCACTTCGACAGCAAGATGACCTTCAACGGCAGCCGCGCCATCGAGATCCCCGGCCCACACTTCGTGACCATCATCACCAAGCAGATGAGGGACACCAGCGACAAGCGCGACCACGTGTGCCAGCGCGAGGTGGCCTACGCCCACAGCGTGCCCCGCATCACCAGCGCCATCGGTAGCGACGAGGATGCTGCCGTTAAATCTGGCACCAAAGCTAAACGTGCTCAGCGTCCGGCAAAATATAGCTACGTTGACGAAAACGGCGAAACTAAAACCTGGACTGGCCAAGGCCGTACTCCAGCTGTAATCAAAAAAGCAATGGATGAGCAAGGTAAATCCCTCGACGATTTCCTGATCAAGCAATACCCATACGATGTTCCAGATTACGCTTAAACCCAGCTTTCTTGTACAAAGTGGTGATAATCGAATTCCGATAATCAACCTCTGGATTACAAAATTTGTGAAAGATTGACTGGTATTCTTAACTATGTTGCTCCTTTTACGCTATGTGGATACGCTGCTTTAATGCCTTTGTATCATGCTATTGCTTCCCGTATGGCTTTCATTTTCTCCTCCTTGTATAAATCCTGGTTGCTGTCTCTTTATGAGGAGTTGTGGCCCGTTGTCAGGCAACGTGGCGTGGTGTGCACTGTGTTTGCTGACGCAACCCCCACTGGTTGGGGCATTGCCACCACCTGTCAGCTCCTTTCCGGGACTTTCGCTTTCCCCCTCCCTATTGCCACGGCGGAACTCATCGCCGCCTGCCTTGCCCGCTGCTGGACAGGGGCTCGGCTGTTGGGCACTGACAATTCCGTGGTGTTGTCGGGGAAGCTGACGTCCTTTCCATGGCTGCTCGCCTGTGTTGCCACCTGGATTCTGCGCGGGACGTCCTTCTGCTACGTCCCTTCGGCCCTCAATCCAGCGGACCTTCCTTCCCGCGGCCTGCTGCCGGCTCTGCGGCCTCTTCCGCGTCTTCGCCTTCGCCCTCAGACGAGTCGGATCTCCCTTTGGGCCGCCTCCCCGCATCGGGAATTCCCGCGGTTCGAATTCTACCGGGTAGGGGAGGCGCTTTTCCCAAGGCAGTCTGGAGCATGCGCTTTAGCAGCCCCGCTGGGCACTTGGCGCTACACAAGTGGCCTCTGGCCTCGCACACATTCCACATCCACCGGTAGGCGCCAACCGGCTCCGTTCTTTGGTGGCCCCTTCGCGCCACCTTCTACTCCTCCCCTAGTCAGGAAGTTCCCCCCCGCCCCGCAGCTCGCGTCGTGCAGGACGTGACAAATGGAAGTAGCACGTCTCACTAGTCTCGTGCAGATGGACAGCACCGCTGAGCAATGGAAGCGGGTAGGCCTTTGGGGCAGCGGCCAATAGCAGCTTTGCTCCTTCGCTTTCTGGGCTCAGAGGCTGGGAAGGGGTGGGTCCGGGGGCGGGCTCAGGGGCGGGCTCAGGGGCGGGGCGGGCGCCCGAAGGTCCTCCGGAGGCCCGGCATTCTGCACGCTTCAAAAGCGCACGTCTGCCGCGCTGTTCTCCTCTTCCTCATCTCCGGGCCTTTCGACCTCACGTGGCCACCATGACCGAGTACAAGCCCACGGTGCGCCTCGCCACCCGCGACGACGTCCCCAGGGCCGTACGCACCCTCGCCGCCGCGTTCGCCGACTACCCCGCCACGCGCCACACCGTCGATCCGGACCGCCACATCGAGCGGGTCACCGAGCTGCAAGAACTCTTCCTCACGCGCGTCGGGCTCGACATCGGCAAGGTGTGGGTCGCGGACGACGGCGCCGCGGTGGCGGTCTGGACCACGCCGGAGAGCGTCGAAGCGGGGGCGGTGTTCGCCGAGATCGGCCCGCGCATGGCCGAGTTGAGCGGTTCCCGGCTGGCCGCGCAGCAACAGATGGAAGGCCTCCTGGCGCCGCACCGGCCCAAGGAGCCCGCGTGGTTCCTGGCCACCGTCGGCGTCTCGCCCGACCACCAGGGCAAGGGTCTGGGCAGCGCCGTCGTGCTCCCCGGAGTGGAGGCGGCCGAGCGCGCCGGGGTGCCCGCCTTCCTGGAGACCTCCGCGCCCCGCAACCTCCCCTTCTACGAGCGGCTCGGCTTCACCGTCACCGCCGACGTCGAGGTGCCCGAAGGACCGCGCACCTGGTGCATGACCCGCAAGCCCGGTGCCTGAGGTACCTTTAAGACCAATGACTTACAAGGCAGCTGTAGATCTTAGCCACTTTTTAAAAGAAAAGGGGGGACTGGAAGGGCTAATTCACTCCCAACGAAGACAAGATCTGCTTTTTGCTTGTACTGGGTCTCTCTGGTTAGACCAGATCTGAGCCTGGGAGCTCTCTGGCTAACTAGGGAACCCACTGCTTAAGCCTCAATAAAGCTTGCCTTGAGTGCTTCAAGTAGTGTGTGCCCGTCTGTTGTGTGACTCTGGTAACTAGAGATCCCTCAGACCCTTTTAGTCAGTGTGGAAAATCTCTAGCAGTAGTAGTTCATGTCATCTTATTATTCAGTATTTATAACTTGCAAAGAAATGAATATCAGAGAGTGAGAGGAACTTGTTTATTGCAGCTTATAATGGTTACAAATAAAGCAATAGCATCACAAATTTCACAAATAAAGCATTTTTTTCACTGCATTCTAGTTGTGGTTTGTCCAAACTCATCAATGTATCTTATCATGTCTGGCTCTAGCTATCCCGCCCCTAACTCCGCCCATCCCGCCCCTAACTCCGCCCAGTTCCGCCCATTCTCCGCCCCATGGCTGACTAATTTTTTTTATTTATGCAGAGGCCGAGGCCGCCTCGGCCTCTGAGCTATTCCAGAAGTAGTGAGGAGGCTTTTTTGGAGGCCTAGGGACGTACCCAATTCGCCCTATAGTGAGTCGTATTACGCGCGCTCACTGGCCGTCGTTTTACAACGTCGTGACTGGGAAAACCCTGGCGTTACCCAACTTAATCGCCTTGCAGCACATCCCCCTTTCGCCAGCTGGCGTAATAGCGAAGAGGCCCGCACCGATCGCCCTTCCCAACAGTTGCGCAGCCTGAATGGCGAATGGGACGCGCCCTGTAGCGGCGCATTAAGCGCGGCGGGTGTGGTGGTTACGCGCAGCGTGACCGCTACACTTGCCAGCGCCCTAGCGCCCGCTCCTTTCGCTTTCTTCCCTTCCTTTCTCGCCACGTTCGCCGGCTTTCCCCGTCAAGCTCTAAATCGGGGGCTCCCTTTAGGGTTCCGATTTAGTGCTTTACGGCACCTCGACCCCAAAAAACTTGATTAGGGTGATGGTTCACGTAGTGGGCCATCGCCCTGATAGACGGTTTTTCGCCCTTTGACGTTGGAGTCCACGTTCTTTAATAGTGGACTCTTGTTCCAAACTGGAACAACACTCAACCCTATCTCGGTCTATTCTTTTGATTTATAAGGGATTTTGCCGATTTCGGCCTATTGGTTAAAAAATGAGCTGATTTAACAAAAATTTAACGCGAATTTTAACAAAATATTAACGCTTACAATTTAGGTGGCACTTTTCGGGGAAATGTGCGCGGAACCCCTATTTGTTTATTTTTCTAAATACATTCAAATATGTATCCGCTCATGAGACAATAACCCTGATAAATGCTTCAATAATATTGAAAAAGGAAGAGTATGAGTATTCAACATTTCCGTGTCGCCCTTATTCCCTTTTTTGCGGCATTTTGCCTTCCTGTTTTTGCTCACCCAGAAACGCTGGTGAAAGTAAAAGATGCTGAAGATCAGTTGGGTGCACGAGTGGGTTACATCGAACTGGATCTCAACAGCGGTAAGATCCTTGAGAGTTTTCGCCCCGAAGAACGTTTTCCAATGATGAGCACTTTTAAAGTTCTGCTATGTGGCGCGGTATTATCCCGTATTGACGCCGGGCAAGAGCAACTCGGTCGCCGCATACACTATTCTCAGAATGACTTGGTTGAGTACTCACCAGTCACAGAAAAGCATCTTACGGATGGCATGACAGTAAGAGAATTATGCAGTGCTGCCATAACCATGAGTGATAACACTGCGGCCAACTTACTTCTGACAACGATCGGAGGACCGAAGGAGCTAACCGCTTTTTTGCACAACATGGGGGATCATGTAACTCGCCTTGATCGTTGGGAACCGGAGCTGAATGAAGCCATACCAAACGACGAGCGTGACACCACGATGCCTGTAGCAATGGCAACAACGTTGCGCAAACTATTAACTGGCGAACTACTTACTCTAGCTTCCCGGCAACAATTAATAGACTGGATGGAGGCGGATAAAGTTGCAGGACCACTTCTGCGCTCGGCCCTTCCGGCTGGCTGGTTTATTGCTGATAAATCTGGAGCCGGTGAGCGTGGGTCTCGCGGTATCATTGCAGCACTGGGGCCAGATGGTAAGCCCTCCCGTATCGTAGTTATCTACACGACGGGGAGTCAGGCAACTATGGATGAACGAAATAGACAGATCGCTGAGATAGGTGCCTCACTGATTAAGCATTGGTAACTGTCAGACCAAGTTTACTCATATATACTTTAGATTGATTTAAAACTTCATTTTTAATTTAAAAGGATCTAGGTGAAGATCCTTTTTGATAATCTCATGACCAAAATCCCTTAACGTGAGTTTTCGTTCCACTGAGCGTCAGACCCCGTAGAAAAGATCAAAGGATCTTCTTGAGATCCTTTTTTTCTGCGCGTAATCTGCTGCTTGCAAACAAAAAAACCACCGCTACCAGCGGTGGTTTGTTTGCCGGATCAAGAGCTACCAACTCTTTTTCCGAAGGTAACTGGCTTCAGCAGAGCGCAGATACCAAATACTGTTCTTCTAGTGTAGCCGTAGTTAGGCCACCACTTCAAGAACTCTGTAGCACCGCCTACATACCTCGCTCTGCTAATCCTGTTACCAGTGGCTGCTGCCAGTGGCGATAAGTCGTGTCTTACCGGGTTGGACTCAAGACGATAGTTACCGGATAAGGCGCAGCGGTCGGGCTGAACGGGGGGTTCGTGCACACAGCCCAGCTTGGAGCGAACGACCTACACCGAACTGAGATACCTACAGCGTGAGCTATGAGAAAGCGCCACGCTTCCCGAAGAGAGAAAGGCGGACAGGTATCCGGTAAGCGGCAGGGTCGGAACAGGAGAGCGCACGAGGGAGCTTCCAGGGGGAAACGCCTGGTATCTTTATAGTCCTGTCGGGTTTCGCCACCTCTGACTTGAGCGTCGATTTTTGTGATGCTCGTCAGGGGGGCGGAGCCTATGGAAAAACGCCAGCAACGCGGCCTTTTTACGGTTCCTGGCCTTTTGCTGGCCTTTTGCTCACATGTTCTTTCCTGCGTTATCCCCTGATTCTGTGGATAACCGTATTACCGCCTTTGAGTGAGCTGATACCGCTCGCCGCAGCCGAACGACCGAGCGCAGCGAGTCAGTGAGCGAGGAAGCGGAAGAGCGCCCAATACGCAAACCGCCTCTCCCCGCGCGTTGGCCGATTCATTAATGCAGCTGGCACGACAGGTTTCCCGACTGGAAAGCGGGCAGTGAGCGCAACGCAATTAATGTGAGTTAGCTCACTCATTAGGCACCCCAGGCTTTACACTTTATGCTTCCGGCTCGTATGTTGTGTGGAATTGTGAGCGGATAACAATTTCACACAGGAAACAGCTATGACCATGATTACGCCAAGCGCGCAATTAACCCTCACTAAAGGGAACAAAAGCTGGAGCTGCAAGCTT

**> Mt-mRFP1-HI-NESS - VB241124-1212gaj - pLV[Exp]-Puro-hPGK>{Mt-Mrfp1-HI-NESS}**

AATGTAGTCTTATGCAATACTCTTGTAGTCTTGCAACATGGTAACGATGAGTTAGCAACATGCCTTACAAGGAGAGAAAAAGCACCGTGCATGCCGATTGGTGGAAGTAAGGTGGTACGATCGTGCCTTATTAGGAAGGCAACAGACGGGTCTGACATGGATTGGACGAACCACTGAATTGCCGCATTGCAGAGATATTGTATTTAAGTGCCTAGCTCGATACATAAACGGGTCTCTCTGGTTAGACCAGATCTGAGCCTGGGAGCTCTCTGGCTAACTAGGGAACCCACTGCTTAAGCCTCAATAAAGCTTGCCTTGAGTGCTTCAAGTAGTGTGTGCCCGTCTGTTGTGTGACTCTGGTAACTAGAGATCCCTCAGACCCTTTTAGTCAGTGTGGAAAATCTCTAGCAGTGGCGCCCGAACAGGGACTTGAAAGCGAAAGGGAAACCAGAGGAGCTCTCTCGACGCAGGACTCGGCTTGCTGAAGCGCGCACGGCAAGAGGCGAGGGGCGGCGACTGGTGAGTACGCCAAAAATTTTGACTAGCGGAGGCTAGAAGGAGAGAGATGGGTGCGAGAGCGTCAGTATTAAGCGGGGGAGAATTAGATCGCGATGGGAAAAAATTCGGTTAAGGCCAGGGGGAAAGAAAAAATATAAATTAAAACATATAGTATGGGCAAGCAGGGAGCTAGAACGATTCGCAGTTAATCCTGGCCTGTTAGAAACATCAGAAGGCTGTAGACAAATACTGGGACAGCTACAACCATCCCTTCAGACAGGATCAGAAGAACTTAGATCATTATATAATACAGTAGCAACCCTCTATTGTGTGCATCAAAGGATAGAGATAAAAGACACCAAGGAAGCTTTAGACAAGATAGAGGAAGAGCAAAACAAAAGTAAGACCACCGCACAGCAAGCGGCCGCTGATCTTCAGACCTGGAGGAGGAGATATGAGGGACAATTGGAGAAGTGAATTATATAAATATAAAGTAGTAAAAATTGAACCATTAGGAGTAGCACCCACCAAGGCAAAGAGAAGAGTGGTGCAGAGAGAAAAAAGAGCAGTGGGAATAGGAGCTTTGTTCCTTGGGTTCTTGGGAGCAGCAGGAAGCACTATGGGCGCAGCGTCAATGACGCTGACGGTACAGGCCAGACAATTATTGTCTGGTATAGTGCAGCAGCAGAACAATTTGCTGAGGGCTATTGAGGCGCAACAGCATCTGTTGCAACTCACAGTCTGGGGCATCAAGCAGCTCCAGGCAAGAATCCTGGCTGTGGAAAGATACCTAAAGGATCAACAGCTCCTGGGGATTTGGGGTTGCTCTGGAAAACTCATTTGCACCACTGCTGTGCCTTGGAATGCTAGTTGGAGTAATAAATCTCTGGAACAGATTTGGAATCACACGACCTGGATGGAGTGGGACAGAGAAATTAACAATTACACAAGCTTAATACACTCCTTAATTGAAGAATCGCAAAACCAGCAAGAAAAGAATGAACAAGAATTATTGGAATTAGATAAATGGGCAAGTTTGTGGAATTGGTTTAACATAACAAATTGGCTGTGGTATATAAAATTATTCATAATGATAGTAGGAGGCTTGGTAGGTTTAAGAATAGTTTTTGCTGTACTTTCTATAGTGAATAGAGTTAGGCAGGGATATTCACCATTATCGTTTCAGACCCACCTCCCAACCCCGAGGGGACCCGACAGGCCCGAAGGAATAGAAGAAGAAGGTGGAGAGAGAGACAGAGACAGATCCATTCGATTAGTGAACGGATCTCGACGGTATCGCTAGCTTTTAAAAGAAAAGGGGGGATTGGGGGGTACAGTGCAGGGGAAAGAATAGTAGACATAATAGCAACAGACATACAAACTAAAGAATTACAAAAACAAATTACAAAAATTCAAAATTTTACTAGTGATTATCGGATCAACTTTGTATAGAAAAGTTGGGGTTGCGCCTTTTCCAAGGCAGCCCTGGGTTTGCGCAGGGACGCGGCTGCTCTGGGCGTGGTTCCGGGAAACGCAGCGGCGCCGACCCTGGGTCTCGCACATTCTTCACGTCCGTTCGCAGCGTCACCCGGATCTTCGCCGCTACCCTTGTGGGCCCCCCGGCGACGCTTCCTGCTCCGCCCCTAAGTCGGGAAGGTTCCTTGCGGTTCGCGGCGTGCCGGACGTGACAAACGGAAGCCGCACGTCTCACTAGTACCCTCGCAGACGGACAGCGCCAGGGAGCAATGGCAGCGCGCCGACCGCGATGGGCTGTGGCCAATAGCGGCTGCTCAGCAGGGCGCGCCGAGAGCAGCGGCCGGGAAGGGGCGGTGCGGGAGGCGGGGTGTGGGGCGGTAGTGTGGGCCCTGTTCCTGCCCGCGCGGTGTTCCGCATTCTGCAAGCCTCCGGAGCGCACGTCGGCAGTCGGCTCCCTCGTTGACCGAATCACCGACCTCTCTCCCCAGGCAAGTTTGTACAAAAAAGCAGGCTGCCACCATGTCCGTCCTGACGCCGCTGCTGCTGCGGGGCTTGACAGGCTCGGCCCGGCGGCTCCCAGTGCCGCGCGCCAAGATCCATTCGTTGGGGGATCTGTCCGTCCTGACGCCGCTGCTGCTGCGGGGCTTGACAGGCTCGGCCCGGCGGCTCCCAGTGCCGCGCGCCAAGATCCATTCGTTGGGGGATCCACCGGTCGCCACCATGGCCTCCTCCGAGGACGTCATCAAGGAGTTCATGCGCTTCAAGGTGCGCATGGAGGGCTCCGTGAACGGCCACGAGTTCGAGATCGAGGGCGAGGGCGAGGGCCGCCCCTACGAGGGCACCCAGACCGCCAAGCTGAAGGTGACCAAGGGCGGCCCCCTGCCCTTCGCCTGGGACATCCTGTCCCCTCAGTTCCAGTACGGCTCCAAGGCCTACGTGAAGCACCCCGCCGACATCCCCGACTACTTGAAGCTGTCCTTCCCCGAGGGCTTCAAGTGGGAGCGCGTGATGAACTTCGAGGACGGCGGCGTGGTGACCGTGACCCAGGACTCCTCCCTGCAGGACGGCGAGTTCATCTACAAGGTGAAGCTGCGCGGCACCAACTTCCCCTCCGACGGCCCCGTAATGCAGAAGAAGACCATGGGCTGGGAGGCCTCCACCGAGCGGATGTACCCCGAGGACGGCGCCCTGAAGGGCGAGATCAAGATGAGGCTGAAGCTGAAGGACGGCGGCCACTACGACGCCGAGGTCAAGACCACCTACATGGCCAAGAAGCCCGTGCAGCTGCCCGGCGCCTACAAGACCGACATCAAGCTGGACATCACCTCCCACAACGAGGACTACACCATCGTGGAACAGTACGAGCGCGCCGAGGGCCGCCACTCCACCGGCGCCGCTGCCGTTAAATCTGGCACCAAAGCTAAACGTGCTCAGCGTCCGGCAAAATATAGCTACGTTGACGAAAACGGCGAAACTAAAACCTGGACTGGCCAAGGCCGTACTCCAGCTGTAATCAAAAAAGCAATGGATGAGCAAGGTAAATCCCTCGACGATTTCCTGATCAAGCAATACCCATACGATGTTCCAGATTACGCTTAAACCCAGCTTTCTTGTACAAAGTGGTGATAATCGAATTCCGATAATCAACCTCTGGATTACAAAATTTGTGAAAGATTGACTGGTATTCTTAACTATGTTGCTCCTTTTACGCTATGTGGATACGCTGCTTTAATGCCTTTGTATCATGCTATTGCTTCCCGTATGGCTTTCATTTTCTCCTCCTTGTATAAATCCTGGTTGCTGTCTCTTTATGAGGAGTTGTGGCCCGTTGTCAGGCAACGTGGCGTGGTGTGCACTGTGTTTGCTGACGCAACCCCCACTGGTTGGGGCATTGCCACCACCTGTCAGCTCCTTTCCGGGACTTTCGCTTTCCCCCTCCCTATTGCCACGGCGGAACTCATCGCCGCCTGCCTTGCCCGCTGCTGGACAGGGGCTCGGCTGTTGGGCACTGACAATTCCGTGGTGTTGTCGGGGAAGCTGACGTCCTTTCCATGGCTGCTCGCCTGTGTTGCCACCTGGATTCTGCGCGGGACGTCCTTCTGCTACGTCCCTTCGGCCCTCAATCCAGCGGACCTTCCTTCCCGCGGCCTGCTGCCGGCTCTGCGGCCTCTTCCGCGTCTTCGCCTTCGCCCTCAGACGAGTCGGATCTCCCTTTGGGCCGCCTCCCCGCATCGGGAATTCCCGCGGTTCGAATTCTACCGGGTAGGGGAGGCGCTTTTCCCAAGGCAGTCTGGAGCATGCGCTTTAGCAGCCCCGCTGGGCACTTGGCGCTACACAAGTGGCCTCTGGCCTCGCACACATTCCACATCCACCGGTAGGCGCCAACCGGCTCCGTTCTTTGGTGGCCCCTTCGCGCCACCTTCTACTCCTCCCCTAGTCAGGAAGTTCCCCCCCGCCCCGCAGCTCGCGTCGTGCAGGACGTGACAAATGGAAGTAGCACGTCTCACTAGTCTCGTGCAGATGGACAGCACCGCTGAGCAATGGAAGCGGGTAGGCCTTTGGGGCAGCGGCCAATAGCAGCTTTGCTCCTTCGCTTTCTGGGCTCAGAGGCTGGGAAGGGGTGGGTCCGGGGGCGGGCTCAGGGGCGGGCTCAGGGGCGGGGCGGGCGCCCGAAGGTCCTCCGGAGGCCCGGCATTCTGCACGCTTCAAAAGCGCACGTCTGCCGCGCTGTTCTCCTCTTCCTCATCTCCGGGCCTTTCGACCTCACGTGGCCACCATGACCGAGTACAAGCCCACGGTGCGCCTCGCCACCCGCGACGACGTCCCCAGGGCCGTACGCACCCTCGCCGCCGCGTTCGCCGACTACCCCGCCACGCGCCACACCGTCGATCCGGACCGCCACATCGAGCGGGTCACCGAGCTGCAAGAACTCTTCCTCACGCGCGTCGGGCTCGACATCGGCAAGGTGTGGGTCGCGGACGACGGCGCCGCGGTGGCGGTCTGGACCACGCCGGAGAGCGTCGAAGCGGGGGCGGTGTTCGCCGAGATCGGCCCGCGCATGGCCGAGTTGAGCGGTTCCCGGCTGGCCGCGCAGCAACAGATGGAAGGCCTCCTGGCGCCGCACCGGCCCAAGGAGCCCGCGTGGTTCCTGGCCACCGTCGGCGTCTCGCCCGACCACCAGGGCAAGGGTCTGGGCAGCGCCGTCGTGCTCCCCGGAGTGGAGGCGGCCGAGCGCGCCGGGGTGCCCGCCTTCCTGGAGACCTCCGCGCCCCGCAACCTCCCCTTCTACGAGCGGCTCGGCTTCACCGTCACCGCCGACGTCGAGGTGCCCGAAGGACCGCGCACCTGGTGCATGACCCGCAAGCCCGGTGCCTGAGGTACCTTTAAGACCAATGACTTACAAGGCAGCTGTAGATCTTAGCCACTTTTTAAAAGAAAAGGGGGGACTGGAAGGGCTAATTCACTCCCAACGAAGACAAGATCTGCTTTTTGCTTGTACTGGGTCTCTCTGGTTAGACCAGATCTGAGCCTGGGAGCTCTCTGGCTAACTAGGGAACCCACTGCTTAAGCCTCAATAAAGCTTGCCTTGAGTGCTTCAAGTAGTGTGTGCCCGTCTGTTGTGTGACTCTGGTAACTAGAGATCCCTCAGACCCTTTTAGTCAGTGTGGAAAATCTCTAGCAGTAGTAGTTCATGTCATCTTATTATTCAGTATTTATAACTTGCAAAGAAATGAATATCAGAGAGTGAGAGGAACTTGTTTATTGCAGCTTATAATGGTTACAAATAAAGCAATAGCATCACAAATTTCACAAATAAAGCATTTTTTTCACTGCATTCTAGTTGTGGTTTGTCCAAACTCATCAATGTATCTTATCATGTCTGGCTCTAGCTATCCCGCCCCTAACTCCGCCCATCCCGCCCCTAACTCCGCCCAGTTCCGCCCATTCTCCGCCCCATGGCTGACTAATTTTTTTTATTTATGCAGAGGCCGAGGCCGCCTCGGCCTCTGAGCTATTCCAGAAGTAGTGAGGAGGCTTTTTTGGAGGCCTAGGGACGTACCCAATTCGCCCTATAGTGAGTCGTATTACGCGCGCTCACTGGCCGTCGTTTTACAACGTCGTGACTGGGAAAACCCTGGCGTTACCCAACTTAATCGCCTTGCAGCACATCCCCCTTTCGCCAGCTGGCGTAATAGCGAAGAGGCCCGCACCGATCGCCCTTCCCAACAGTTGCGCAGCCTGAATGGCGAATGGGACGCGCCCTGTAGCGGCGCATTAAGCGCGGCGGGTGTGGTGGTTACGCGCAGCGTGACCGCTACACTTGCCAGCGCCCTAGCGCCCGCTCCTTTCGCTTTCTTCCCTTCCTTTCTCGCCACGTTCGCCGGCTTTCCCCGTCAAGCTCTAAATCGGGGGCTCCCTTTAGGGTTCCGATTTAGTGCTTTACGGCACCTCGACCCCAAAAAACTTGATTAGGGTGATGGTTCACGTAGTGGGCCATCGCCCTGATAGACGGTTTTTCGCCCTTTGACGTTGGAGTCCACGTTCTTTAATAGTGGACTCTTGTTCCAAACTGGAACAACACTCAACCCTATCTCGGTCTATTCTTTTGATTTATAAGGGATTTTGCCGATTTCGGCCTATTGGTTAAAAAATGAGCTGATTTAACAAAAATTTAACGCGAATTTTAACAAAATATTAACGCTTACAATTTAGGTGGCACTTTTCGGGGAAATGTGCGCGGAACCCCTATTTGTTTATTTTTCTAAATACATTCAAATATGTATCCGCTCATGAGACAATAACCCTGATAAATGCTTCAATAATATTGAAAAAGGAAGAGTATGAGTATTCAACATTTCCGTGTCGCCCTTATTCCCTTTTTTGCGGCATTTTGCCTTCCTGTTTTTGCTCACCCAGAAACGCTGGTGAAAGTAAAAGATGCTGAAGATCAGTTGGGTGCACGAGTGGGTTACATCGAACTGGATCTCAACAGCGGTAAGATCCTTGAGAGTTTTCGCCCCGAAGAACGTTTTCCAATGATGAGCACTTTTAAAGTTCTGCTATGTGGCGCGGTATTATCCCGTATTGACGCCGGGCAAGAGCAACTCGGTCGCCGCATACACTATTCTCAGAATGACTTGGTTGAGTACTCACCAGTCACAGAAAAGCATCTTACGGATGGCATGACAGTAAGAGAATTATGCAGTGCTGCCATAACCATGAGTGATAACACTGCGGCCAACTTACTTCTGACAACGATCGGAGGACCGAAGGAGCTAACCGCTTTTTTGCACAACATGGGGGATCATGTAACTCGCCTTGATCGTTGGGAACCGGAGCTGAATGAAGCCATACCAAACGACGAGCGTGACACCACGATGCCTGTAGCAATGGCAACAACGTTGCGCAAACTATTAACTGGCGAACTACTTACTCTAGCTTCCCGGCAACAATTAATAGACTGGATGGAGGCGGATAAAGTTGCAGGACCACTTCTGCGCTCGGCCCTTCCGGCTGGCTGGTTTATTGCTGATAAATCTGGAGCCGGTGAGCGTGGGTCTCGCGGTATCATTGCAGCACTGGGGCCAGATGGTAAGCCCTCCCGTATCGTAGTTATCTACACGACGGGGAGTCAGGCAACTATGGATGAACGAAATAGACAGATCGCTGAGATAGGTGCCTCACTGATTAAGCATTGGTAACTGTCAGACCAAGTTTACTCATATATACTTTAGATTGATTTAAAACTTCATTTTTAATTTAAAAGGATCTAGGTGAAGATCCTTTTTGATAATCTCATGACCAAAATCCCTTAACGTGAGTTTTCGTTCCACTGAGCGTCAGACCCCGTAGAAAAGATCAAAGGATCTTCTTGAGATCCTTTTTTTCTGCGCGTAATCTGCTGCTTGCAAACAAAAAAACCACCGCTACCAGCGGTGGTTTGTTTGCCGGATCAAGAGCTACCAACTCTTTTTCCGAAGGTAACTGGCTTCAGCAGAGCGCAGATACCAAATACTGTTCTTCTAGTGTAGCCGTAGTTAGGCCACCACTTCAAGAACTCTGTAGCACCGCCTACATACCTCGCTCTGCTAATCCTGTTACCAGTGGCTGCTGCCAGTGGCGATAAGTCGTGTCTTACCGGGTTGGACTCAAGACGATAGTTACCGGATAAGGCGCAGCGGTCGGGCTGAACGGGGGGTTCGTGCACACAGCCCAGCTTGGAGCGAACGACCTACACCGAACTGAGATACCTACAGCGTGAGCTATGAGAAAGCGCCACGCTTCCCGAAGAGAGAAAGGCGGACAGGTATCCGGTAAGCGGCAGGGTCGGAACAGGAGAGCGCACGAGGGAGCTTCCAGGGGGAAACGCCTGGTATCTTTATAGTCCTGTCGGGTTTCGCCACCTCTGACTTGAGCGTCGATTTTTGTGATGCTCGTCAGGGGGGCGGAGCCTATGGAAAAACGCCAGCAACGCGGCCTTTTTACGGTTCCTGGCCTTTTGCTGGCCTTTTGCTCACATGTTCTTTCCTGCGTTATCCCCTGATTCTGTGGATAACCGTATTACCGCCTTTGAGTGAGCTGATACCGCTCGCCGCAGCCGAACGACCGAGCGCAGCGAGTCAGTGAGCGAGGAAGCGGAAGAGCGCCCAATACGCAAACCGCCTCTCCCCGCGCGTTGGCCGATTCATTAATGCAGCTGGCACGACAGGTTTCCCGACTGGAAAGCGGGCAGTGAGCGCAACGCAATTAATGTGAGTTAGCTCACTCATTAGGCACCCCAGGCTTTACACTTTATGCTTCCGGCTCGTATGTTGTGTGGAATTGTGAGCGGATAACAATTTCACACAGGAAACAGCTATGACCATGATTACGCCAAGCGCGCAATTAACCCTCACTAAAGGGAACAAAAGCTGGAGCTGCAAGCTT

**> Mt-Dimer1-HI-NESS - VB241124-1215tdy - pLV[Exp]-Puro-hPGK>{Mt- Dimer1-HI-NESS}**

AATGTAGTCTTATGCAATACTCTTGTAGTCTTGCAACATGGTAACGATGAGTTAGCAACATGCCTTACAAGGAGAGAAAAAGCACCGTGCATGCCGATTGGTGGAAGTAAGGTGGTACGATCGTGCCTTATTAGGAAGGCAACAGACGGGTCTGACATGGATTGGACGAACCACTGAATTGCCGCATTGCAGAGATATTGTATTTAAGTGCCTAGCTCGATACATAAACGGGTCTCTCTGGTTAGACCAGATCTGAGCCTGGGAGCTCTCTGGCTAACTAGGGAACCCACTGCTTAAGCCTCAATAAAGCTTGCCTTGAGTGCTTCAAGTAGTGTGTGCCCGTCTGTTGTGTGACTCTGGTAACTAGAGATCCCTCAGACCCTTTTAGTCAGTGTGGAAAATCTCTAGCAGTGGCGCCCGAACAGGGACTTGAAAGCGAAAGGGAAACCAGAGGAGCTCTCTCGACGCAGGACTCGGCTTGCTGAAGCGCGCACGGCAAGAGGCGAGGGGCGGCGACTGGTGAGTACGCCAAAAATTTTGACTAGCGGAGGCTAGAAGGAGAGAGATGGGTGCGAGAGCGTCAGTATTAAGCGGGGGAGAATTAGATCGCGATGGGAAAAAATTCGGTTAAGGCCAGGGGGAAAGAAAAAATATAAATTAAAACATATAGTATGGGCAAGCAGGGAGCTAGAACGATTCGCAGTTAATCCTGGCCTGTTAGAAACATCAGAAGGCTGTAGACAAATACTGGGACAGCTACAACCATCCCTTCAGACAGGATCAGAAGAACTTAGATCATTATATAATACAGTAGCAACCCTCTATTGTGTGCATCAAAGGATAGAGATAAAAGACACCAAGGAAGCTTTAGACAAGATAGAGGAAGAGCAAAACAAAAGTAAGACCACCGCACAGCAAGCGGCCGCTGATCTTCAGACCTGGAGGAGGAGATATGAGGGACAATTGGAGAAGTGAATTATATAAATATAAAGTAGTAAAAATTGAACCATTAGGAGTAGCACCCACCAAGGCAAAGAGAAGAGTGGTGCAGAGAGAAAAAAGAGCAGTGGGAATAGGAGCTTTGTTCCTTGGGTTCTTGGGAGCAGCAGGAAGCACTATGGGCGCAGCGTCAATGACGCTGACGGTACAGGCCAGACAATTATTGTCTGGTATAGTGCAGCAGCAGAACAATTTGCTGAGGGCTATTGAGGCGCAACAGCATCTGTTGCAACTCACAGTCTGGGGCATCAAGCAGCTCCAGGCAAGAATCCTGGCTGTGGAAAGATACCTAAAGGATCAACAGCTCCTGGGGATTTGGGGTTGCTCTGGAAAACTCATTTGCACCACTGCTGTGCCTTGGAATGCTAGTTGGAGTAATAAATCTCTGGAACAGATTTGGAATCACACGACCTGGATGGAGTGGGACAGAGAAATTAACAATTACACAAGCTTAATACACTCCTTAATTGAAGAATCGCAAAACCAGCAAGAAAAGAATGAACAAGAATTATTGGAATTAGATAAATGGGCAAGTTTGTGGAATTGGTTTAACATAACAAATTGGCTGTGGTATATAAAATTATTCATAATGATAGTAGGAGGCTTGGTAGGTTTAAGAATAGTTTTTGCTGTACTTTCTATAGTGAATAGAGTTAGGCAGGGATATTCACCATTATCGTTTCAGACCCACCTCCCAACCCCGAGGGGACCCGACAGGCCCGAAGGAATAGAAGAAGAAGGTGGAGAGAGAGACAGAGACAGATCCATTCGATTAGTGAACGGATCTCGACGGTATCGCTAGCTTTTAAAAGAAAAGGGGGGATTGGGGGGTACAGTGCAGGGGAAAGAATAGTAGACATAATAGCAACAGACATACAAACTAAAGAATTACAAAAACAAATTACAAAAATTCAAAATTTTACTAGTGATTATCGGATCAACTTTGTATAGAAAAGTTGGGGTTGCGCCTTTTCCAAGGCAGCCCTGGGTTTGCGCAGGGACGCGGCTGCTCTGGGCGTGGTTCCGGGAAACGCAGCGGCGCCGACCCTGGGTCTCGCACATTCTTCACGTCCGTTCGCAGCGTCACCCGGATCTTCGCCGCTACCCTTGTGGGCCCCCCGGCGACGCTTCCTGCTCCGCCCCTAAGTCGGGAAGGTTCCTTGCGGTTCGCGGCGTGCCGGACGTGACAAACGGAAGCCGCACGTCTCACTAGTACCCTCGCAGACGGACAGCGCCAGGGAGCAATGGCAGCGCGCCGACCGCGATGGGCTGTGGCCAATAGCGGCTGCTCAGCAGGGCGCGCCGAGAGCAGCGGCCGGGAAGGGGCGGTGCGGGAGGCGGGGTGTGGGGCGGTAGTGTGGGCCCTGTTCCTGCCCGCGCGGTGTTCCGCATTCTGCAAGCCTCCGGAGCGCACGTCGGCAGTCGGCTCCCTCGTTGACCGAATCACCGACCTCTCTCCCCAGGCAAGTTTGTACAAAAAAGCAGGCTGCCACCATGTCCGTCCTGACGCCGCTGCTGCTGCGGGGCTTGACAGGCTCGGCCCGGCGGCTCCCAGTGCCGCGCGCCAAGATCCATTCGTTGGGGGATCTGTCCGTCCTGACGCCGCTGCTGCTGCGGGGCTTGACAGGCTCGGCCCGGCGGCTCCCAGTGCCGCGCGCCAAGATCCATTCGTTGGGGGATCCACCGGTCGCCACCATGGTGGCCTCCTCCGAGGACGTCATCAAAGAGTTCATGCGCTTCAAGGTGCGCATGGAGGGCTCCGTGAACGGCCACGAGTTCGAGATCGAGGGCGAGGGCGAGGGCCGCCCCTACGAGGGCACCCAGACCGCCAAGCTGAAGGTGACCAAGGGCGGCCCCCTGCCCTTCGCCTGGGACATCCTGTCCCCCCAGTTCCAGTACGGCTCCAAGGTGTACGTGAAGCACCCCGCCGACATCCCCGACTACAAGAAGCTGTCCTTCCCCGAGGGCTTCAAGTGGGAGCGCGTGATGAACTTCGAGGACGGCGGCGTGGTGACCGTGACCCAGGACTCCTCCCTGCAGGACGGCTGCTTTATCTACAAGGTGAAGTTCCGCGGCGTGAACTTCCCCTCCGACGGCCCCGTAATGCAGAAGAAGACCATGGGCTGGGAGGCCTCCACCGAGCGCCTGTACCCCCGCGACGGCGTGCTGAAGGGCGAGATCCACCAGGCCCTGAAGCTGAAGGACGGCGGCCACTACCTGGTGGAGTTCAAGACCATCTACATGGCCAAGAAGCCCGTGCAGCTGCCCGGCTACTACTACGTGGACACCAAGCTGGACATCACCTCCCACAACGAGGACTACACCATCGTGGAACAGTACGAGCGCGCCGAGGGCCGCCACCACCTGTTCCTGGCTGCCGTTAAATCTGGCACCAAAGCTAAACGTGCTCAGCGTCCGGCAAAATATAGCTACGTTGACGAAAACGGCGAAACTAAAACCTGGACTGGCCAAGGCCGTACTCCAGCTGTAATCAAAAAAGCAATGGATGAGCAAGGTAAATCCCTCGACGATTTCCTGATCAAGCAATACCCATACGATGTTCCAGATTACGCTTAAACCCAGCTTTCTTGTACAAAGTGGTGATAATCGAATTCCGATAATCAACCTCTGGATTACAAAATTTGTGAAAGATTGACTGGTATTCTTAACTATGTTGCTCCTTTTACGCTATGTGGATACGCTGCTTTAATGCCTTTGTATCATGCTATTGCTTCCCGTATGGCTTTCATTTTCTCCTCCTTGTATAAATCCTGGTTGCTGTCTCTTTATGAGGAGTTGTGGCCCGTTGTCAGGCAACGTGGCGTGGTGTGCACTGTGTTTGCTGACGCAACCCCCACTGGTTGGGGCATTGCCACCACCTGTCAGCTCCTTTCCGGGACTTTCGCTTTCCCCCTCCCTATTGCCACGGCGGAACTCATCGCCGCCTGCCTTGCCCGCTGCTGGACAGGGGCTCGGCTGTTGGGCACTGACAATTCCGTGGTGTTGTCGGGGAAGCTGACGTCCTTTCCATGGCTGCTCGCCTGTGTTGCCACCTGGATTCTGCGCGGGACGTCCTTCTGCTACGTCCCTTCGGCCCTCAATCCAGCGGACCTTCCTTCCCGCGGCCTGCTGCCGGCTCTGCGGCCTCTTCCGCGTCTTCGCCTTCGCCCTCAGACGAGTCGGATCTCCCTTTGGGCCGCCTCCCCGCATCGGGAATTCCCGCGGTTCGAATTCTACCGGGTAGGGGAGGCGCTTTTCCCAAGGCAGTCTGGAGCATGCGCTTTAGCAGCCCCGCTGGGCACTTGGCGCTACACAAGTGGCCTCTGGCCTCGCACACATTCCACATCCACCGGTAGGCGCCAACCGGCTCCGTTCTTTGGTGGCCCCTTCGCGCCACCTTCTACTCCTCCCCTAGTCAGGAAGTTCCCCCCCGCCCCGCAGCTCGCGTCGTGCAGGACGTGACAAATGGAAGTAGCACGTCTCACTAGTCTCGTGCAGATGGACAGCACCGCTGAGCAATGGAAGCGGGTAGGCCTTTGGGGCAGCGGCCAATAGCAGCTTTGCTCCTTCGCTTTCTGGGCTCAGAGGCTGGGAAGGGGTGGGTCCGGGGGCGGGCTCAGGGGCGGGCTCAGGGGCGGGGCGGGCGCCCGAAGGTCCTCCGGAGGCCCGGCATTCTGCACGCTTCAAAAGCGCACGTCTGCCGCGCTGTTCTCCTCTTCCTCATCTCCGGGCCTTTCGACCTCACGTGGCCACCATGACCGAGTACAAGCCCACGGTGCGCCTCGCCACCCGCGACGACGTCCCCAGGGCCGTACGCACCCTCGCCGCCGCGTTCGCCGACTACCCCGCCACGCGCCACACCGTCGATCCGGACCGCCACATCGAGCGGGTCACCGAGCTGCAAGAACTCTTCCTCACGCGCGTCGGGCTCGACATCGGCAAGGTGTGGGTCGCGGACGACGGCGCCGCGGTGGCGGTCTGGACCACGCCGGAGAGCGTCGAAGCGGGGGCGGTGTTCGCCGAGATCGGCCCGCGCATGGCCGAGTTGAGCGGTTCCCGGCTGGCCGCGCAGCAACAGATGGAAGGCCTCCTGGCGCCGCACCGGCCCAAGGAGCCCGCGTGGTTCCTGGCCACCGTCGGCGTCTCGCCCGACCACCAGGGCAAGGGTCTGGGCAGCGCCGTCGTGCTCCCCGGAGTGGAGGCGGCCGAGCGCGCCGGGGTGCCCGCCTTCCTGGAGACCTCCGCGCCCCGCAACCTCCCCTTCTACGAGCGGCTCGGCTTCACCGTCACCGCCGACGTCGAGGTGCCCGAAGGACCGCGCACCTGGTGCATGACCCGCAAGCCCGGTGCCTGAGGTACCTTTAAGACCAATGACTTACAAGGCAGCTGTAGATCTTAGCCACTTTTTAAAAGAAAAGGGGGGACTGGAAGGGCTAATTCACTCCCAACGAAGACAAGATCTGCTTTTTGCTTGTACTGGGTCTCTCTGGTTAGACCAGATCTGAGCCTGGGAGCTCTCTGGCTAACTAGGGAACCCACTGCTTAAGCCTCAATAAAGCTTGCCTTGAGTGCTTCAAGTAGTGTGTGCCCGTCTGTTGTGTGACTCTGGTAACTAGAGATCCCTCAGACCCTTTTAGTCAGTGTGGAAAATCTCTAGCAGTAGTAGTTCATGTCATCTTATTATTCAGTATTTATAACTTGCAAAGAAATGAATATCAGAGAGTGAGAGGAACTTGTTTATTGCAGCTTATAATGGTTACAAATAAAGCAATAGCATCACAAATTTCACAAATAAAGCATTTTTTTCACTGCATTCTAGTTGTGGTTTGTCCAAACTCATCAATGTATCTTATCATGTCTGGCTCTAGCTATCCCGCCCCTAACTCCGCCCATCCCGCCCCTAACTCCGCCCAGTTCCGCCCATTCTCCGCCCCATGGCTGACTAATTTTTTTTATTTATGCAGAGGCCGAGGCCGCCTCGGCCTCTGAGCTATTCCAGAAGTAGTGAGGAGGCTTTTTTGGAGGCCTAGGGACGTACCCAATTCGCCCTATAGTGAGTCGTATTACGCGCGCTCACTGGCCGTCGTTTTACAACGTCGTGACTGGGAAAACCCTGGCGTTACCCAACTTAATCGCCTTGCAGCACATCCCCCTTTCGCCAGCTGGCGTAATAGCGAAGAGGCCCGCACCGATCGCCCTTCCCAACAGTTGCGCAGCCTGAATGGCGAATGGGACGCGCCCTGTAGCGGCGCATTAAGCGCGGCGGGTGTGGTGGTTACGCGCAGCGTGACCGCTACACTTGCCAGCGCCCTAGCGCCCGCTCCTTTCGCTTTCTTCCCTTCCTTTCTCGCCACGTTCGCCGGCTTTCCCCGTCAAGCTCTAAATCGGGGGCTCCCTTTAGGGTTCCGATTTAGTGCTTTACGGCACCTCGACCCCAAAAAACTTGATTAGGGTGATGGTTCACGTAGTGGGCCATCGCCCTGATAGACGGTTTTTCGCCCTTTGACGTTGGAGTCCACGTTCTTTAATAGTGGACTCTTGTTCCAAACTGGAACAACACTCAACCCTATCTCGGTCTATTCTTTTGATTTATAAGGGATTTTGCCGATTTCGGCCTATTGGTTAAAAAATGAGCTGATTTAACAAAAATTTAACGCGAATTTTAACAAAATATTAACGCTTACAATTTAGGTGGCACTTTTCGGGGAAATGTGCGCGGAACCCCTATTTGTTTATTTTTCTAAATACATTCAAATATGTATCCGCTCATGAGACAATAACCCTGATAAATGCTTCAATAATATTGAAAAAGGAAGAGTATGAGTATTCAACATTTCCGTGTCGCCCTTATTCCCTTTTTTGCGGCATTTTGCCTTCCTGTTTTTGCTCACCCAGAAACGCTGGTGAAAGTAAAAGATGCTGAAGATCAGTTGGGTGCACGAGTGGGTTACATCGAACTGGATCTCAACAGCGGTAAGATCCTTGAGAGTTTTCGCCCCGAAGAACGTTTTCCAATGATGAGCACTTTTAAAGTTCTGCTATGTGGCGCGGTATTATCCCGTATTGACGCCGGGCAAGAGCAACTCGGTCGCCGCATACACTATTCTCAGAATGACTTGGTTGAGTACTCACCAGTCACAGAAAAGCATCTTACGGATGGCATGACAGTAAGAGAATTATGCAGTGCTGCCATAACCATGAGTGATAACACTGCGGCCAACTTACTTCTGACAACGATCGGAGGACCGAAGGAGCTAACCGCTTTTTTGCACAACATGGGGGATCATGTAACTCGCCTTGATCGTTGGGAACCGGAGCTGAATGAAGCCATACCAAACGACGAGCGTGACACCACGATGCCTGTAGCAATGGCAACAACGTTGCGCAAACTATTAACTGGCGAACTACTTACTCTAGCTTCCCGGCAACAATTAATAGACTGGATGGAGGCGGATAAAGTTGCAGGACCACTTCTGCGCTCGGCCCTTCCGGCTGGCTGGTTTATTGCTGATAAATCTGGAGCCGGTGAGCGTGGGTCTCGCGGTATCATTGCAGCACTGGGGCCAGATGGTAAGCCCTCCCGTATCGTAGTTATCTACACGACGGGGAGTCAGGCAACTATGGATGAACGAAATAGACAGATCGCTGAGATAGGTGCCTCACTGATTAAGCATTGGTAACTGTCAGACCAAGTTTACTCATATATACTTTAGATTGATTTAAAACTTCATTTTTAATTTAAAAGGATCTAGGTGAAGATCCTTTTTGATAATCTCATGACCAAAATCCCTTAACGTGAGTTTTCGTTCCACTGAGCGTCAGACCCCGTAGAAAAGATCAAAGGATCTTCTTGAGATCCTTTTTTTCTGCGCGTAATCTGCTGCTTGCAAACAAAAAAACCACCGCTACCAGCGGTGGTTTGTTTGCCGGATCAAGAGCTACCAACTCTTTTTCCGAAGGTAACTGGCTTCAGCAGAGCGCAGATACCAAATACTGTTCTTCTAGTGTAGCCGTAGTTAGGCCACCACTTCAAGAACTCTGTAGCACCGCCTACATACCTCGCTCTGCTAATCCTGTTACCAGTGGCTGCTGCCAGTGGCGATAAGTCGTGTCTTACCGGGTTGGACTCAAGACGATAGTTACCGGATAAGGCGCAGCGGTCGGGCTGAACGGGGGGTTCGTGCACACAGCCCAGCTTGGAGCGAACGACCTACACCGAACTGAGATACCTACAGCGTGAGCTATGAGAAAGCGCCACGCTTCCCGAAGAGAGAAAGGCGGACAGGTATCCGGTAAGCGGCAGGGTCGGAACAGGAGAGCGCACGAGGGAGCTTCCAGGGGGAAACGCCTGGTATCTTTATAGTCCTGTCGGGTTTCGCCACCTCTGACTTGAGCGTCGATTTTTGTGATGCTCGTCAGGGGGGCGGAGCCTATGGAAAAACGCCAGCAACGCGGCCTTTTTACGGTTCCTGGCCTTTTGCTGGCCTTTTGCTCACATGTTCTTTCCTGCGTTATCCCCTGATTCTGTGGATAACCGTATTACCGCCTTTGAGTGAGCTGATACCGCTCGCCGCAGCCGAACGACCGAGCGCAGCGAGTCAGTGAGCGAGGAAGCGGAAGAGCGCCCAATACGCAAACCGCCTCTCCCCGCGCGTTGGCCGATTCATTAATGCAGCTGGCACGACAGGTTTCCCGACTGGAAAGCGGGCAGTGAGCGCAACGCAATTAATGTGAGTTAGCTCACTCATTAGGCACCCCAGGCTTTACACTTTATGCTTCCGGCTCGTATGTTGTGTGGAATTGTGAGCGGATAACAATTTCACACAGGAAACAGCTATGACCATGATTACGCCAAGCGCGCAATTAACCCTCACTAAAGGGAACAAAAGCTGGAGCTGCAAGCTT

**> Mt-DsRED-Express2-HI-NESS - VB240525-1016avy - pLV[Exp]-Puro-hPGK>{Mt- DsRED-Express2-HI-NESS}**

AATGTAGTCTTATGCAATACTCTTGTAGTCTTGCAACATGGTAACGATGAGTTAGCAACATGCCTTACAAGGAGAGAAAAAGCACCGTGCATGCCGATTGGTGGAAGTAAGGTGGTACGATCGTGCCTTATTAGGAAGGCAACAGACGGGTCTGACATGGATTGGACGAACCACTGAATTGCCGCATTGCAGAGATATTGTATTTAAGTGCCTAGCTCGATACATAAACGGGTCTCTCTGGTTAGACCAGATCTGAGCCTGGGAGCTCTCTGGCTAACTAGGGAACCCACTGCTTAAGCCTCAATAAAGCTTGCCTTGAGTGCTTCAAGTAGTGTGTGCCCGTCTGTTGTGTGACTCTGGTAACTAGAGATCCCTCAGACCCTTTTAGTCAGTGTGGAAAATCTCTAGCAGTGGCGCCCGAACAGGGACTTGAAAGCGAAAGGGAAACCAGAGGAGCTCTCTCGACGCAGGACTCGGCTTGCTGAAGCGCGCACGGCAAGAGGCGAGGGGCGGCGACTGGTGAGTACGCCAAAAATTTTGACTAGCGGAGGCTAGAAGGAGAGAGATGGGTGCGAGAGCGTCAGTATTAAGCGGGGGAGAATTAGATCGCGATGGGAAAAAATTCGGTTAAGGCCAGGGGGAAAGAAAAAATATAAATTAAAACATATAGTATGGGCAAGCAGGGAGCTAGAACGATTCGCAGTTAATCCTGGCCTGTTAGAAACATCAGAAGGCTGTAGACAAATACTGGGACAGCTACAACCATCCCTTCAGACAGGATCAGAAGAACTTAGATCATTATATAATACAGTAGCAACCCTCTATTGTGTGCATCAAAGGATAGAGATAAAAGACACCAAGGAAGCTTTAGACAAGATAGAGGAAGAGCAAAACAAAAGTAAGACCACCGCACAGCAAGCGGCCGCTGATCTTCAGACCTGGAGGAGGAGATATGAGGGACAATTGGAGAAGTGAATTATATAAATATAAAGTAGTAAAAATTGAACCATTAGGAGTAGCACCCACCAAGGCAAAGAGAAGAGTGGTGCAGAGAGAAAAAAGAGCAGTGGGAATAGGAGCTTTGTTCCTTGGGTTCTTGGGAGCAGCAGGAAGCACTATGGGCGCAGCGTCAATGACGCTGACGGTACAGGCCAGACAATTATTGTCTGGTATAGTGCAGCAGCAGAACAATTTGCTGAGGGCTATTGAGGCGCAACAGCATCTGTTGCAACTCACAGTCTGGGGCATCAAGCAGCTCCAGGCAAGAATCCTGGCTGTGGAAAGATACCTAAAGGATCAACAGCTCCTGGGGATTTGGGGTTGCTCTGGAAAACTCATTTGCACCACTGCTGTGCCTTGGAATGCTAGTTGGAGTAATAAATCTCTGGAACAGATTTGGAATCACACGACCTGGATGGAGTGGGACAGAGAAATTAACAATTACACAAGCTTAATACACTCCTTAATTGAAGAATCGCAAAACCAGCAAGAAAAGAATGAACAAGAATTATTGGAATTAGATAAATGGGCAAGTTTGTGGAATTGGTTTAACATAACAAATTGGCTGTGGTATATAAAATTATTCATAATGATAGTAGGAGGCTTGGTAGGTTTAAGAATAGTTTTTGCTGTACTTTCTATAGTGAATAGAGTTAGGCAGGGATATTCACCATTATCGTTTCAGACCCACCTCCCAACCCCGAGGGGACCCGACAGGCCCGAAGGAATAGAAGAAGAAGGTGGAGAGAGAGACAGAGACAGATCCATTCGATTAGTGAACGGATCTCGACGGTATCGCTAGCTTTTAAAAGAAAAGGGGGGATTGGGGGGTACAGTGCAGGGGAAAGAATAGTAGACATAATAGCAACAGACATACAAACTAAAGAATTACAAAAACAAATTACAAAAATTCAAAATTTTACTAGTGATTATCGGATCAACTTTGTATAGAAAAGTTGGGGTTGCGCCTTTTCCAAGGCAGCCCTGGGTTTGCGCAGGGACGCGGCTGCTCTGGGCGTGGTTCCGGGAAACGCAGCGGCGCCGACCCTGGGTCTCGCACATTCTTCACGTCCGTTCGCAGCGTCACCCGGATCTTCGCCGCTACCCTTGTGGGCCCCCCGGCGACGCTTCCTGCTCCGCCCCTAAGTCGGGAAGGTTCCTTGCGGTTCGCGGCGTGCCGGACGTGACAAACGGAAGCCGCACGTCTCACTAGTACCCTCGCAGACGGACAGCGCCAGGGAGCAATGGCAGCGCGCCGACCGCGATGGGCTGTGGCCAATAGCGGCTGCTCAGCAGGGCGCGCCGAGAGCAGCGGCCGGGAAGGGGCGGTGCGGGAGGCGGGGTGTGGGGCGGTAGTGTGGGCCCTGTTCCTGCCCGCGCGGTGTTCCGCATTCTGCAAGCCTCCGGAGCGCACGTCGGCAGTCGGCTCCCTCGTTGACCGAATCACCGACCTCTCTCCCCAGGCAAGTTTGTACAAAAAAGCAGGCTGCCACCATGTCCGTCCTGACGCCGCTGCTGCTGCGGGGCTTGACAGGCTCGGCCCGGCGGCTCCCAGTGCCGCGCGCCAAGATCCATTCGTTGGGGGATCTGTCCGTCCTGACGCCGCTGCTGCTGCGGGGCTTGACAGGCTCGGCCCGGCGGCTCCCAGTGCCGCGCGCCAAGATCCATTCGTTGGGGGATCCACCGGTCGCCACCATGGATAGCACTGAGAACGTCATCAAGCCCTTCATGCGCTTCAAGGTGCACATGGAGGGCTCCGTGAACGGCCACGAGTTCGAGATCGAGGGCGAGGGCGAGGGCAAGCCCTACGAGGGCACCCAGACCGCCAAGCTGCAGGTGACCAAGGGCGGCCCCCTGCCCTTCGCCTGGGACATCCTGTCCCCCCAGTTCCAGTACGGCTCCAAGGTGTACGTGAAGCACCCCGCCGACATCCCCGACTACAAGAAGCTGTCCTTCCCCGAGGGCTTCAAGTGGGAGCGCGTGATGAACTTCGAGGACGGCGGCGTGGTGACCGTGACCCAGGACTCCTCCCTGCAGGACGGCACCTTCATCTACCACGTGAAGTTCATCGGCGTGAACTTCCCCTCCGACGGCCCCGTAATGCAGAAGAAGACTCTGGGCTGGGAGCCCTCCACCGAGCGCCTGTACCCCCGCGACGGCGTGCTGAAGGGCGAGATCCACAAGGCGCTGAAGCTGAAGGGCGGCGGCCACTACCTGGTGGAGTTCAAGTCAATCTACATGGCCAAGAAGCCCGTGAAGCTGCCCGGCTACTACTACGTGGACTCCAAGCTGGACATCACCTCCCACAACGAGGACTACACCGTGGTGGAGCAGTACGAGCGCGCCGAGGCCCGCCACCACCTGTTCCAGGCTGCCGTTAAATCTGGCACCAAAGCTAAACGTGCTCAGCGTCCGGCAAAATATAGCTACGTTGACGAAAACGGCGAAACTAAAACCTGGACTGGCCAAGGCCGTACTCCAGCTGTAATCAAAAAAGCAATGGATGAGCAAGGTAAATCCCTCGACGATTTCCTGATCAAGCAATACCCATACGATGTTCCAGATTACGCTTAAACCCAGCTTTCTTGTACAAAGTGGTGATAATCGAATTCCGATAATCAACCTCTGGATTACAAAATTTGTGAAAGATTGACTGGTATTCTTAACTATGTTGCTCCTTTTACGCTATGTGGATACGCTGCTTTAATGCCTTTGTATCATGCTATTGCTTCCCGTATGGCTTTCATTTTCTCCTCCTTGTATAAATCCTGGTTGCTGTCTCTTTATGAGGAGTTGTGGCCCGTTGTCAGGCAACGTGGCGTGGTGTGCACTGTGTTTGCTGACGCAACCCCCACTGGTTGGGGCATTGCCACCACCTGTCAGCTCCTTTCCGGGACTTTCGCTTTCCCCCTCCCTATTGCCACGGCGGAACTCATCGCCGCCTGCCTTGCCCGCTGCTGGACAGGGGCTCGGCTGTTGGGCACTGACAATTCCGTGGTGTTGTCGGGGAAGCTGACGTCCTTTCCATGGCTGCTCGCCTGTGTTGCCACCTGGATTCTGCGCGGGACGTCCTTCTGCTACGTCCCTTCGGCCCTCAATCCAGCGGACCTTCCTTCCCGCGGCCTGCTGCCGGCTCTGCGGCCTCTTCCGCGTCTTCGCCTTCGCCCTCAGACGAGTCGGATCTCCCTTTGGGCCGCCTCCCCGCATCGGGAATTCCCGCGGTTCGAATTCTACCGGGTAGGGGAGGCGCTTTTCCCAAGGCAGTCTGGAGCATGCGCTTTAGCAGCCCCGCTGGGCACTTGGCGCTACACAAGTGGCCTCTGGCCTCGCACACATTCCACATCCACCGGTAGGCGCCAACCGGCTCCGTTCTTTGGTGGCCCCTTCGCGCCACCTTCTACTCCTCCCCTAGTCAGGAAGTTCCCCCCCGCCCCGCAGCTCGCGTCGTGCAGGACGTGACAAATGGAAGTAGCACGTCTCACTAGTCTCGTGCAGATGGACAGCACCGCTGAGCAATGGAAGCGGGTAGGCCTTTGGGGCAGCGGCCAATAGCAGCTTTGCTCCTTCGCTTTCTGGGCTCAGAGGCTGGGAAGGGGTGGGTCCGGGGGCGGGCTCAGGGGCGGGCTCAGGGGCGGGGCGGGCGCCCGAAGGTCCTCCGGAGGCCCGGCATTCTGCACGCTTCAAAAGCGCACGTCTGCCGCGCTGTTCTCCTCTTCCTCATCTCCGGGCCTTTCGACCTCACGTGGCCACCATGACCGAGTACAAGCCCACGGTGCGCCTCGCCACCCGCGACGACGTCCCCAGGGCCGTACGCACCCTCGCCGCCGCGTTCGCCGACTACCCCGCCACGCGCCACACCGTCGATCCGGACCGCCACATCGAGCGGGTCACCGAGCTGCAAGAACTCTTCCTCACGCGCGTCGGGCTCGACATCGGCAAGGTGTGGGTCGCGGACGACGGCGCCGCGGTGGCGGTCTGGACCACGCCGGAGAGCGTCGAAGCGGGGGCGGTGTTCGCCGAGATCGGCCCGCGCATGGCCGAGTTGAGCGGTTCCCGGCTGGCCGCGCAGCAACAGATGGAAGGCCTCCTGGCGCCGCACCGGCCCAAGGAGCCCGCGTGGTTCCTGGCCACCGTCGGCGTCTCGCCCGACCACCAGGGCAAGGGTCTGGGCAGCGCCGTCGTGCTCCCCGGAGTGGAGGCGGCCGAGCGCGCCGGGGTGCCCGCCTTCCTGGAGACCTCCGCGCCCCGCAACCTCCCCTTCTACGAGCGGCTCGGCTTCACCGTCACCGCCGACGTCGAGGTGCCCGAAGGACCGCGCACCTGGTGCATGACCCGCAAGCCCGGTGCCTGAGGTACCTTTAAGACCAATGACTTACAAGGCAGCTGTAGATCTTAGCCACTTTTTAAAAGAAAAGGGGGGACTGGAAGGGCTAATTCACTCCCAACGAAGACAAGATCTGCTTTTTGCTTGTACTGGGTCTCTCTGGTTAGACCAGATCTGAGCCTGGGAGCTCTCTGGCTAACTAGGGAACCCACTGCTTAAGCCTCAATAAAGCTTGCCTTGAGTGCTTCAAGTAGTGTGTGCCCGTCTGTTGTGTGACTCTGGTAACTAGAGATCCCTCAGACCCTTTTAGTCAGTGTGGAAAATCTCTAGCAGTAGTAGTTCATGTCATCTTATTATTCAGTATTTATAACTTGCAAAGAAATGAATATCAGAGAGTGAGAGGAACTTGTTTATTGCAGCTTATAATGGTTACAAATAAAGCAATAGCATCACAAATTTCACAAATAAAGCATTTTTTTCACTGCATTCTAGTTGTGGTTTGTCCAAACTCATCAATGTATCTTATCATGTCTGGCTCTAGCTATCCCGCCCCTAACTCCGCCCATCCCGCCCCTAACTCCGCCCAGTTCCGCCCATTCTCCGCCCCATGGCTGACTAATTTTTTTTATTTATGCAGAGGCCGAGGCCGCCTCGGCCTCTGAGCTATTCCAGAAGTAGTGAGGAGGCTTTTTTGGAGGCCTAGGGACGTACCCAATTCGCCCTATAGTGAGTCGTATTACGCGCGCTCACTGGCCGTCGTTTTACAACGTCGTGACTGGGAAAACCCTGGCGTTACCCAACTTAATCGCCTTGCAGCACATCCCCCTTTCGCCAGCTGGCGTAATAGCGAAGAGGCCCGCACCGATCGCCCTTCCCAACAGTTGCGCAGCCTGAATGGCGAATGGGACGCGCCCTGTAGCGGCGCATTAAGCGCGGCGGGTGTGGTGGTTACGCGCAGCGTGACCGCTACACTTGCCAGCGCCCTAGCGCCCGCTCCTTTCGCTTTCTTCCCTTCCTTTCTCGCCACGTTCGCCGGCTTTCCCCGTCAAGCTCTAAATCGGGGGCTCCCTTTAGGGTTCCGATTTAGTGCTTTACGGCACCTCGACCCCAAAAAACTTGATTAGGGTGATGGTTCACGTAGTGGGCCATCGCCCTGATAGACGGTTTTTCGCCCTTTGACGTTGGAGTCCACGTTCTTTAATAGTGGACTCTTGTTCCAAACTGGAACAACACTCAACCCTATCTCGGTCTATTCTTTTGATTTATAAGGGATTTTGCCGATTTCGGCCTATTGGTTAAAAAATGAGCTGATTTAACAAAAATTTAACGCGAATTTTAACAAAATATTAACGCTTACAATTTAGGTGGCACTTTTCGGGGAAATGTGCGCGGAACCCCTATTTGTTTATTTTTCTAAATACATTCAAATATGTATCCGCTCATGAGACAATAACCCTGATAAATGCTTCAATAATATTGAAAAAGGAAGAGTATGAGTATTCAACATTTCCGTGTCGCCCTTATTCCCTTTTTTGCGGCATTTTGCCTTCCTGTTTTTGCTCACCCAGAAACGCTGGTGAAAGTAAAAGATGCTGAAGATCAGTTGGGTGCACGAGTGGGTTACATCGAACTGGATCTCAACAGCGGTAAGATCCTTGAGAGTTTTCGCCCCGAAGAACGTTTTCCAATGATGAGCACTTTTAAAGTTCTGCTATGTGGCGCGGTATTATCCCGTATTGACGCCGGGCAAGAGCAACTCGGTCGCCGCATACACTATTCTCAGAATGACTTGGTTGAGTACTCACCAGTCACAGAAAAGCATCTTACGGATGGCATGACAGTAAGAGAATTATGCAGTGCTGCCATAACCATGAGTGATAACACTGCGGCCAACTTACTTCTGACAACGATCGGAGGACCGAAGGAGCTAACCGCTTTTTTGCACAACATGGGGGATCATGTAACTCGCCTTGATCGTTGGGAACCGGAGCTGAATGAAGCCATACCAAACGACGAGCGTGACACCACGATGCCTGTAGCAATGGCAACAACGTTGCGCAAACTATTAACTGGCGAACTACTTACTCTAGCTTCCCGGCAACAATTAATAGACTGGATGGAGGCGGATAAAGTTGCAGGACCACTTCTGCGCTCGGCCCTTCCGGCTGGCTGGTTTATTGCTGATAAATCTGGAGCCGGTGAGCGTGGGTCTCGCGGTATCATTGCAGCACTGGGGCCAGATGGTAAGCCCTCCCGTATCGTAGTTATCTACACGACGGGGAGTCAGGCAACTATGGATGAACGAAATAGACAGATCGCTGAGATAGGTGCCTCACTGATTAAGCATTGGTAACTGTCAGACCAAGTTTACTCATATATACTTTAGATTGATTTAAAACTTCATTTTTAATTTAAAAGGATCTAGGTGAAGATCCTTTTTGATAATCTCATGACCAAAATCCCTTAACGTGAGTTTTCGTTCCACTGAGCGTCAGACCCCGTAGAAAAGATCAAAGGATCTTCTTGAGATCCTTTTTTTCTGCGCGTAATCTGCTGCTTGCAAACAAAAAAACCACCGCTACCAGCGGTGGTTTGTTTGCCGGATCAAGAGCTACCAACTCTTTTTCCGAAGGTAACTGGCTTCAGCAGAGCGCAGATACCAAATACTGTTCTTCTAGTGTAGCCGTAGTTAGGCCACCACTTCAAGAACTCTGTAGCACCGCCTACATACCTCGCTCTGCTAATCCTGTTACCAGTGGCTGCTGCCAGTGGCGATAAGTCGTGTCTTACCGGGTTGGACTCAAGACGATAGTTACCGGATAAGGCGCAGCGGTCGGGCTGAACGGGGGGTTCGTGCACACAGCCCAGCTTGGAGCGAACGACCTACACCGAACTGAGATACCTACAGCGTGAGCTATGAGAAAGCGCCACGCTTCCCGAAGAGAGAAAGGCGGACAGGTATCCGGTAAGCGGCAGGGTCGGAACAGGAGAGCGCACGAGGGAGCTTCCAGGGGGAAACGCCTGGTATCTTTATAGTCCTGTCGGGTTTCGCCACCTCTGACTTGAGCGTCGATTTTTGTGATGCTCGTCAGGGGGGCGGAGCCTATGGAAAAACGCCAGCAACGCGGCCTTTTTACGGTTCCTGGCCTTTTGCTGGCCTTTTGCTCACATGTTCTTTCCTGCGTTATCCCCTGATTCTGTGGATAACCGTATTACCGCCTTTGAGTGAGCTGATACCGCTCGCCGCAGCCGAACGACCGAGCGCAGCGAGTCAGTGAGCGAGGAAGCGGAAGAGCGCCCAATACGCAAACCGCCTCTCCCCGCGCGTTGGCCGATTCATTAATGCAGCTGGCACGACAGGTTTCCCGACTGGAAAGCGGGCAGTGAGCGCAACGCAATTAATGTGAGTTAGCTCACTCATTAGGCACCCCAGGCTTTACACTTTATGCTTCCGGCTCGTATGTTGTGTGGAATTGTGAGCGGATAACAATTTCACACAGGAAACAGCTATGACCATGATTACGCCAAGCGCGCAATTAACCCTCACTAAAGGGAACAAAAGCTGGAGCTGCAAGCTT

**> Mt-TagRFP-N/C-HI-NESS - VB230529-1292kdj - pLV[Exp]-Puro-hPGK>{Mt- TagRFP-N/C -HI-NESS}**

AATGTAGTCTTATGCAATACTCTTGTAGTCTTGCAACATGGTAACGATGAGTTAGCAACATGCCTTACAAGGAGAGAAAAAGCACCGTGCATGCCGATTGGTGGAAGTAAGGTGGTACGATCGTGCCTTATTAGGAAGGCAACAGACGGGTCTGACATGGATTGGACGAACCACTGAATTGCCGCATTGCAGAGATATTGTATTTAAGTGCCTAGCTCGATACATAAACGGGTCTCTCTGGTTAGACCAGATCTGAGCCTGGGAGCTCTCTGGCTAACTAGGGAACCCACTGCTTAAGCCTCAATAAAGCTTGCCTTGAGTGCTTCAAGTAGTGTGTGCCCGTCTGTTGTGTGACTCTGGTAACTAGAGATCCCTCAGACCCTTTTAGTCAGTGTGGAAAATCTCTAGCAGTGGCGCCCGAACAGGGACTTGAAAGCGAAAGGGAAACCAGAGGAGCTCTCTCGACGCAGGACTCGGCTTGCTGAAGCGCGCACGGCAAGAGGCGAGGGGCGGCGACTGGTGAGTACGCCAAAAATTTTGACTAGCGGAGGCTAGAAGGAGAGAGATGGGTGCGAGAGCGTCAGTATTAAGCGGGGGAGAATTAGATCGCGATGGGAAAAAATTCGGTTAAGGCCAGGGGGAAAGAAAAAATATAAATTAAAACATATAGTATGGGCAAGCAGGGAGCTAGAACGATTCGCAGTTAATCCTGGCCTGTTAGAAACATCAGAAGGCTGTAGACAAATACTGGGACAGCTACAACCATCCCTTCAGACAGGATCAGAAGAACTTAGATCATTATATAATACAGTAGCAACCCTCTATTGTGTGCATCAAAGGATAGAGATAAAAGACACCAAGGAAGCTTTAGACAAGATAGAGGAAGAGCAAAACAAAAGTAAGACCACCGCACAGCAAGCGGCCGCTGATCTTCAGACCTGGAGGAGGAGATATGAGGGACAATTGGAGAAGTGAATTATATAAATATAAAGTAGTAAAAATTGAACCATTAGGAGTAGCACCCACCAAGGCAAAGAGAAGAGTGGTGCAGAGAGAAAAAAGAGCAGTGGGAATAGGAGCTTTGTTCCTTGGGTTCTTGGGAGCAGCAGGAAGCACTATGGGCGCAGCGTCAATGACGCTGACGGTACAGGCCAGACAATTATTGTCTGGTATAGTGCAGCAGCAGAACAATTTGCTGAGGGCTATTGAGGCGCAACAGCATCTGTTGCAACTCACAGTCTGGGGCATCAAGCAGCTCCAGGCAAGAATCCTGGCTGTGGAAAGATACCTAAAGGATCAACAGCTCCTGGGGATTTGGGGTTGCTCTGGAAAACTCATTTGCACCACTGCTGTGCCTTGGAATGCTAGTTGGAGTAATAAATCTCTGGAACAGATTTGGAATCACACGACCTGGATGGAGTGGGACAGAGAAATTAACAATTACACAAGCTTAATACACTCCTTAATTGAAGAATCGCAAAACCAGCAAGAAAAGAATGAACAAGAATTATTGGAATTAGATAAATGGGCAAGTTTGTGGAATTGGTTTAACATAACAAATTGGCTGTGGTATATAAAATTATTCATAATGATAGTAGGAGGCTTGGTAGGTTTAAGAATAGTTTTTGCTGTACTTTCTATAGTGAATAGAGTTAGGCAGGGATATTCACCATTATCGTTTCAGACCCACCTCCCAACCCCGAGGGGACCCGACAGGCCCGAAGGAATAGAAGAAGAAGGTGGAGAGAGAGACAGAGACAGATCCATTCGATTAGTGAACGGATCTCGACGGTATCGCTAGCTTTTAAAAGAAAAGGGGGGATTGGGGGGTACAGTGCAGGGGAAAGAATAGTAGACATAATAGCAACAGACATACAAACTAAAGAATTACAAAAACAAATTACAAAAATTCAAAATTTTACTAGTGATTATCGGATCAACTTTGTATAGAAAAGTTGGGGTTGCGCCTTTTCCAAGGCAGCCCTGGGTTTGCGCAGGGACGCGGCTGCTCTGGGCGTGGTTCCGGGAAACGCAGCGGCGCCGACCCTGGGTCTCGCACATTCTTCACGTCCGTTCGCAGCGTCACCCGGATCTTCGCCGCTACCCTTGTGGGCCCCCCGGCGACGCTTCCTGCTCCGCCCCTAAGTCGGGAAGGTTCCTTGCGGTTCGCGGCGTGCCGGACGTGACAAACGGAAGCCGCACGTCTCACTAGTACCCTCGCAGACGGACAGCGCCAGGGAGCAATGGCAGCGCGCCGACCGCGATGGGCTGTGGCCAATAGCGGCTGCTCAGCAGGGCGCGCCGAGAGCAGCGGCCGGGAAGGGGCGGTGCGGGAGGCGGGGTGTGGGGCGGTAGTGTGGGCCCTGTTCCTGCCCGCGCGGTGTTCCGCATTCTGCAAGCCTCCGGAGCGCACGTCGGCAGTCGGCTCCCTCGTTGACCGAATCACCGACCTCTCTCCCCAGGCAAGTTTGTACAAAAAAGCAGGCTGCCACCATGTCCGTCCTGACGCCGCTGCTGCTGCGGGGCTTGACAGGCTCGGCCCGGCGGCTCCCAGTGCCGCGCGCCAAGATCCATTCGTTGGGGGATCTGTCCGTCCTGACGCCGCTGCTGCTGCGGGGCTTGACAGGCTCGGCCCGGCGGCTCCCAGTGCCGCGCGCCAAGGCCGCTGTGAAGTCCGGAACAAAGGCCAAGAGAGCCCAAAGACCCGCCAAGTACTCTTATGTGGATGAGAATGGAGAGACCAAGACATGGACCGGTCAGGGTAGAACACCTGCAGTGATTAAGAAGGCTATGGACGAACAGGGAAAGAGCCTGGATGACTTTCTCATTAAACAGATCCATTCGTTGGGGGATCCACCGGTCGCCACCATGAGCGAGCTGATTAAGGAGAACATGCACATGAAGCTGTACATGGAGGGCACCGTGAACAACCACCACTTCAAGTGCACATCCGAGGGCGAAGGCAAGCCCTACGAGGGCACCCAGACCATGAGAATCAAGGTGGTCGAGGGCGGCCCTCTCCCCTTCGCCTTCGACATCCTGGCTACCAGCTTCATGTACGGCAGCAGAACCTTCATCAACCACACCCAGGGCATCCCCGACTTCTTTAAGCAGTCCTTCCCTGAGGGCTTCACATGGGAGAGAGTCACCACATACGAAGACGGGGGCGTGCTGACCGCTACCCAGGACACCAGCCTCCAGGACGGCTGCCTCATCTACAACGTCAAGATCAGAGGGGTGAACTTCCCATCCAACGGCCCTGTGATGCAGAAGAAAACACTCGGCTGGGAGGCCAACACCGAGATGCTGTACCCCGCTGACGGCGGCCTGGAAGGCAGAAGCGACATGGCCCTGAAGCTCGTGGGCGGGGGCCACCTGATCTGCAACTTCAAGACCACATACAGATCCAAGAAACCCGCTAAGAACCTCAAGATGCCCGGCGTCTACTATGTGGACCACAGACTGGAAAGAATCAAGGAGGCCGACAAAGAGACCTACGTCGAGCAGCACGAGGTGGCTGTGGCCAGATACTGCGACCTCCCTAGCAAACTGGGGCACAAGGCTGCCGTTAAATCTGGCACCAAAGCTAAACGTGCTCAGCGTCCGGCAAAATATAGCTACGTTGACGAAAACGGCGAAACTAAAACCTGGACTGGCCAAGGCCGTACTCCAGCTGTAATCAAAAAAGCAATGGATGAGCAAGGTAAATCCCTCGACGATTTCCTGATCAAGCAATACCCATACGATGTTCCAGATTACGCTTAAACCCAGCTTTCTTGTACAAAGTGGTGATAATCGAATTCCGATAATCAACCTCTGGATTACAAAATTTGTGAAAGATTGACTGGTATTCTTAACTATGTTGCTCCTTTTACGCTATGTGGATACGCTGCTTTAATGCCTTTGTATCATGCTATTGCTTCCCGTATGGCTTTCATTTTCTCCTCCTTGTATAAATCCTGGTTGCTGTCTCTTTATGAGGAGTTGTGGCCCGTTGTCAGGCAACGTGGCGTGGTGTGCACTGTGTTTGCTGACGCAACCCCCACTGGTTGGGGCATTGCCACCACCTGTCAGCTCCTTTCCGGGACTTTCGCTTTCCCCCTCCCTATTGCCACGGCGGAACTCATCGCCGCCTGCCTTGCCCGCTGCTGGACAGGGGCTCGGCTGTTGGGCACTGACAATTCCGTGGTGTTGTCGGGGAAGCTGACGTCCTTTCCATGGCTGCTCGCCTGTGTTGCCACCTGGATTCTGCGCGGGACGTCCTTCTGCTACGTCCCTTCGGCCCTCAATCCAGCGGACCTTCCTTCCCGCGGCCTGCTGCCGGCTCTGCGGCCTCTTCCGCGTCTTCGCCTTCGCCCTCAGACGAGTCGGATCTCCCTTTGGGCCGCCTCCCCGCATCGGGAATTCCCGCGGTTCGAATTCTACCGGGTAGGGGAGGCGCTTTTCCCAAGGCAGTCTGGAGCATGCGCTTTAGCAGCCCCGCTGGGCACTTGGCGCTACACAAGTGGCCTCTGGCCTCGCACACATTCCACATCCACCGGTAGGCGCCAACCGGCTCCGTTCTTTGGTGGCCCCTTCGCGCCACCTTCTACTCCTCCCCTAGTCAGGAAGTTCCCCCCCGCCCCGCAGCTCGCGTCGTGCAGGACGTGACAAATGGAAGTAGCACGTCTCACTAGTCTCGTGCAGATGGACAGCACCGCTGAGCAATGGAAGCGGGTAGGCCTTTGGGGCAGCGGCCAATAGCAGCTTTGCTCCTTCGCTTTCTGGGCTCAGAGGCTGGGAAGGGGTGGGTCCGGGGGCGGGCTCAGGGGCGGGCTCAGGGGCGGGGCGGGCGCCCGAAGGTCCTCCGGAGGCCCGGCATTCTGCACGCTTCAAAAGCGCACGTCTGCCGCGCTGTTCTCCTCTTCCTCATCTCCGGGCCTTTCGACCTCACGTGGCCACCATGACCGAGTACAAGCCCACGGTGCGCCTCGCCACCCGCGACGACGTCCCCAGGGCCGTACGCACCCTCGCCGCCGCGTTCGCCGACTACCCCGCCACGCGCCACACCGTCGATCCGGACCGCCACATCGAGCGGGTCACCGAGCTGCAAGAACTCTTCCTCACGCGCGTCGGGCTCGACATCGGCAAGGTGTGGGTCGCGGACGACGGCGCCGCGGTGGCGGTCTGGACCACGCCGGAGAGCGTCGAAGCGGGGGCGGTGTTCGCCGAGATCGGCCCGCGCATGGCCGAGTTGAGCGGTTCCCGGCTGGCCGCGCAGCAACAGATGGAAGGCCTCCTGGCGCCGCACCGGCCCAAGGAGCCCGCGTGGTTCCTGGCCACCGTCGGCGTCTCGCCCGACCACCAGGGCAAGGGTCTGGGCAGCGCCGTCGTGCTCCCCGGAGTGGAGGCGGCCGAGCGCGCCGGGGTGCCCGCCTTCCTGGAGACCTCCGCGCCCCGCAACCTCCCCTTCTACGAGCGGCTCGGCTTCACCGTCACCGCCGACGTCGAGGTGCCCGAAGGACCGCGCACCTGGTGCATGACCCGCAAGCCCGGTGCCTGAGGTACCTTTAAGACCAATGACTTACAAGGCAGCTGTAGATCTTAGCCACTTTTTAAAAGAAAAGGGGGGACTGGAAGGGCTAATTCACTCCCAACGAAGACAAGATCTGCTTTTTGCTTGTACTGGGTCTCTCTGGTTAGACCAGATCTGAGCCTGGGAGCTCTCTGGCTAACTAGGGAACCCACTGCTTAAGCCTCAATAAAGCTTGCCTTGAGTGCTTCAAGTAGTGTGTGCCCGTCTGTTGTGTGACTCTGGTAACTAGAGATCCCTCAGACCCTTTTAGTCAGTGTGGAAAATCTCTAGCAGTAGTAGTTCATGTCATCTTATTATTCAGTATTTATAACTTGCAAAGAAATGAATATCAGAGAGTGAGAGGAACTTGTTTATTGCAGCTTATAATGGTTACAAATAAAGCAATAGCATCACAAATTTCACAAATAAAGCATTTTTTTCACTGCATTCTAGTTGTGGTTTGTCCAAACTCATCAATGTATCTTATCATGTCTGGCTCTAGCTATCCCGCCCCTAACTCCGCCCATCCCGCCCCTAACTCCGCCCAGTTCCGCCCATTCTCCGCCCCATGGCTGACTAATTTTTTTTATTTATGCAGAGGCCGAGGCCGCCTCGGCCTCTGAGCTATTCCAGAAGTAGTGAGGAGGCTTTTTTGGAGGCCTAGGGACGTACCCAATTCGCCCTATAGTGAGTCGTATTACGCGCGCTCACTGGCCGTCGTTTTACAACGTCGTGACTGGGAAAACCCTGGCGTTACCCAACTTAATCGCCTTGCAGCACATCCCCCTTTCGCCAGCTGGCGTAATAGCGAAGAGGCCCGCACCGATCGCCCTTCCCAACAGTTGCGCAGCCTGAATGGCGAATGGGACGCGCCCTGTAGCGGCGCATTAAGCGCGGCGGGTGTGGTGGTTACGCGCAGCGTGACCGCTACACTTGCCAGCGCCCTAGCGCCCGCTCCTTTCGCTTTCTTCCCTTCCTTTCTCGCCACGTTCGCCGGCTTTCCCCGTCAAGCTCTAAATCGGGGGCTCCCTTTAGGGTTCCGATTTAGTGCTTTACGGCACCTCGACCCCAAAAAACTTGATTAGGGTGATGGTTCACGTAGTGGGCCATCGCCCTGATAGACGGTTTTTCGCCCTTTGACGTTGGAGTCCACGTTCTTTAATAGTGGACTCTTGTTCCAAACTGGAACAACACTCAACCCTATCTCGGTCTATTCTTTTGATTTATAAGGGATTTTGCCGATTTCGGCCTATTGGTTAAAAAATGAGCTGATTTAACAAAAATTTAACGCGAATTTTAACAAAATATTAACGCTTACAATTTAGGTGGCACTTTTCGGGGAAATGTGCGCGGAACCCCTATTTGTTTATTTTTCTAAATACATTCAAATATGTATCCGCTCATGAGACAATAACCCTGATAAATGCTTCAATAATATTGAAAAAGGAAGAGTATGAGTATTCAACATTTCCGTGTCGCCCTTATTCCCTTTTTTGCGGCATTTTGCCTTCCTGTTTTTGCTCACCCAGAAACGCTGGTGAAAGTAAAAGATGCTGAAGATCAGTTGGGTGCACGAGTGGGTTACATCGAACTGGATCTCAACAGCGGTAAGATCCTTGAGAGTTTTCGCCCCGAAGAACGTTTTCCAATGATGAGCACTTTTAAAGTTCTGCTATGTGGCGCGGTATTATCCCGTATTGACGCCGGGCAAGAGCAACTCGGTCGCCGCATACACTATTCTCAGAATGACTTGGTTGAGTACTCACCAGTCACAGAAAAGCATCTTACGGATGGCATGACAGTAAGAGAATTATGCAGTGCTGCCATAACCATGAGTGATAACACTGCGGCCAACTTACTTCTGACAACGATCGGAGGACCGAAGGAGCTAACCGCTTTTTTGCACAACATGGGGGATCATGTAACTCGCCTTGATCGTTGGGAACCGGAGCTGAATGAAGCCATACCAAACGACGAGCGTGACACCACGATGCCTGTAGCAATGGCAACAACGTTGCGCAAACTATTAACTGGCGAACTACTTACTCTAGCTTCCCGGCAACAATTAATAGACTGGATGGAGGCGGATAAAGTTGCAGGACCACTTCTGCGCTCGGCCCTTCCGGCTGGCTGGTTTATTGCTGATAAATCTGGAGCCGGTGAGCGTGGGTCTCGCGGTATCATTGCAGCACTGGGGCCAGATGGTAAGCCCTCCCGTATCGTAGTTATCTACACGACGGGGAGTCAGGCAACTATGGATGAACGAAATAGACAGATCGCTGAGATAGGTGCCTCACTGATTAAGCATTGGTAACTGTCAGACCAAGTTTACTCATATATACTTTAGATTGATTTAAAACTTCATTTTTAATTTAAAAGGATCTAGGTGAAGATCCTTTTTGATAATCTCATGACCAAAATCCCTTAACGTGAGTTTTCGTTCCACTGAGCGTCAGACCCCGTAGAAAAGATCAAAGGATCTTCTTGAGATCCTTTTTTTCTGCGCGTAATCTGCTGCTTGCAAACAAAAAAACCACCGCTACCAGCGGTGGTTTGTTTGCCGGATCAAGAGCTACCAACTCTTTTTCCGAAGGTAACTGGCTTCAGCAGAGCGCAGATACCAAATACTGTTCTTCTAGTGTAGCCGTAGTTAGGCCACCACTTCAAGAACTCTGTAGCACCGCCTACATACCTCGCTCTGCTAATCCTGTTACCAGTGGCTGCTGCCAGTGGCGATAAGTCGTGTCTTACCGGGTTGGACTCAAGACGATAGTTACCGGATAAGGCGCAGCGGTCGGGCTGAACGGGGGGTTCGTGCACACAGCCCAGCTTGGAGCGAACGACCTACACCGAACTGAGATACCTACAGCGTGAGCTATGAGAAAGCGCCACGCTTCCCGAAGAGAGAAAGGCGGACAGGTATCCGGTAAGCGGCAGGGTCGGAACAGGAGAGCGCACGAGGGAGCTTCCAGGGGGAAACGCCTGGTATCTTTATAGTCCTGTCGGGTTTCGCCACCTCTGACTTGAGCGTCGATTTTTGTGATGCTCGTCAGGGGGGCGGAGCCTATGGAAAAACGCCAGCAACGCGGCCTTTTTACGGTTCCTGGCCTTTTGCTGGCCTTTTGCTCACATGTTCTTTCCTGCGTTATCCCCTGATTCTGTGGATAACCGTATTACCGCCTTTGAGTGAGCTGATACCGCTCGCCGCAGCCGAACGACCGAGCGCAGCGAGTCAGTGAGCGAGGAAGCGGAAGAGCGCCCAATACGCAAACCGCCTCTCCCCGCGCGTTGGCCGATTCATTAATGCAGCTGGCACGACAGGTTTCCCGACTGGAAAGCGGGCAGTGAGCGCAACGCAATTAATGTGAGTTAGCTCACTCATTAGGCACCCCAGGCTTTACACTTTATGCTTCCGGCTCGTATGTTGTGTGGAATTGTGAGCGGATAACAATTTCACACAGGAAACAGCTATGACCATGATTACGCCAAGCGCGCAATTAACCCTCACTAAAGGGAACAAAAGCTGGAGCTGCAAGCTT

**Supplementary Figure 4:** Fluorescent protein sequence alignment. Protein alignment of mRFP1, Dimer1, and DsRED-Express2 fluorescent proteins sequences downloaded from fpbase.org and generated using CLUSTAL 0 (1.2.4) through the EMBL-EBI online server (https://www.ebi.ac.uk/Tools/msa/clustalo/).

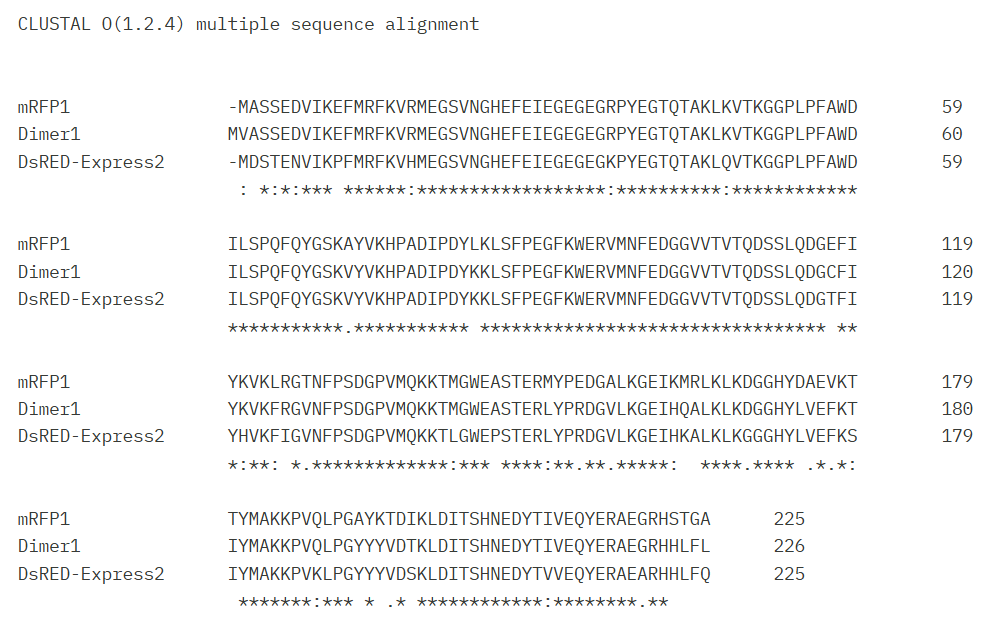

>mRFP1

MASSEDVIKEFMRFKVRMEGSVNGHEFEIEGEGEGRPYEGTQTAKLKVTKGGPLPFAWDILSPQFQYGSKAYVKHPADIPDYLKLSFPEGFKWERVMNFEDGGVVTVTQDSSLQDGEFIYKVKLRGTNFPSDGPVMQKKTMGWEASTERMYPEDGALKGEIKMRLKLKDGGHYDAEVKTTYMAKKPVQLPGAYKTDIKLDITSHNEDYTIVEQYERAEGRHSTGA

>Dimer1

MVASSEDVIKEFMRFKVRMEGSVNGHEFEIEGEGEGRPYEGTQTAKLKVTKGGPLPFAWDILSPQFQYGSKVYVKHPADIPDYKKLSFPEGFKWERVMNFEDGGVVTVTQDSSLQDGCFIYKVKFRGVNFPSDGPVMQKKTMGWEASTERLYPRDGVLKGEIHQALKLKDGGHYLVEFKTIYMAKKPVQLPGYYYVDTKLDITSHNEDYTIVEQYERAEGRHHLFL

>DsRED-Express2

MDSTENVIKPFMRFKVHMEGSVNGHEFEIEGEGEGKPYEGTQTAKLQVTKGGPLPFAWDILSPQFQYGSKVYVKHPADIPDYKKLSFPEGFKWERVMNFEDGGVVTVTQDSSLQDGTFIYHVKFIGVNFPSDGPVMQKKTLGWEPSTERLYPRDGVLKGEIHKALKLKGGGHYLVEFKSIYMAKKPVKLPGYYYVDSKLDITSHNEDYTVVEQYERAEARHHLFQ

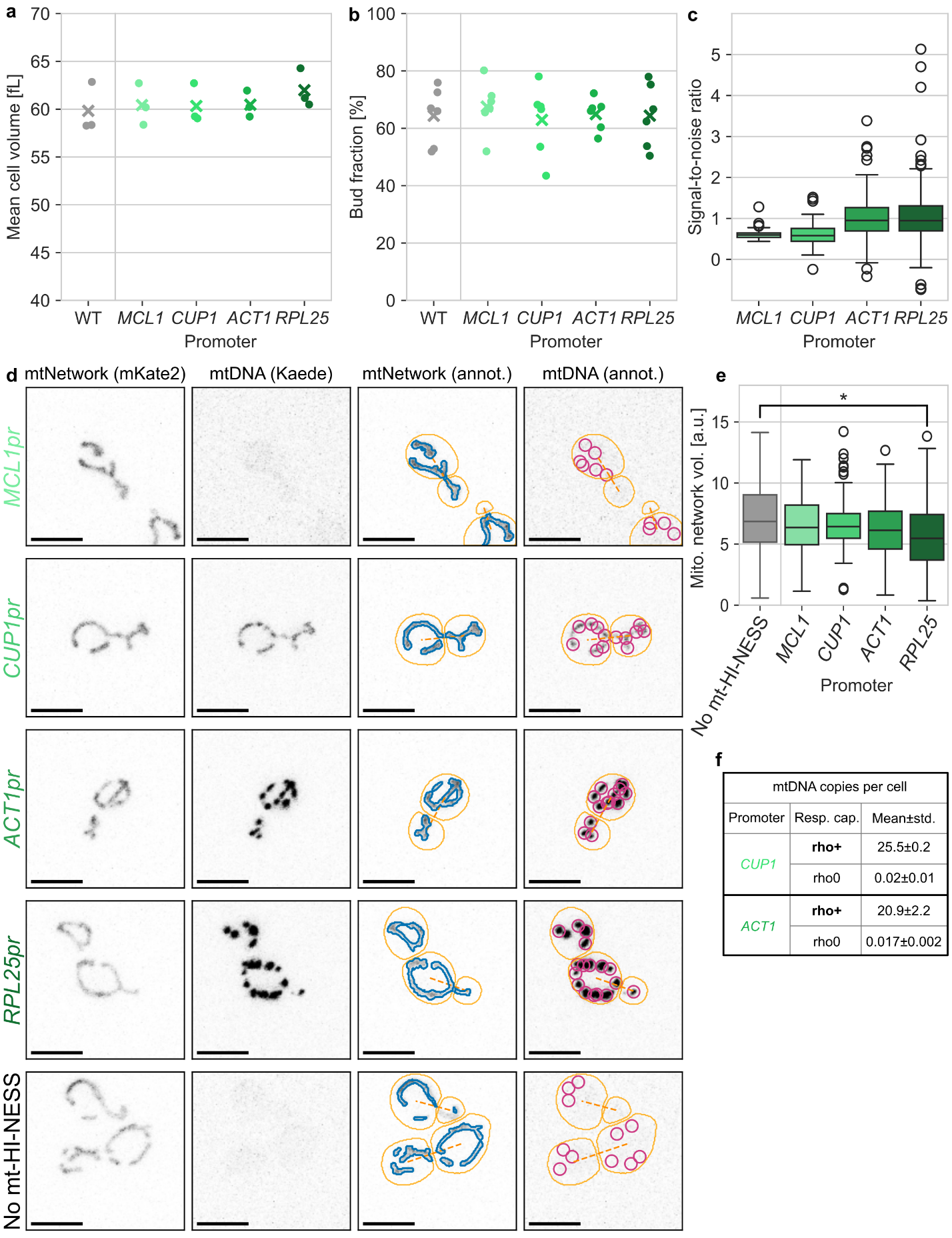

**Supplementary Figure 5:** A-B) Mean cell volumes as measured with a Coulter counter (A) and bud fractions (B) for wild-type cells (WT) and strains expressing the mt-Kaede-HI-NESS reporter from different promoters as indicated. C) Signal-to-noise ratios (see Methods) for automatically detected spots in the mt-Kaede-HI-NESS channel. D) Corresponding representative confocal images (maximum projections) as well as network segmentation masks and detected spots are shown for budding yeast expressing the mt-Kaede-HI-NESS reporter from different promoters as indicated, as well as for a no-reporter control. All strains express mKate2 targeted to the mitochondrial matrix for visualization of the mitochondrial network. Cell outlines and dashed lines connecting buds with their mother cells are shown in yellow. Automatic segmentation of the mitochondrial network is shown in blue. Automatically detected spots are shown in pink. Note that for comparability, the same microscopy settings were used for all strains. Scale bars denote 2 μm. E) Quantification of the mitochondrial network volume.F) mtDNA copies per cell as determined by qPCR for strains expressing mt-Kaede-HI-NESS from the CUP1 or ACT1 promoters, and corresponding mip1Δ cells.

| **MMY116-2C** | *Mat α, ADE2* | Skotheim lab stock |
| --- | --- | --- |
| **DBY079-2** | *Mat α, ADE2 HO:PGK1-SU9-mKATE2-KanMX4* | This study |
| **DBY080-2** | *Mat α, ADE2 ura3::URA3-CUP1pr-SU9-Linker-mt-Kaede-HI-NESS-CYC1term HO:PGK1-SU9-mKATE2-KanMX4* | This study |
| **DBY083-1** | *Mat α, ADE2 ura3::URA3-MLC1pr-SU9-Linker-mt-Kaede-HI-NESS-CYC1term HO:PGK1-SU9-mKATE2-KanMX4* | This study |
| **DBY084-1** | *Mat α, ADE2 ura3::URA3-ACT1pr-SU9-Linker-mt-Kaede-HI-NESS-CYC1term HO:PGK1-SU9-mKATE2-KanMX4* | This study |
| **DBY085-2** | *Mat α, ADE2 ura3::URA3-RPL25pr-SU9-Linker-mt-Kaede-HI-NESS-CYC1term HO:PGK1-SU9-mKATE2-KanMX4* | This study |
| **KSY346-1** | *Mat α, ADE2 ura3::URA3-CUP1pr-SU9-Linker-mt-Kaede-HI-NESS-CYC1term HO:PGK1-SU9-mKATE2-KanMX4* *mip1::CglaTRP1* | This study |
| **KSY347-1** | *Mat α, ADE2 ura3::URA3-ACT1pr-SU9-Linker-mt-Kaede-HI-NESS-CYC1term HO:PGK1-SU9-mKATE2-KanMX4* *mip1::CglaTRP1* | This study |

Supplementary Table 1: List of budding yeast strains used in this study. All strains were derived from W303.

| **Gene** | **Fwd/rev** | **Sequence** |
| --- | --- | --- |
| *COX2* | Fwd | GTTGATGCTACTCCTGGTAGATT |
| *COX2* | Rev | TTGCATGACCTGTCCCACAC |
| *COX3* | Fwd | TTGAAGCTGTACAACCTACC |
| *COX3* | Rev | CCTGCGATTAAGGCATGATG |
| *ACT1* | Fwd | AGTTGCCCCAGAAGAACACC |
| *ACT1* | Rev | GGACAAAACGGCTTGGATGG |
| *MRX6* | Fwd | CATCCGACGTGGTGCTCTTA |
| *MRX6* | Rev | TCTCATCTCTCCCTCCACCC |

Supplementary Table2: List of primers used for qPCR quantification of mtDNA copy number in budding yeast.

Supplementary Table 3: Antibodies used in Immunofluorescence and Immunoblotting

- Autoanti-dsDNA DSHB (<http://antibodyregistry.org/AB_10805293>) Used 1:200
- Anti-TOMM20 antibody [EPR15581-54] Used 1:1000
- Tom20 Antibody (F-10): #sc-17764 Used 1:1000
- HA-Tag (C29F4) Rabbit Monoclonal Antibody #3724 Used 1:1000
- HA-Tag Antibody (F-7): #sc-7392 Used 1:1000
- Thermo Fisher Goat anti-Rabbit 405 Cat # A-31556
- Thermo Fisher Goat anti-Rabbit 488 Cat # A-11034
- Thermo Fisher Goat anti-Rabbit 647 Cat # A-21245
- Thermo Fisher Donkey anti-Mouse 405 Plus Cat # A-48257
- Thermo Fisher Goat anti-Mouse 405 Cat # A-31553
- Thermo Fisher Goat anti-Mouse 488 Cat # A-11029
- Thermo Fisher Donkey anti-Mouse 647 Cat # A-31571

ImageJ Macro for mtDNA nucleoids analysis:

mainImage = getTitle();

roiManager("Deselect");

roiCount = roiManager("count");

// Set folder to save everything

saveDir = "Directory”

// Save all ROIs to .zip before processing

roiManager("Save", saveDir + "saved_ROIs.zip");

for (i = 0; i < roiCount; i++) {

selectWindow(mainImage);

roiManager("Select", i);

run("Duplicate...", "title=ROI_" + i);

selectWindow("ROI_" + i);

run("Clear Outside");

run("Auto Local Threshold", "method=Phansalkar radius=15 parameter_1=0 parameter_2=0 white");

run("Convert to Mask");

run("Analyze Particles...", "size=0.05-2.5 circularity=0.30-1.00 show=Overlay summarize exclude clear")

// Save binary mask image

saveAs("Tiff", saveDir + "ROI_" + i + "_mask.tif");

close();

run("Clear Results");

}

print("ROI set and mask images saved to: " + saveDir);
